# Mebendazole for Differentiation Therapy of Acute Myeloid Leukemia Identified by a Lineage Maturation Index

**DOI:** 10.1101/688192

**Authors:** Yulin Li, Daniel Thomas, Anja Deutzmann, Ravindra Majeti, Dean W. Felsher, David L. Dill

## Abstract

Accurate assessment of changes in cellular differentiation status in response to drug treatments or genetic perturbations is crucial for understanding tumorigenesis and developing novel therapeutics for human cancer. We have developed a novel computational approach, the Lineage Maturation Index (LMI), to define the changes in differentiation state of hematopoietic malignancies based on their gene expression profiles. We have confirmed that the LMI approach can detect known changes of differentiation state in both normal and malignant hematopoietic cells. To discover novel differentiation therapies, we applied this approach to analyze the gene expression profiles of HL-60 leukemia cells treated with a small molecule drug library. Among multiple drugs that significantly increased the LMIs, we identified mebendazole, an anti-helminthic clinically used for decades with no known significant toxicity. We tested the differentiation activity of mebendazole using primary leukemia blast cells isolated from human acute myeloid leukemia (AML) patients. We determined that treatment with mebendazole induces dramatic differentiation of leukemia blast cells as shown by cellular morphology and cell surface markers. Furthermore, mebendazole treatment significantly extended the survival of leukemia-bearing mice in a xenograft model. These findings suggest that mebendazole may be utilized as a low toxicity therapeutic for human acute myeloid leukemia and confirm the LMI approach as a robust tool for the discovery of novel differentiation therapies for cancer.

## Introduction

Hematopoietic system has a hierarchical arrangement. Through proliferation and differentiation, hematopoietic stem cells (HSCs) give rise to progenitor cells with intermediate maturation status, which further differentiate into mature blood cells^1-3^. A hallmark feature of AML leukemia cells is the blockade of differentiation at a distinct developmental stage. The differentiation block prevents leukemia cells from terminal differentiation, and the eventual apoptosis observed in the normal maturation of leukocytes, allowing leukemia cells to self-renew and proliferate^4,5^. Therefore, differentiation therapy that uses small molecules to specifically induce terminal differentiation could be an effective treatment without the toxicity of conventional chemotherapy^5,6^. The efficacy of differentiation therapy has been demonstrated by the successful treatment of acute promyelocytic leukemia (APL) with *all-trans* retinoic acid (ATRA) and arsenic oxide^7-9^.

ATRA, a differentiation agent discovered serendipitously, has been the standard therapy for APL leukemia for the past 30 years^8,10^. Although ATRA is highly efficient for leukemia harboring *PML-RAR*α gene fusions, ATRA does not show differentiation responses in non-APL leukemia. Recent development of epigenetic differentiation therapy for IDH1 mutated and IDH2 mutated AML with ivosidenib and enasidenib, respectively, has reinvigorated such approaches for other molecular subtypes of AML^11,12^. Large-scale drug response data repositories of cancer cells have been made available together with the associated gene expression profiles^13,14^. The availability of these data sets presents an opportunity for systematic identification of drugs that can induce differentiation in cancer cells, including leukemia cells. To exploit these data sets, we set out to develop a computational approach that could describe differentiation as a function of global gene expression changes.

Here, we have developed a robust computational method, called the Lineage Maturation Index (LMI), for assessing the changes in differentiation status of hematopoietic cells based on global gene expression profiles. To define the differentiation state of a specific cell (such as a leukemia cell), we project its gene expression profile onto a “reference lineage vector” that represents the physiological differentiation process from HSCs to the mature cells in the appropriate lineage. Upon drug treatment, a shift in the projection from stem cells towards mature cells indicates differentiation. We have validated that the LMI method can detect the differentiation of both normal hematopoietic populations and leukemia cells. We have used our LMI approach to analyze publicly available drug response data sets to identify drugs that induce leukemia cell differentiation. We have discovered multiple candidate drugs that induce LMI shifts. More importantly, we have demonstrated the therapeutic potential for our top candidate, mebendazole, which induces robust differentiation of primary human leukemia blasts *in vitro* and displays significant anti-leukemic activity in a xenograft model of non-APL leukemia *in vivo*.

## Results

### Development of the LMI approach

The genes that are differentially expressed between the HSCs and mature cells from a specific lineage can be identified by comparing the global gene expression profiles of the two cell populations. The expression profile of thousands of genes that change significantly can be considered as a single point in an *N*-dimensional space (Figure. 1A), where *N* is the number of differentially expressed genes. Based on the expression of these genes, a “reference lineage vector” can be derived from a point representing an HSC to a point representing a mature cell (such as a granulocyte). Differentiation state of a specific cell (such as leukemia cell) can be defined by projecting its expression profile onto this reference lineage vector. We define this projection as the Lineage Maturation Index (LMI) (Figure. 1B, also see Materials and Methods). If two cells have different LMIs, the cell with the larger LMI is more differentiated and has a point on the reference lineage vector that is closer to that of a mature cell. Since our primary interest is the discovery of drugs that can induce differentiation of leukemia cells, we have focused on *<u>changes</u>* in the LMIs induced by various chemical treatments (Figure. 1C).

**Figure. 1.**
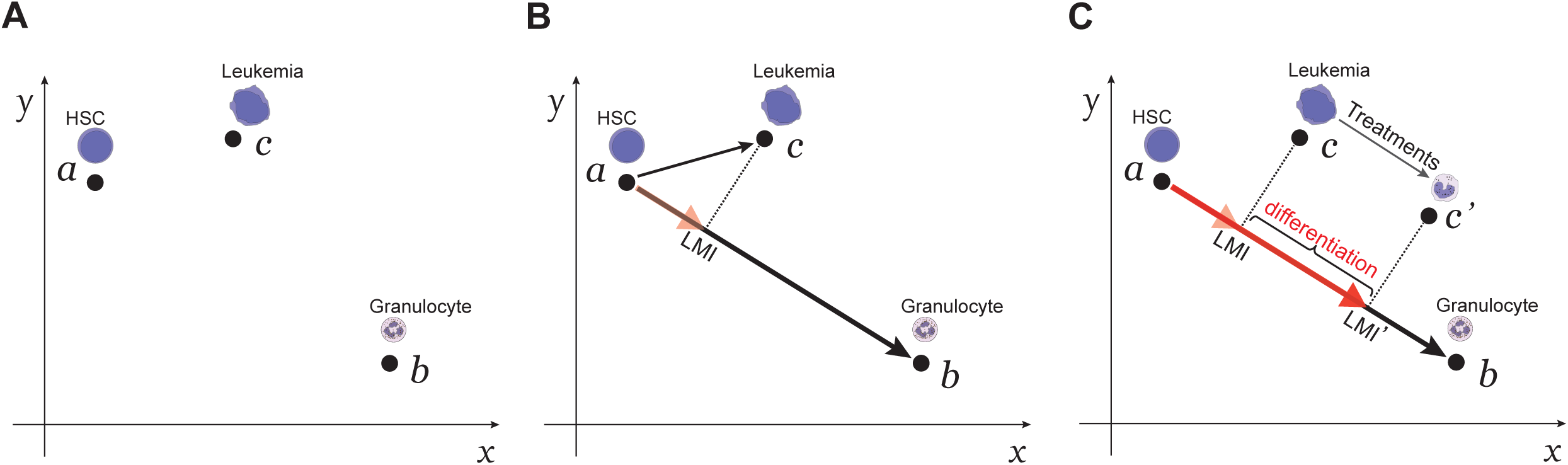
The concept of using the LMI approach to detect changes in differentiation states. (**A**) The gene expression profiles of the normal hematopoietic cells and AML leukemia cells can be represented by points (***a, b***, and ***c***) within an N-dimensional space. In this diagram, only two dimensions (*x* and *y*) are shown. (**B**) A myeloid-specific reference lineage vector is derived from a point representing the gene expression profile of the HSCs and a point representing that of mature granulocytes. The LMI for a specific myeloid cell (such as an AML leukemia cell) is the scalar projection (dashed line) of that cell type’s gene expression profile onto this lineage vector (red arrowhead). (**C**) Drug treatment of the AML leukemia cell can induce a shift in LMI (from LMI to LMI’). A shift of LMI towards the mature cells indicates differentiation.

### LMI can detect changes in differentiation status of both normal hematopoietic and leukemia cells

To examine whether the LMI method could reliably detect changes in differentiation status, we analyzed multiple data sets, including normal hematopoiesis, the classical model of ATRA differentiation therapy, and drugs known to modulate differentiation.

Within each hematopoietic lineage, committed progenitors undergo amplification and sequential maturation into peripheral blood cells. Based on the expression of surface markers, distinct intermediate populations have been isolated and functionally studied ^2^. We performed LMI analysis with several human hematopoietic lineages to evaluate whether the results accurately reflect the maturity of intermediate developmental stages along the lineage. In each case, the lineage vector was based on gene expression data for HSCs and the respective mature cells in the lineage, and we computed the LMIs of the intermediate cells in the same lineage from the same data set. In the analysis of human myeloid lineage from HSCs to granulocytes (HSC to GRAN3)^15^, the cell populations with intermediate maturation status showed monotonically increasing LMIs, corresponding to their previously established developmental sequence (Figure. 2A). Other lineages, such as erythroid, monocytic, and B lymphoid lineages, similarly showed a perfect match to the experimentally-derived developmental sequence (Figure. S1).

**Figure. 2.**
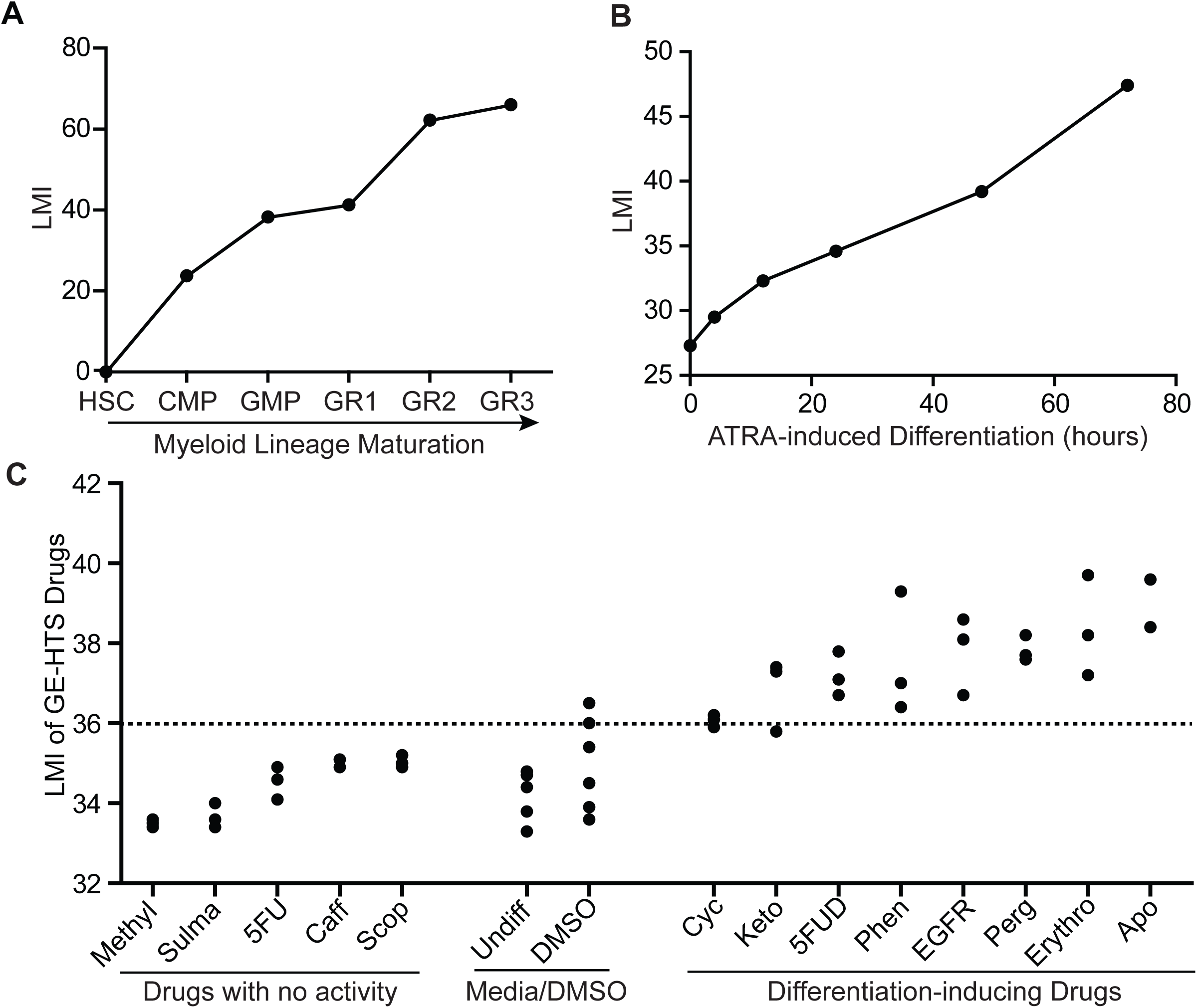
LMI detects changes in differentiation status in known examples of differentiation. (**A**) LMIs of distinct human myeloid cells along the developmental stages from the most immature stem cells to the mature granulocytes. (**B**) LMI analysis of APL leukemia (NB4 cells) treated with ATRA for 72 hours. (**C**) LMIs for the drugs from GE-HTS project. The t test for each drug treatment compared to DMSO controls is shown in Table S1.

We then tested whether LMI analysis could detect differentiation in the classical model of differentiation therapy, ATRA treatment of APL^7-9,16,17^. NB4 leukemia cell line was derived from an APL patient^18^. When treated with ATRA, NB4 cells differentiate along the myeloid lineage as shown previously by changes in cellular morphology and surface markers^19^. To test whether our method could computationally detect the *in vitro* differentiation process, we analyzed the LMIs of NB4 cells treated with ATRA in a 72-hour time course by projecting the gene expression profile vectors onto the normal human myeloid reference lineage vector. Notably, we observed an increase in LMI as early as 4 hours following ATRA treatment. From 4 hours to 72 hours after treatment, we saw a monotonic increase in LMI, which indicates progressive myeloid maturation (Figure. 2B). Therefore, LMI method can detect the maturation process of leukemic cells in the classical ATRA differentiation therapy.

Next, we determined whether our method could detect differentiation activities of chemicals that were previously reported^20^. The GE-HTS project has experimentally identified several chemicals that moderately induced the differentiation of HL-60 cells. We analyzed the data set from the GE-HTS project and computed the LMI for each drug treatment according to the myeloid reference lineage vector. Seven out of eight differentiation-inducing drugs from their screening significantly increased the LMIs compared with DMSO treatment. The other five drugs that were not found to have differentiation activity did not significantly affect the LMI (Figure. 2C, Table S1). This finding suggests that our method is highly sensitive and specific in identifying drugs that can induce differentiation. Thus, we have validated that the LMI method can assess the changes in differentiation states of both normal hematopoietic and leukemia cells. Furthermore, it can detect chemicals with known differentiation activities with high sensitivity and specificity.

### Identification of mebendazole as a differentiation agent by screening the HL-60 drug response C-MAP data sets

After confirming that our LMI approach is sensitive and specific in detecting changes in differentiation status, we set out to search for novel differentiation therapies by analyzing large-scale drug response data sets. To this end, we exploited the Connectivity Map (C-MAP), a catalog of microarray data collected from five human cancer cell lines treated with a drug library containing 1309 chemicals. Most of these drugs are already FDA-approved for clinical use^13,21^. One cell line included in the data set, HL-60, is a leukemia cell line derived from an AML patient ^22^. We analyzed 1235 drug treatment arrays for HL-60 leukemia cells and extracted the LMIs according to the human myeloid reference lineage in order to identify drugs that could induce a significant increase in LMI compared to DMSO controls (Figure. 3A, Table S2). Notably, five out of the 20 chemicals with the highest increases in LMIs were either ATRA or ATRA analogs (Figure. 3B-C, Table S2). This finding further supports that our method is very robust in identifying chemicals with differentiation activity. Ten unique drugs from the top 20 hits with the highest increases in LMI were further tested by *in vitro* nitroblue tetrazolium (NBT) reduction assay, to assess the differentiation of HL-60 cells into mature granulocytes. We also included ten randomly chosen drugs with low or no LMI shifts as control. After three days of treatment, several drugs with high LMI shifts, such as ATRA, mebendazole, etoposide, and dihydroergotamine, showed strong differentiation-inducing activities as measured by the positive NBT staining (Figure. 3C-D, Figure. S2A-B). In contrast, HL-60 cells treated with drugs that did not affect LMI did not show significant changes in NBT staining.

**Figure. 3.**
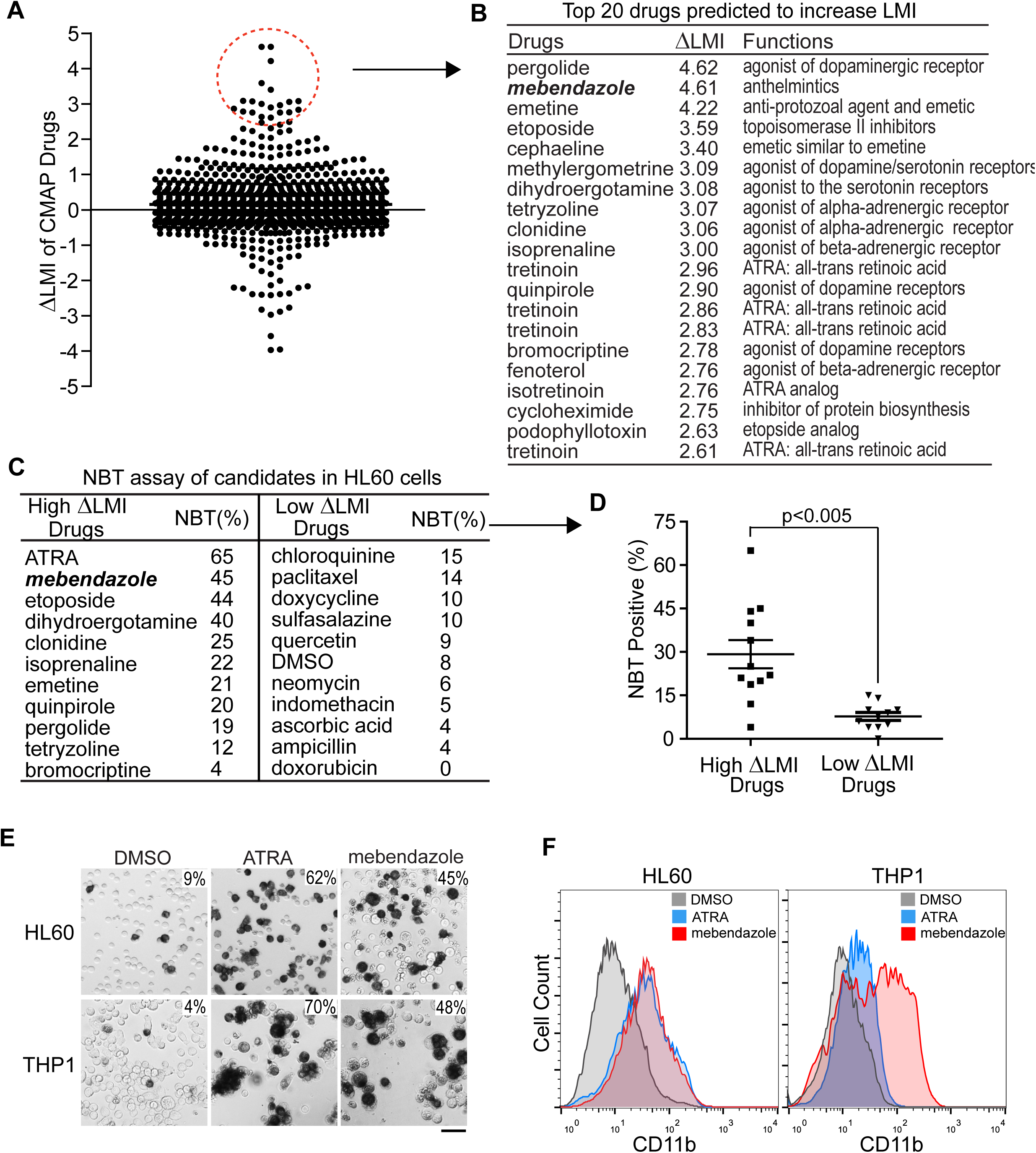
LMI analysis of HL-60 drug responses yields candidates with differentiation activity. (**A**) LMIs of 1235 arrays for HL-60 cells upon treatment with a drug library. ΔLMI for each drug is drug-induced LMI minus the LMI of the respective DMSO control. (**B**) Top 20 candidate drugs predicted to induce differentiation of HL-60 cells. (**C**) NBT assay of unique drugs from the top 20 candidates in HL-60 leukemia cells. The ten control drugs were randomly chosen from the drug library. All the drug treatments were carried out for three days. The NBT assay results were plotted in (**D**). Student’s t test with unequal variances, p<0.005. Mean+/-SEM is shown on the graph. (**E**) NBT staining of HL-60 and THP-1 leukemia cells treated with ATRA (1μM) and Mebendazole (0.33μM) for four days. The numbers at the top right corner indicate percent positive cells. Scale bar is 50 μm in length. (**F**) Flow cytometric analysis of CD11b expression in HL-60 and THP-1 leukemia cells treated for four days with ATRA (1μM) and mebendazole (0.33μM).

We further narrowed down the drug candidates by excluding multiple drugs with either systemic side effects (such as dihydroergotamine, clonidine, emetine, and isoprenaline) or myelosuppressive toxicity (such as etoposide). Our top candidate was mebendazole, an anti-helminthic drug used for decades with an excellent clinical safety profile^23^. Indeed, we observed significant differentiation activity for mebendazole in two AML leukemia cell lines (HL-60 and THP-1) by NBT staining and flow cytometric analysis of the myeloid marker CD11b expression (Figure. 3E-F). Thus, we focused our subsequent efforts on investigating the anti-leukemic activity of mebendazole.

### Preclinical testing of the anti-leukemic activities of mebendazole *in vitro* and *in vivo*

Next, we examined whether mebendazole has anti-leukemic activity in primary leukemia blasts from AML patients. We purified primary leukemia blasts from five human AML patients, including samples with *FLT3, NPM1* and *IDH1* mutations, and cultured them *ex vivo*^24^, followed by treatment with either DMSO or mebendazole (1μM) for a week. The changes in cellular differentiation status were investigated by examining cell surface marker expression with flow cytometry, and cellular morphology with Wright’s staining. In all five primary AML samples, mebendazole treatment induced significant differentiation in the leukemia blasts as evidenced by increased expression of myeloid markers (CD11b, CD11c, and CD14) (Figure. 4A, 4C, and Figure. S3A). We also examined cellular morphological changes in four out of five primary AML samples using Wright’s staining. Myeloid maturation was indeed induced after mebendazole treatment as shown by increased chromatin condensation, decreased nuclear-to-cytoplasmic ratio, reduction in cytoplasmic basophilia, and accumulation of granules in the cytoplasm (Figure. 4B-C, and Figure. S3B). Thus, mebendazole has potent differentiation activities *in vitro* in multiple primary human AML leukemias with diverse genetic abnormalities.

**Figure. 4.**
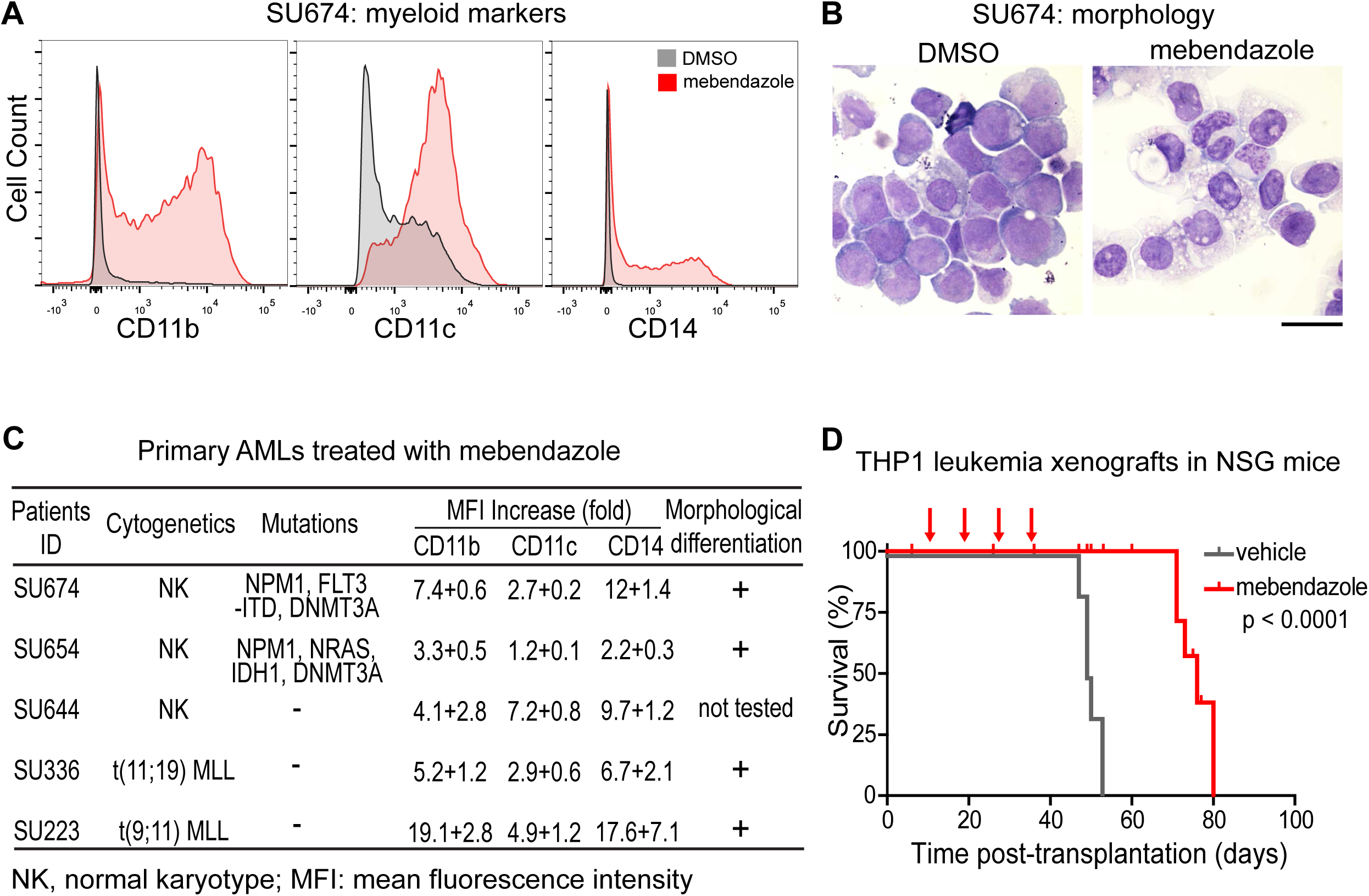
Mebendazole has anti-leukemic activity both *in vitro* and *in vivo*. (**A**) Expression of CD11b, CD11c, and CD14 in primary human leukemia sample (SU674) as shown by flow cytometric analysis. The leukemia blasts were treated with mebendazole (1μM) for seven days. (**B**) Morphology of primary human leukemia sample (SU674) as shown by Wright’s staining. The leukemia blasts were treated with mebendazole (1μM) for nine days. Scale bar is 20 μm in length. (**C**) Changes of surface markers and cell morphologies of five primary AML leukemia samples treated with mebendazole. Data are presented as mean+standard deviation. (**D**) Survival of NSG mice with THP-1 leukemia xenografts treated with either mebendazole (n=5, red line) or vehicle control (n=6, grey line).

We further evaluated the *in vivo* anti-leukemic activity of mebendazole by engrafting the highly aggressive human THP-1 leukemia cells in immunodeficient NOD/SCID/IL2Rγ null (NSG) mice. All leukemia-bearing NSG mice treated with vehicle control succumbed to the disease with a median survival of 49 days. In contrast, mebendazole as a single agent (50 mg/kg, 7 doses per cycle, 4 cycles) extended the median survival of the leukemia-bearing mice to 76 days without significant toxicity (Log-rank test, p<0.0001) (Figure. 4D). These results confirm mebendazole as a potent anti-leukemic agent both *in vitro* and *in vivo*.

## Discussion

To discover novel differentiation therapies, we have developed and validated a simple computational approach called the Lineage Maturation Index (LMI). By applying the LMI approach on C-MAP drug response data sets, we have identified mebendazole, a commonly used anthelmintic, to have a potent anti-leukemic activity against human leukemia *in vitro* and *in vivo*. The use of conventional anti-leukemic drugs, such as cytarabine, etoposide, and daunorubicin, is often limited due to their bone marrow toxicity. In contrast, long-term clinical use of high doses of mebendazole (100-200 mg/kg/day, 12-48 weeks) does not show serious side effects^23,25^. Sporadic clinical case reports have recently shown that mebendazole has therapeutic effects against adrenal and colon cancers^26,27^. There are ongoing clinical trials using mebendazole for treatment of brain and gastrointestinal cancers (NCT02644291, NCT01837862, and NCT03628079) based on preclinical studies^26,28–30^. Importantly, we showed that mebendazole induced differentiation responses in a range of non-APL AMLs including subtypes with mutations in *IDH1, FLT3* and *MLL* rearrangements. Our findings warrant a clinical trial of mebendazole for the treatment of human AML leukemia as monotherapy and potentially in combination with isocitrate dehydrogenase inhibitors also known to induce differentiation phenotype. AML is mainly a disease of the elderly, who have more comorbidity and are less tolerant to the intensive chemotherapy. Due to its potent anti-leukemic activity and low toxicity, mebendazole is particularly suitable for the treatment of elderly leukemia patients^31–33^.

As an anti-helminthic, mebendazole binds tightly to parasite tubulin and blocks its polymerization. In contrast, mebendazole only weakly interacts with mammalian tubulin, with an inhibition constant that is 250-400 times higher^34^. It is therefore unlikely that plasma levels of mebendazole are sufficient to block mammalian tubulin *in vivo*. Thus, the mechanism of action of mebendazole on various cancers *in vitro* and *in vivo* may be independent of its interaction with tubulin. Mebendazole has been recently reported to suppress multiple oncogenic targets in cancer cells, including hedgehog, VEGFR2, BRAF, BCR-ABL, and MYB^35–38^. In particular, the expression of MYB oncogene was downregulated following mebendazole treatment in multiple AML cell lines^38^. These and other targets may provide a potential explanation for the differentiation phenotype observed in our study. Of note, LMI also predicted etoposide and several dopamine receptor agonists including bromocriptine to induce a differentiation response in HL-60 cells, which we confirmed experimentally. Bromocriptine has recently been shown to have selective cytotoxicity in myelodysplasia and secondary AML^23^.

Several approaches have been taken towards the discovery of novel differentiation agents. Screening of small molecule compound libraries using reporter assays and/or high-content imaging is highly effective but labor intensive^39,40^. The development of genome-wide library screening approaches using CRISPR technology in particular, may be more feasible to identify genetic targets of differentiation therapy^41^. Our computational approach utilizes existing drug response data sets and is well-suited to detect changes in differentiation state of both normal hematopoietic and leukemia cells. This method may also be used to assess the changes in differentiation status and identify novel differentiation agents for solid tumors, provided a consistent hierarchy of maturation can be defined in the corresponding normal tissue of origin.

## Methods

### Algorithm

LMI is computed for a specific hematopoietic cell by comparing its gene expression profile to the gene expression profiles of normal immature and mature cells in a hematopoietic lineage. Only genes with at least a 4-fold change (difference of two in log_2_ transformed data) in expression in the normalized microarray expression data were used. In the following equation, ***a*** and ***b*** are vectors of gene expression values from an immature cell type, such as hematopoietic stem cell (HSC), and a mature cell, such as granulocyte. If ***c*** is the gene expression vector of a hematopoietic cell sample, the LMI is the scalar projection of ***c***−***a*** onto the vector ***b***− ***a*** (see Figure. 1):

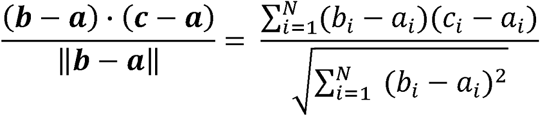

### Data processing and normalization

Mouse and Human Affymetrix gene expression arrays were used. We randomly selected 500 Mouse 430 2.0 arrays from GEO. fRMA (“frozen RMA”)^42,43^ vectors were computed using the makeVectorsAffyBatch command with 5 replicates of 100 arrays each, using the BrainArray version 17.1.0 Mouse 4302 ENSG alternative CDF file. Human arrays were a mixture of different platforms from the U133 family: U133A, U133 plus 2.0, U133A2, and U133AAofAv2 arrays. These arrays have varying numbers of probes, but almost all of the U133A probe sets appear on all of the other U133 array types, and each probe set has the same list of probes on each array. We converted the CEL files for all of the arrays to U133A array CEL files by assigning probe intensities to their corresponding positions on the U133A, based on probe set names. The few probes on the U133A array that did not appear on all of the other arrays were assigned the same values in all of the converted arrays. Thousands of arrays were downloaded from GEO and converted, and 500 of these were randomly chosen to reduce computation. fRMA vectors were then computed as with the mouse arrays, using BrainArray version 17.1.0 HGU133A ENSG CDF file.

### Microarray data sets

Normal hematopoiesis cell types: 211 arrays for 38 types of purified normal human hematopoietic cells (Broad Institute “Differentiation map portal”; GSE24759). We used average expression for all replicates of each cell type in each data set, except that we averaged HSC1 and HSC2 cell types in the Differentiation Map to obtain the gene expression signature for human HSCs.

In the Result section, we analyzed the following data sets; dynamic differentiation of normal hematopoietic cells: 22 arrays for the differentiation of human erythroid lineages (GSE36984); APL differentiation therapy using ATRA: 6 arrays for ATRA treated human NB4 cells (GSE19201 and GSE19203); GE-HTS analysis: 75 arrays for HL-60 cells treated with 13 drugs from the GE-HTS project (GSE982, HL-60 cells treated with 13 drugs); Connectivity Map screening: 1235 arrays from the Connectivity Map.

http://www.broadinstitute.org/scientific-community/science/projects/connectivity-map/connectivity-map.

To make the LMI values independent of the data sets being assessed, each individual array was normalized with fRMA, using the vectors computed from random arrays as described above.

### C-MAP data processing

C-MAP arrays were converted to the U133A format as described above. C-MAP arrays are organized in batches of two sizes, some of which have one control sample and some of which have several control samples. We computed LMIs separately for batches of control arrays, using the mean expression levels when there were several controls for a batch. LMIs were computed for each drug-treated sample. The change in LMI for each treated sample was the LMI of the treated sample minus the LMI for the controls from that batch.

### NBT assay

Differentiation of HL-60 and THP-1 cells was visualized by a nitro-blue tetrazolium (NBT) reduction assay. The colorless NBT can be reduced to the insoluble blue formazan within the differentiated cells. Cells were seeded onto T25 flasks at approximately 0.5 million/ml. Tested drugs were dissolved in either water or DMSO and the final concentration of DMSO in the culture media remained below 0.05%. Drug treatments were performed in triplicates for three days. Following treatment with various drugs, cells were incubated at 37°C for 60 minutes with 1 mg/ml NBT, and 5 µg/mL TPA (12-O-tetradecanoylphorbol-13-acetate). The percentage of dark blue cells was counted by light microscopy for at least 200 cells per sample.

### Flow cytometric analysis

Following a 4-day drug treatment, approximately 1-2 million HL-60 and THP-1 cells were resuspended in PBS containing 1% fetal bovine serum and stained with PE anti-human CD11b antibody (Biolegend). Stained cells were analyzed on FACScalibur flow cytometer (Becton Dickinson). Data were analyzed with FlowJo software (Tree Star). Live cells were gated based on forward and side scatter.

### Mebendazole treatment and differentiation assay for primary human AMLs

Patients’ AML blasts were sorted for live propidium-iodide negative cells on BD FACS ARIA II after thawing in IMDM 20% serum with DNase (2000u/ml). Cells were grown in triplicates at 10^6^/ml in Myelocult H5100 (Stem Cell Technologies) with (20ng/ml) IL-3, SCF, FLT3L, TPO, G-CSF and GM-CSF (Peprotech) for up to 6 days in the presence of mebendazole or DMSO. Differentiation was assessed by flow cytometry using anti-human CD11b-PE-Cy7 (ICRF44), CD11c-PAC-Blue (B-ly6), CD14-APC-Cy7 (MLP9), CD15-FITC (HI98) and CD33-APC (WM53), CD117-PE (YB5.B8) (all from BD Biosciences) and compared to isotype controls and FMO stain. Data were analyzed with FlowJo software (Tree Star). Live cells were gated based on propidium iodide staining.

### Wright’s staining of primary AMLs treated with mebendazole

Following mebendazole treatment, cells were stained with modified Wright Stain (Sigma). Briefly, cells were first stained with 0.8ml of Wright Stain for 3-4 minutes. Equal volume of deionized water was then added to the Wright Stain on the slides and staining continued for another 6-10 minutes. Slides were washed in tap water and air-dried before examination under a DMI 6000 microscope (Leica).

### Mebendazole treatment of mice with leukemia xenografts

Six million THP-1 human leukemia cells were transplanted into 16 NSG mice via tail vein injection. Five days after injection, mebendazole (50mg/kg, dissolved in soybean oil, oral gavage) or vehicle (soybean oil) was administered daily for seven days followed by two days of rest. The treatment cycle was repeated three more times for a total of 28 doses. Overall survival of each group was compared using Kaplan-Meier analysis.

### Animals and human patients’ samples

Animal studies, procurement of patients’ samples, and experimental protocols were approved by Stanford University’s Institutional Animal Care and Use Committee (IACUC) and Institutional Review Board (IRB). All experimental methods were performed in accordance with the relevant national and as well as Stanford University’s guidelines and regulations. Primary bone marrow and peripheral blood AML samples were obtained prior to treatment with informed consent from all patients according to institutional guidelines (Stanford University IRB No. 6453). The animal studies using NSG mice were approved by the Stanford University IACUC committee under approval number APLAC14045.

## Acknowledgements

We thank the researchers who have deposited their microarray data in Gene Expression Omnibus. We also acknowledge the Stanford Hematology Division Tissue Bank and the patients for donating their samples.

## Author contributions

Y.L. and D.L.D. designed the study, developed the approach, and analyzed the data. D.T. and Y.L. carried out the *in vitro* and *in vivo* studies using primary human leukemia blasts. Y.L. and A.D. performed the *in vitro* drug screening and *in vivo* animal therapeutic study. R.M. and D.W.F. assisted in the experimental design. Y.L. and D.L.D. wrote the paper.

## Data Availability

All data generated or analyzed during this study are included in this article and its Supplementary Information files.

## Supplementary Figure Legends

**Figure. S1.** LMI can differentiate among various intermediate populations within normal human hematopoietic system. (**A**) human erythroid lineage. (**B**) human B lymphocyte lineage. (**C**) human monocytic lineage.

**Figure. S2.** NBT staining of HL-60 leukemia cells treated with C-MAP drugs. (**A**) Treatments with drugs that have high LMI increases. (**B**) Treatments with drugs that are randomly chosen from the C-MAP drug library. Scale bar is 50 μm in length.

**Figure. S3.** Differentiation upon mebendazole treatment in primary AML blasts from patients. (**A**) Flow cytometric analysis of CD11b, CD11c, and CD14 expression in primary AML samples treated with mebendazole. (**B**) Morphological changes of primary AML leukemic cells treated with mebendazole as shown by Wright’s staining. Scale bar is 20 μm in length.

