## Supplementary figures and images for "Mebendazole for Differentiation Therapy of Acute Myeloid Leukemia Identified by a Lineage Maturation Index"

### Supplementary Fig S1-S3

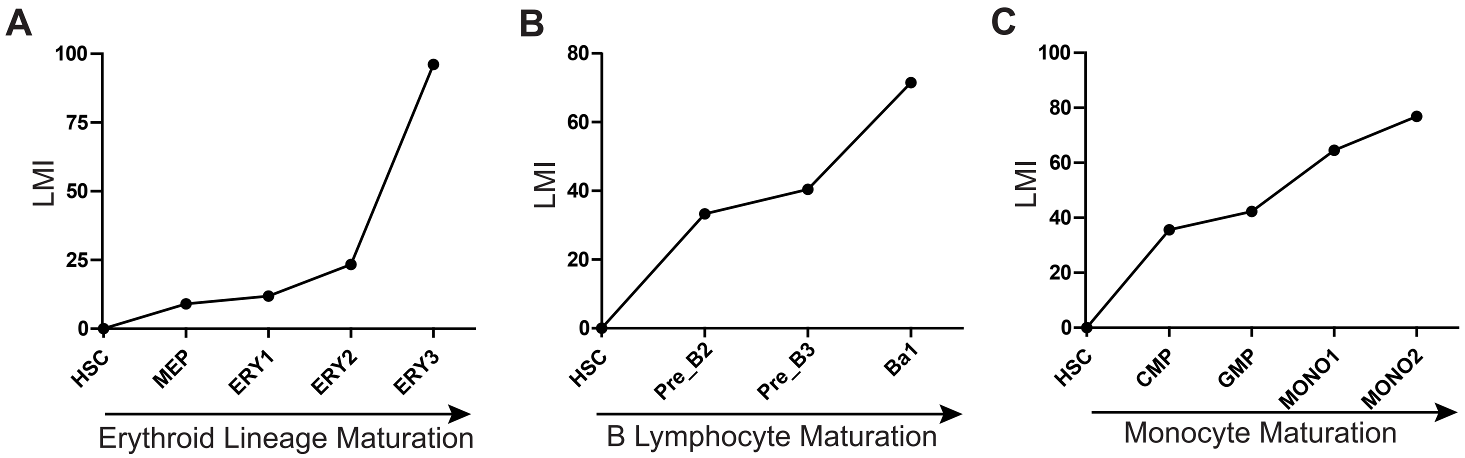

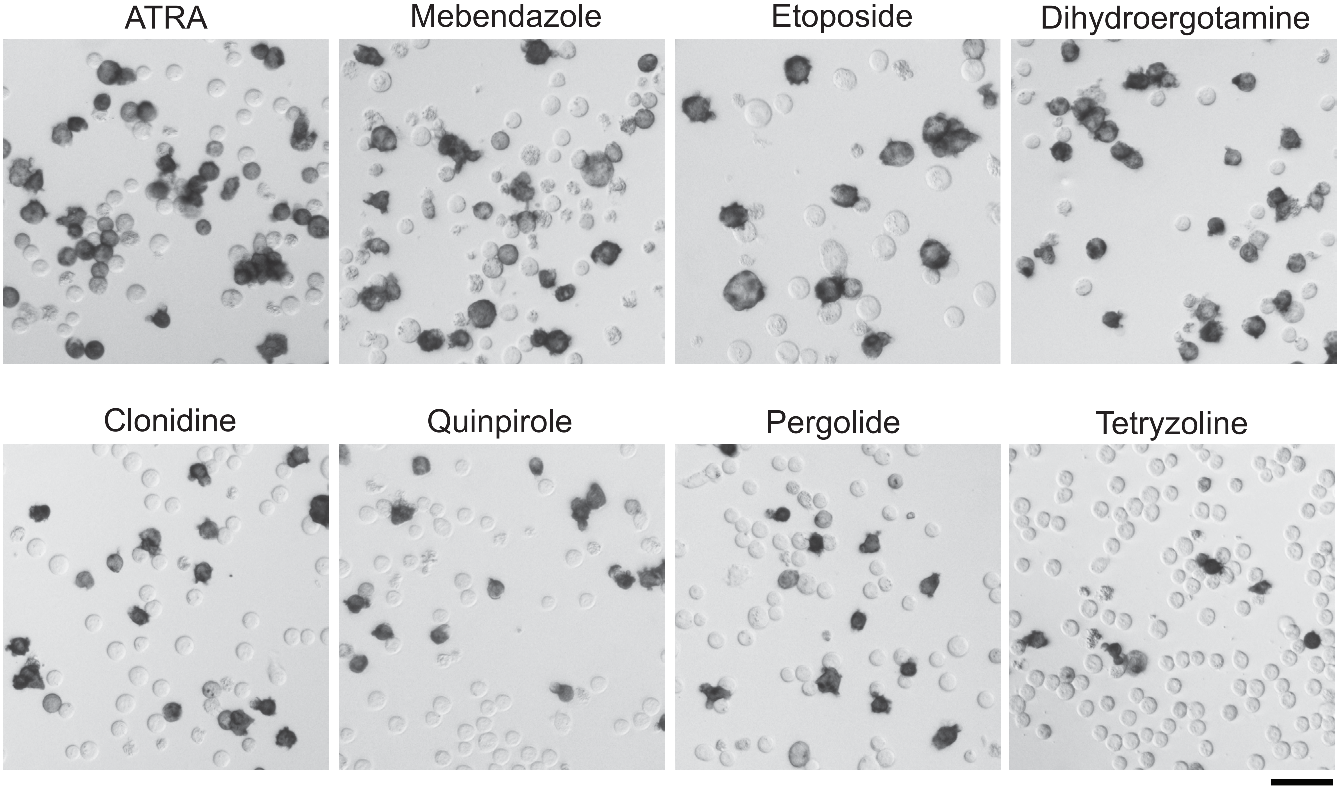

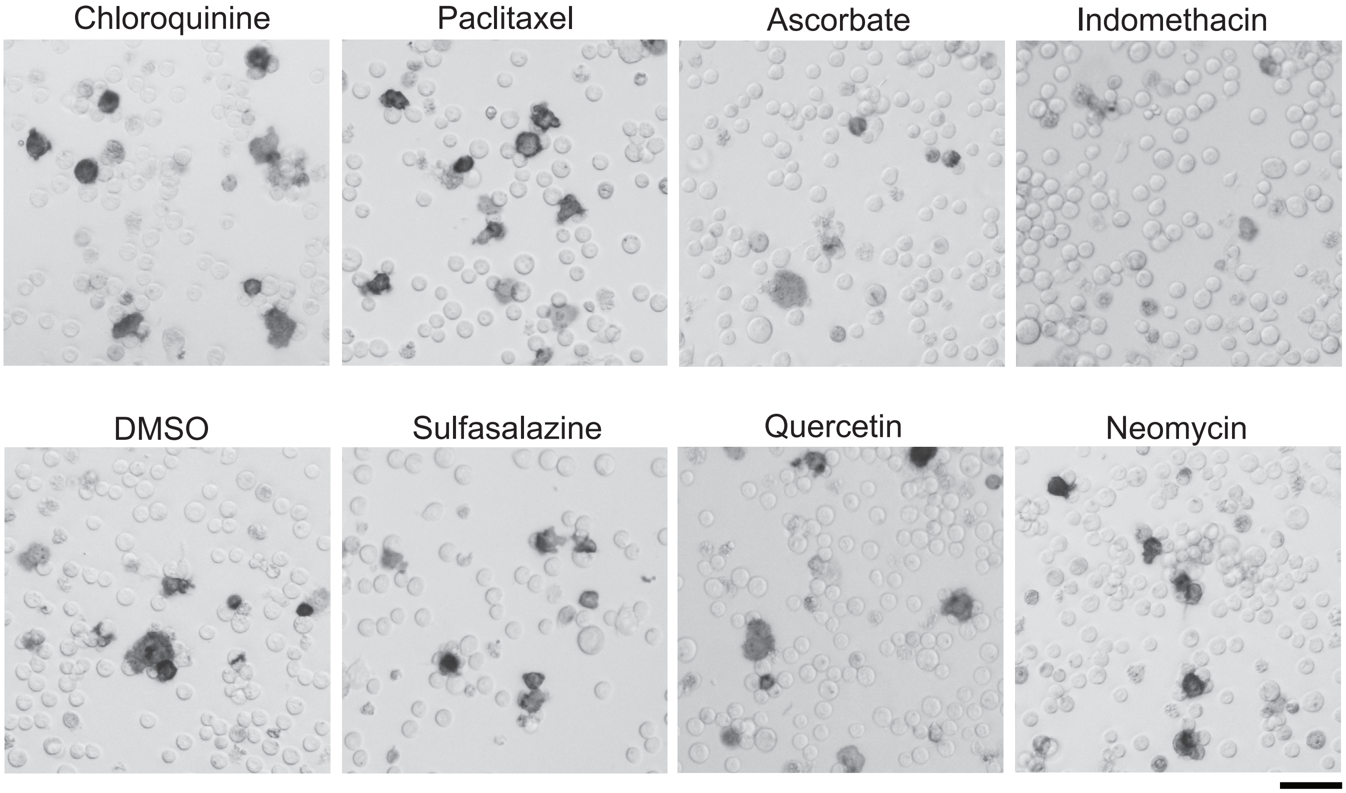

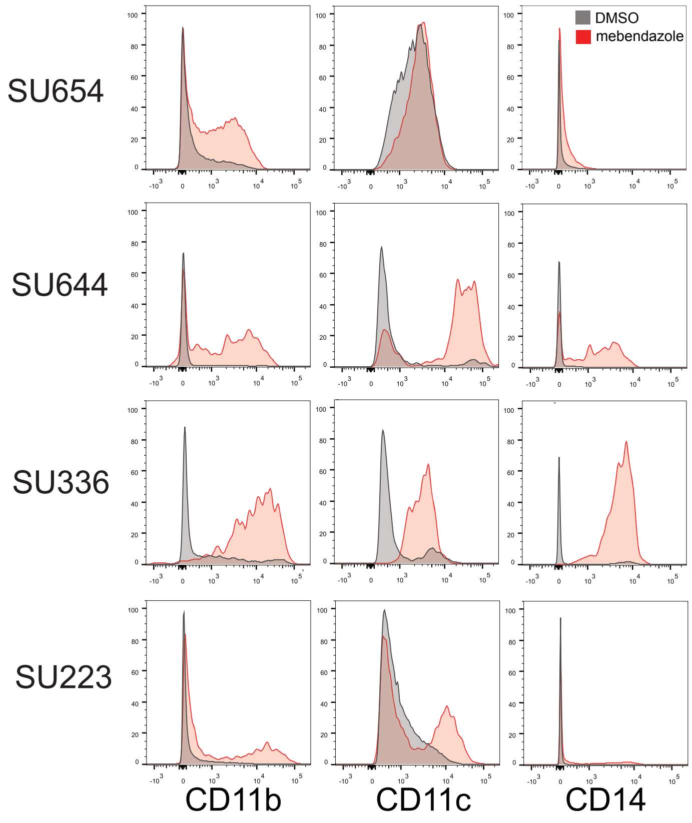

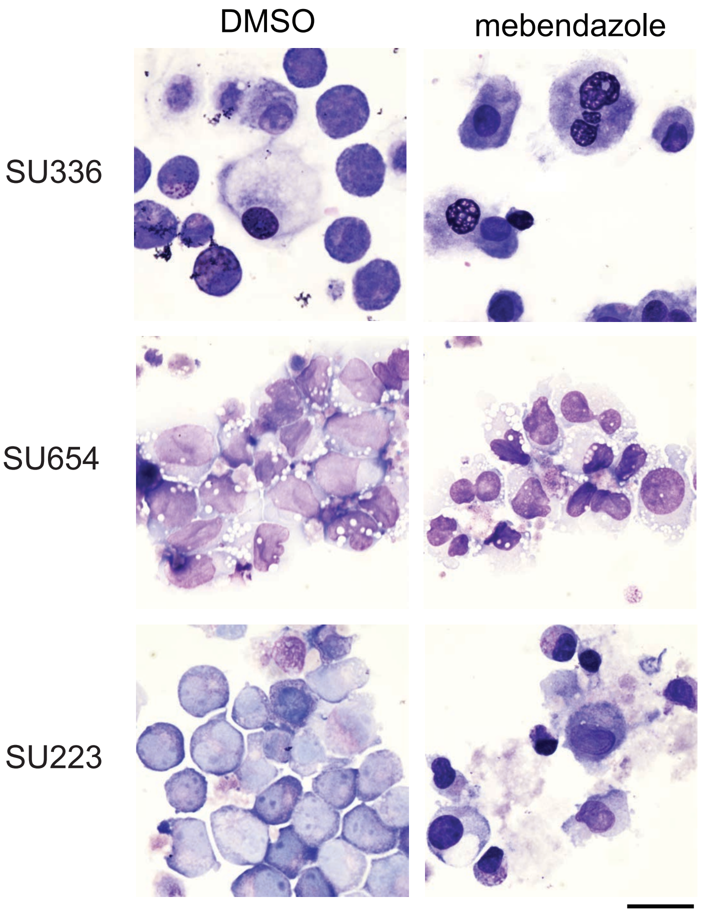
