## Supplementary Tables 1-2 for "Mebendazole for Differentiation Therapy of Acute Myeloid Leukemia Identified by a Lineage Maturation Index"

**Table S1: LMIs of the drugs from GE-HTS project.**

| Treatment | Array 1 | Array 2 | Array 3 | Array 4 | Array 5 | Array 6 | P value |
|---|---|---|---|---|---|---|---|
| Undiff | 33.8 | 33.8 | 33.3 | 34.4 | 34.8 | 34.7 | 0.143 |
| DMSO | 33.6 | 33.9 | 34.5 | 36.5 | 36 | 35.4 | 1.000 |
| Erythro | 37.2 | 38.2 | 39.7 | | | | 0.005 |
| Methyl | 33.6 | 33.4 | 33.5 | | | | 0.072 |
| Sulma | 33.4 | 34 | 33.6 | | | | 0.105 |
| Keto | 37.3 | 37.4 | 35.8 | | | | 0.049 |
| EGFR | 38.6 | 38.1 | 36.7 | | | | 0.009 |
| Phen | 36.4 | 37 | 39.3 | | | | 0.025 |
| 5FUD | 37.8 | 36.7 | 37.1 | | | | 0.019 |
| Perg | 38.2 | 37.7 | 37.6 | | | | 0.005 |
| Scop | 35.2 | 35 | 34.9 | | | | 0.945 |
| 5FU | 34.1 | 34.6 | 34.9 | | | | 0.549 |
| Caff | 35.1 | 34.9 | 34.9 | | | | 0.982 |
| Apo | 38.4 | 39.6 | 38.4 | | | | 0.001 |
| Cyc | 36.1 | 36.2 | 35.9 | | | | 0.166 |
| VitD | 46.7 | 47.3 | 47.3 | | | | 0.000 |
| PMA | 40.8 | 39.9 | 40.2 | | | | 0.000 |
| ATRA | 39.2 | 40.3 | 39.5 | | | | 0.000 |

**Note:**
All treaments were compared to DMSO using Student's t test.
Results of the t tests were plotted in Figure 2C.
VitD, PMA and ATRA were the positive controls.
Drugs reported to have differentiation activity in the GE-HTS project were highlighted in red.
Cyc (cyclazosin) treatment did not show a statistically significant increase in LMIs.

**Table S2. LMI of 1235 arrays for the HL-60 C-MAP drug treatments**

| Drug Name | LMIs of Drugs | LMIs of DMSO conti | LMI Shifts | perturbation_scan_id | vehicle_scan_id4 |
|---|---|---|---|---|---|
| pergolide | 37.61878802 | 33.00109826 | 4.617689757 | 5500024024213121906560.B | .H07.G08.E09.D10.B11.A12 |
| mebendazole | 38.07285774 | 33.45935203 | 4.613505706 | 5500024024213121906560.G | .H01.G02.E03.D04.B05.A06 |
| emetine | 39.23596421 | 35.01997127 | 4.215992936 | 5.50E+25 | .H01.G02.E03.D04.B05.A06 |
| etoposide | 37.04604708 | 33.45797002 | 3.58807706 | 622623112706.A10 | .H07.G08.E09.D10.B11.A12 |
| cephaeline | 38.55816628 | 35.15552717 | 3.402639109 | 5500024024214122006604.G | .H01.G02.E03.D04.A06 |
| methylergometrine | 36.54416434 | 33.45797002 | 3.086194328 | 622623112706.D07 | .H07.G08.E09.D10.B11.A12 |
| dihydroergotamine | 37.82977533 | 34.74789333 | 3.081882002 | 614615111406.B04 | .H01.G02.E03.D04.B05.A06 |
| tetryzoline | 37.03346312 | 33.96076648 | 3.072696644 | 5500024025833041107269.H | .H01.G02.E03.D04.B05.A06 |
| clonidine | 36.50844166 | 33.4530122 | 3.055429454 | 622623112706.G06 | .H01.G02.E03.B05.A06 |
| (-)-isoprenaline | 40.24499304 | 37.24787327 | 2.997119765 | 5500024035736031208612.C | .H01.G02.E03.D04.B05.A06 |
| tretinoin | 40.27693543 | 37.31290545 | 2.964029982 | 5500024035736031208612.G | .G08.E09.D10.B11.A12 |
| quinpirole | 37.07850086 | 34.17633596 | 2.9021649 | 5500024030760072207033.A | .H07.G08.E09.B11.A12 |
| tretinoin | 39.00594942 | 36.14209273 | 2.863856694 | 5500024017802120306174.G | .H01.G02.E03.D04.B05.A06 |
| tretinoin | 37.25512615 | 34.42118584 | 2.833940317 | 5500024030401071707292.G | .H01.G02.E03.D04.B05.A06 |
| bromocriptine | 38.21125473 | 35.4281341 | 2.783120629 | 612613111306.H10 | .H07.G08.E09.D10.B11.A12 |
| fenoterol | 35.76007621 | 33.00109826 | 2.758977951 | 5500024024213121906560.G | .H07.G08.E09.D10.B11.A12 |
| isotretinoin | 35.75633787 | 33.00109826 | 2.755239602 | 5500024024213121906560.B | .H07.G08.E09.D10.B11.A12 |
| cicloheximide | 37.15489975 | 34.40088765 | 2.754012101 | 5.50E+33 | .H07.G08.E09.D10.B11 |
| podophyllotoxin | 36.59543193 | 33.96076648 | 2.634665452 | 5500024025833041107269.A | .H01.G02.E03.D04.B05.A06 |
| co-dergocrine mesilat | 37.63427517 | 35.01997127 | 2.6143039 | 5500024024213121906559.G | .H01.G02.E03.D04.B05.A06 |
| tretinoin | 35.93951186 | 33.4530122 | 2.486499662 | 622623112706.H04 | .H01.G02.E03.B05.A06 |
| menadione | 37.62249204 | 35.15552717 | 2.466964871 | 5.50E+25 | .H01.G02.E03.D04.A06 |
| alprostadil | 37.12019437 | 34.70962788 | 2.410566484 | 5500024030760072207033.A | .H01.G02.E03.D04.B05.A06 |
| corbadrine | 36.70730928 | 34.40088765 | 2.306421632 | 5500024030401071707292.G | .H07.G08.E09.D10.B11 |
| picrotoxinin | 37.3181547 | 35.01997127 | 2.298183431 | 5500024024213121906559.B | .H01.G02.E03.D04.B05.A06 |
| mephentermine | 36.2033931 | 33.96129529 | 2.242097807 | 5500024025833041107269.D | .H07.E09.D10.B11.A12 |
| salbutamol | 35.68992145 | 33.45935203 | 2.230569424 | 5500024024213121906560.F | .H01.G02.E03.D04.B05.A06 |
| niclosamide | 37.47162574 | 35.2669347 | 2.204691037 | 612613111306.B04 | .H01.G02.E03.D04.B05.A06 |
| isoetarine | 36.47392415 | 34.40088765 | 2.073036503 | 5500024030401071707292.G | .H07.G08.E09.D10.B11 |
| medrysone | 36.00088244 | 33.96129529 | 2.039587153 | 5500024025833041107269.H | .H07.E09.D10.B11.A12 |
| clenbuterol | 35.48204703 | 33.45797002 | 2.024077013 | 622623112706.C08 | .H07.G08.E09.D10.B11.A12 |
| (+)-isoprenaline | 36.23280246 | 34.21031976 | 2.022482701 | 5500024030760072207028.D | .H07.G08.E09.D10.B11.A12 |
| metoprolol | 35.90485872 | 33.96076648 | 1.944092245 | 5500024025833041107269.A | .H01.G02.E03.D04.B05.A06 |
| thioridazine | 38.06162041 | 36.14209273 | 1.919527677 | 5500024017802120306174.C | .H01.G02.E03.D04.B05.A06 |
| pivampicillin | 36.08538825 | 34.17633596 | 1.909052296 | 5500024030760072207033.H | .H07.G08.E09.B11.A12 |
| iopanoic acid | 36.03645394 | 34.17633596 | 1.860117984 | 5500024030760072207033.D | .H07.G08.E09.B11.A12 |
| nimodipine | 39.03814807 | 37.24471367 | 1.793434395 | 5500024030700072107987.A | .H01.G02.E03.D04.B05.A06 |
| colchicine | 35.24285004 | 33.45797002 | 1.784880028 | 622623112706.F09 | .H07.G08.E09.D10.B11.A12 |
| nocodazole | 36.5178268 | 34.74789333 | 1.769933477 | 614615111406.C05 | .H01.G02.E03.D04.B05.A06 |
| pyrvinium | 35.94053312 | 34.17633596 | 1.764197167 | 5500024030760072207033.F | .H07.G08.E09.B11.A12 |
| dinoprost | 36.89645834 | 35.15552717 | 1.740931174 | 5500024024214122006604.C | .H01.G02.E03.D04.A06 |
| levonorgestrel | 35.70091065 | 33.96129529 | 1.739615358 | 5500024025833041107269.H | .H07.E09.D10.B11.A12 |
| trichostatin A | 36.67804214 | 34.94387722 | 1.734164918 | EC2004030506AA | EC2004030502AA |
| Prestwick-983 | 39.16936879 | 37.46016387 | 1.709204926 | 5500024030700072107987.B | .H07.G08.E09.D10.B11.A12 |
| ipratropium bromide | 36.76906708 | 35.08636713 | 1.682699952 | 6.41E+22 | .H07.G08.E09.D10.B11.A12 |
| puromycin | 36.82046358 | 35.15552717 | 1.664936412 | 5500024024214122006604.C | .H01.G02.E03.D04.A06 |
| thioridazine | 38.94278027 | 37.31290545 | 1.629874814 | 5500024035736031208612.C | .G08.E09.D10.B11.A12 |
| alpha-ergocryptine | 35.5751487 | 33.96129529 | 1.613853414 | 5500024025833041107269.C | .H07.E09.D10.B11.A12 |
| zardaverine | 36.30579708 | 34.70962788 | 1.596169204 | 5500024030760072207033.C | .H01.G02.E03.D04.B05.A06 |
| orciprenaline | 36.72274614 | 35.13162255 | 1.591123588 | 5500024024214122006604.D | .H07.G08.E09.D10.B11.A12 |
| vigabatrin | 36.7285016 | 35.15552717 | 1.57297443 | 5500024024214122006604.B | .H01.G02.E03.D04.A06 |
| genistein | 37.6908361 | 36.14209273 | 1.548743373 | 5500024017802120306174.B | .H01.G02.E03.D04.B05.A06 |
| labetalol | 34.98443078 | 33.4530122 | 1.531418579 | 622623112706.H06 | .H01.G02.E03.B05.A06 |
| nadide | 35.44293943 | 33.96076648 | 1.482172953 | 5500024025833041107269.C | .H01.G02.E03.D04.B05.A06 |
| betazole | 39.08816571 | 37.63864853 | 1.449517186 | 5500024024213121906564.D | .H07.G08.D10.B11.A12 |
| probucol | 34.89271771 | 33.45797002 | 1.434747692 | 622623112706.D08 | .H07.G08.E09.D10.B11.A12 |
| epivincamine | 36.5148752 | 35.08636713 | 1.428508073 | 640641112706.B08 | .H07.G08.E09.D10.B11.A12 |
| tinidazole | 35.38709443 | 33.96129529 | 1.425799146 | 5500024025833041107269.C | .H07.E09.D10.B11.A12 |
| sulfapyridine | 35.3782681 | 33.96076648 | 1.417501618 | 5500024025833041107269.B | .H01.G02.E03.D04.B05.A06 |
| procarbazine | 35.56014959 | 34.17633596 | 1.383813638 | 5500024030760072207033.C | .H07.G08.E09.B11.A12 |
| cefoperazone | 34.79522354 | 33.45797002 | 1.337253525 | 622623112706.A11 | .H07.G08.E09.D10.B11.A12 |
| suloctidil | 38.33544692 | 37.00786033 | 1.327586586 | 5500024024213121906563.B | .H01.G02.E03.D04.B05.A06 |
| terbutaline | 34.7771468 | 33.4530122 | 1.324134603 | 622623112706.A05 | .H01.G02.E03.B05.A06 |
| oxedrine | 38.55338232 | 37.24787327 | 1.30550905 | 5500024035736031208612.B | .H01.G02.E03.D04.B05.A06 |
| adenosine phosphate | 34.74637614 | 33.45797002 | 1.288406124 | 622623112706.B12 | .H07.G08.E09.D10.B11.A12 |
| minocycline | 34.27960567 | 33.00109826 | 1.278507406 | 5500024024213121906560.B | .H07.G08.E09.D10.B11.A12 |

| | | | | | |
|---|---|---|---|---|---|
| methyldopa | 34.73089855 | 33.45797002 | 1.272928536 | 622623112706.B08 | .H07.G08.E09.D10.B11.A12 |
| tanespimycin | 37.41090461 | 36.14209273 | 1.268811883 | 5500024017802120306174.H | .H01.G02.E03.D04.B05.A06 |
| sodium phenylbutyrat | 36.19585457 | 34.94387722 | 1.251977344 | EC2004030505AA | EC2004030502AA |
| sirolimus | 37.38065671 | 36.14209273 | 1.238563978 | 5500024017802120306174.H | .H01.G02.E03.D04.B05.A06 |
| nitrendipine | 38.47482725 | 37.24471367 | 1.23011358 | 5500024030700072107987.D | .H01.G02.E03.D04.B05.A06 |
| tribenoside | 35.39776883 | 34.17633596 | 1.221432876 | 5500024030760072207033.H | .H07.G08.E09.B11.A12 |
| fenbendazole | 34.66783611 | 33.45935203 | 1.20848408 | 5500024024213121906560.C | .H01.G02.E03.D04.B05.A06 |
| tolazoline | 36.47538332 | 35.2669347 | 1.208448618 | 612613111306.A01 | .H01.G02.E03.D04.B05.A06 |
| terazosin | 35.1686715 | 33.96076648 | 1.207905022 | 5500024025833041107269.C | .H01.G02.E03.D04.B05.A06 |
| leflunomide | 35.15656113 | 33.96076648 | 1.195794652 | 5500024025833041107269.A | .H01.G02.E03.D04.B05.A06 |
| dydrogesterone | 36.21255939 | 35.01997127 | 1.19258812 | 5500024024213121906559.C | .H01.G02.E03.D04.B05.A06 |
| ethotoin | 36.24958436 | 35.05833814 | 1.191246214 | 5500024024213121906559.C | .H07.G08.E09.D10.B11.A12 |
| sirolimus | 36.12983309 | 34.94387722 | 1.18595587 | EC2004030504AA | EC2004030502AA |
| dobutamine | 34.62309094 | 33.45797002 | 1.165120926 | 622623112706.H11 | .H07.G08.E09.D10.B11.A12 |
| thioridazine | 35.57987637 | 34.42118584 | 1.158690534 | 5500024030401071707292.C | .H01.G02.E03.D04.B05.A06 |
| dipivefrine | 36.24446957 | 35.08636713 | 1.158102443 | 640641112706.H09 | .H07.G08.E09.D10.B11.A12 |
| trichostatin A | 34.60433485 | 33.4530122 | 1.151322653 | 622623112706.F06 | .H01.G02.E03.B05.A06 |
| disulfiram | 35.89892333 | 34.74789333 | 1.151030004 | 614615111406.G03 | .H01.G02.E03.D04.B05.A06 |
| clorgiline | 34.59688955 | 33.45797002 | 1.138919535 | 6.23E+21 | .H07.G08.E09.D10.B11.A12 |
| helveticoside | 36.18138784 | 35.05833814 | 1.123049702 | 5500024024213121906559.D | .H07.G08.E09.D10.B11.A12 |
| tanespimycin | 37.25859978 | 36.14209273 | 1.116507047 | 5500024017802120306174.D | .H01.G02.E03.D04.B05.A06 |
| tanespimycin | 37.25782303 | 36.14209273 | 1.115730305 | 5500024017802120306174.D | .H01.G02.E03.D04.B05.A06 |
| sanguinarine | 35.82495099 | 34.70962788 | 1.11532311 | 5500024030760072207033.C | .H01.G02.E03.D04.B05.A06 |
| felodipine | 36.26394616 | 35.15552717 | 1.108418993 | 5500024024214122006604.F | .H01.G02.E03.D04.A06 |
| meptazinol | 35.80590567 | 34.70962788 | 1.096277791 | 5500024030760072207033.G | .H01.G02.E03.D04.B05.A06 |
| kinetin | 35.049997 | 33.96076648 | 1.089230518 | 5500024025833041107269.G | .H01.G02.E03.D04.B05.A06 |
| monorden | 35.50906946 | 34.42118584 | 1.087883627 | 5.50E+22 | .H01.G02.E03.D04.B05.A06 |
| alclometasone | 35.04764119 | 33.96076648 | 1.086874712 | 5500024025833041107269.C | .H01.G02.E03.D04.B05.A06 |
| glafenine | 34.08374321 | 33.00109826 | 1.08264495 | 5.50E+28 | .H07.G08.E09.D10.B11.A12 |
| gossypol | 36.13134996 | 35.05833814 | 1.073011817 | 5500024024213121906559.B | .H07.G08.E09.D10.B11.A12 |
| clofazimine | 34.49988551 | 33.45797002 | 1.041915497 | 622623112706.A08 | .H07.G08.E09.D10.B11.A12 |
| albendazole | 34.49144571 | 33.4530122 | 1.038433508 | 622623112706.H03 | .H01.G02.E03.B05.A06 |
| monorden | 38.35130907 | 37.31290545 | 1.038403621 | 5.50E+28 | .G08.E09.D10.B11.A12 |
| sulfafurazole | 34.47503053 | 33.45797002 | 1.017060511 | 6.23E+19 | .H07.G08.E09.D10.B11.A12 |
| lycorine | 36.0718233 | 35.05833814 | 1.013485162 | 5500024024213121906559.C | .H07.G08.E09.D10.B11.A12 |
| monorden | 37.15160021 | 36.14209273 | 1.009507481 | 5.50E+22 | .H01.G02.E03.D04.B05.A06 |
| haloperidol | 37.14868775 | 36.14209273 | 1.006595018 | 5500024017802120306174.H | .H01.G02.E03.D04.B05.A06 |
| etilefrine | 35.71525263 | 34.70962788 | 1.005624753 | 5500024030760072207033.C | .H01.G02.E03.D04.B05.A06 |
| tranylcypromine | 36.10656156 | 35.10850208 | 0.998059486 | 614615111406.F10 | .H07.G08.E09.D10.B11.A12 |
| monorden | 36.15922886 | 35.1615202 | 0.99770866 | EC2004073018AA | EC2004073014AA |
| napelline | 34.9445334 | 33.96076648 | 0.983766925 | 5.50E+26 | .H01.G02.E03.D04.B05.A06 |
| hexetidine | 36.13273385 | 35.15552717 | 0.977206684 | 5500024024214122006604.A | .H01.G02.E03.D04.A06 |
| dipyridamole | 36.39708991 | 35.4281341 | 0.968955814 | 612613111306.F09 | .H07.G08.E09.D10.B11.A12 |
| astemizole | 35.71358342 | 34.74789333 | 0.965690091 | 614615111406.H04 | .H01.G02.E03.D04.B05.A06 |
| strophanthidin | 34.91156136 | 33.96076648 | 0.950794885 | 5500024025833041107269.D | .H01.G02.E03.D04.B05.A06 |
| anisomycin | 37.9584653 | 37.00786033 | 0.950604969 | 5500024024213121906563.A | .H01.G02.E03.D04.B05.A06 |
| piribedil | 35.12548936 | 34.17633596 | 0.949153399 | 5500024030760072207033.G | .H07.G08.E09.B11.A12 |
| monensin | 34.90808649 | 33.96129529 | 0.946791204 | 5500024025833041107269.A | .H07.E09.D10.B11.A12 |
| atropine oxide | 35.69259977 | 34.74789333 | 0.944706438 | 614615111406.G04 | .H01.G02.E03.D04.B05.A06 |
| flunixin | 34.87564012 | 33.96129529 | 0.914344833 | 5500024025833041107269.G | .H07.E09.D10.B11.A12 |
| hexestrol | 34.85938791 | 33.96076648 | 0.898621437 | 5500024025833041107269.F | .H01.G02.E03.D04.B05.A06 |
| ethaverine | 35.10143555 | 34.21031976 | 0.891115792 | 5500024030760072207028.F | .H07.G08.E09.D10.B11.A12 |
| (+)-chelidonine | 35.95320999 | 35.08636713 | 0.866842861 | 640641112706.A07 | .H07.G08.E09.D10.B11.A12 |
| vidarabine | 35.267069 | 34.40088765 | 0.866181352 | 5500024030401071707292.H | .H07.G08.E09.D10.B11 |
| propantheline bromid | 34.99429429 | 34.13330462 | 0.860989674 | 5500024030760072207028.C | .H01.G02.D04.B05.A06 |
| succinylsulfathiazole | 35.8762855 | 35.01997127 | 0.856314231 | 5500024024213121906559.A | .H01.G02.E03.D04.B05.A06 |
| alexidine | 34.81754385 | 33.96129529 | 0.856248558 | 5500024025833041107269.B | .H07.E09.D10.B11.A12 |
| trichostatin A | 34.80915105 | 33.96129529 | 0.847855759 | 5500024025833041107269.D | .H07.E09.D10.B11.A12 |
| ethoxyquin | 34.80825795 | 33.96129529 | 0.846962658 | 5.50E+31 | .H07.E09.D10.B11.A12 |
| alfuzosin | 34.29654598 | 33.45797002 | 0.83857596 | 622623112706.H08 | .H07.G08.E09.D10.B11.A12 |
| alvespimycin | 35.25844788 | 34.42118584 | 0.837262041 | 5500024030401071707292.F | .H01.G02.E03.D04.B05.A06 |
| mestranol | 34.96869028 | 34.13330462 | 0.835385663 | 5500024030760072207028.D | .H01.G02.D04.B05.A06 |
| trichlormethiazide | 34.96535166 | 34.13330462 | 0.83204704 | 5500024030760072207028.F | .H01.G02.D04.B05.A06 |
| amantadine | 37.67075836 | 36.83996258 | 0.830795777 | 5500024024213121906563.A | .H07.G08.E09.D10.B11.A12 |
| dihydroergocristine | 35.93561975 | 35.11129025 | 0.824329501 | 640641112706.B06 | .H01.G02.E03.D04.B05 |
| 15-delta prostaglandi | 38.13549291 | 37.31290545 | 0.822587458 | 5500024035736031208612.C | .G08.E09.D10.B11.A12 |
| sulfadimidine | 34.77846092 | 33.96129529 | 0.81716563 | 5.50E+32 | .H07.E09.D10.B11.A12 |
| meclofenoxate | 34.76059775 | 33.96129529 | 0.799302458 | 5500024025833041107269.H | .H07.E09.D10.B11.A12 |
| prochlorperazine | 38.10699232 | 37.31290545 | 0.794086867 | 5500024035736031208612.F | .G08.E09.D10.B11.A12 |

| | | | | | |
|---|---|---|---|---|---|
| noretynodrel | 38.43104462 | 37.63864853 | 0.792396088 | 5500024024213121906564.C | .H07.G08.D10.B11.A12 |
| rolipram | 38.03593614 | 37.24471367 | 0.791222466 | 5500024030700072107987.G | .H01.G02.E03.D04.B05.A06 |
| sirolimus | 38.10019147 | 37.31290545 | 0.787286021 | 5.50E+31 | .G08.E09.D10.B11.A12 |
| tanespimycin | 38.09902052 | 37.31290545 | 0.786115066 | 5500024035736031208612.D | .G08.E09.D10.B11.A12 |
| monobenzone | 34.99520176 | 34.21031976 | 0.784882007 | 5500024030760072207028.C | .H07.G08.E09.D10.B11.A12 |
| seneciphylline | 35.80368957 | 35.01997127 | 0.783718301 | 5500024024213121906559.F | .H01.G02.E03.D04.B05.A06 |
| geldanamycin | 38.09466582 | 37.31290545 | 0.781760365 | 5500024035736031208612.D | .G08.E09.D10.B11.A12 |
| hesperetin | 36.19298745 | 35.4281341 | 0.764853349 | 612613111306.C07 | .H07.G08.E09.D10.B11.A12 |
| tanespimycin | 35.1830758 | 34.42118584 | 0.761889967 | 5500024030401071707292.F | .H01.G02.E03.D04.B05.A06 |
| prednisolone | 33.7563712 | 33.00109826 | 0.755272941 | 5500024024213121906560.D | .H07.G08.E09.D10.B11.A12 |
| thioguanosine | 37.76008378 | 37.00786033 | 0.75222345 | 5500024024213121906563.H | .H01.G02.E03.D04.B05.A06 |
| myricetin | 37.58854378 | 36.83996258 | 0.748581199 | 5500024024213121906563.C | .H07.G08.E09.D10.B11.A12 |
| isocarboxazid | 34.7088888 | 33.96129529 | 0.747593512 | 5500024025833041107269.D | .H07.E09.D10.B11.A12 |
| perhexiline | 33.74539706 | 33.00109826 | 0.744298801 | 5500024024213121906560.A | .H07.G08.E09.D10.B11.A12 |
| bucladesine | 35.14206007 | 34.40088765 | 0.741172422 | 5500024030401071707292.A | .H07.G08.E09.D10.B11 |
| wortmannin | 36.88270884 | 36.14209273 | 0.740616116 | 5500024017802120306174.A | .H01.G02.E03.D04.B05.A06 |
| ketanserin | 34.19648861 | 33.45797002 | 0.738518598 | 622623112706.G10 | .H07.G08.E09.D10.B11.A12 |
| Prestwick-1083 | 34.9141842 | 34.17633596 | 0.737848248 | 5500024030760072207033.B | .H07.G08.E09.B11.A12 |
| fulvestrant | 36.87950806 | 36.14209273 | 0.737415327 | 5500024017802120306174.H | .H01.G02.E03.D04.B05.A06 |
| riluzole | 34.19118178 | 33.45935203 | 0.73182975 | 5500024024213121906560.H | .H01.G02.E03.D04.B05.A06 |
| enoxacin | 34.1806638 | 33.45797002 | 0.722693783 | 622623112706.F08 | .H07.G08.E09.D10.B11.A12 |
| epiandrosterone | 35.87780523 | 35.15552717 | 0.722278068 | 5500024024214122006604.D | .H01.G02.E03.D04.A06 |
| acetylsalicylic acid | 36.86361931 | 36.14209273 | 0.721526586 | 5500024017802120306174.H | .H01.G02.E03.D04.B05.A06 |
| pyridoxine | 35.80193583 | 35.08636713 | 0.715568706 | 640641112706.G11 | .H07.G08.E09.D10.B11.A12 |
| arcaine | 34.84653264 | 34.13330462 | 0.713228021 | 5500024030760072207028.C | .H01.G02.D04.B05.A06 |
| trichostatin A | 34.16855489 | 33.45797002 | 0.710584875 | 622623112706.C07 | .H07.G08.E09.D10.B11.A12 |
| isoflupredone | 38.34652553 | 37.63864853 | 0.707877005 | 5500024024213121906564.A | .H07.G08.D10.B11.A12 |
| sirolimus | 36.84428851 | 36.14209273 | 0.702195783 | 5.50E+25 | .H01.G02.E03.D04.B05.A06 |
| tanespimycin | 36.83985847 | 36.14209273 | 0.69776574 | 5500024017802120306174.F | .H01.G02.E03.D04.B05.A06 |
| amphotericin B | 35.85131543 | 35.15552717 | 0.695788266 | 5500024024214122006604.D | .H01.G02.E03.D04.A06 |
| nomifensine | 35.43521246 | 34.74789333 | 0.687319135 | 614615111406.F06 | .H01.G02.E03.D04.B05.A06 |
| chlorphenamine | 35.4336778 | 34.74789333 | 0.685784468 | 614615111406.G05 | .H01.G02.E03.D04.B05.A06 |
| genistein | 37.9976951 | 37.31290545 | 0.684789649 | 5500024035736031208612.B | .G08.E09.D10.B11.A12 |
| tanespimycin | 37.99422912 | 37.31290545 | 0.681323668 | 5500024035736031208612.D | .G08.E09.D10.B11.A12 |
| karakoline | 35.72922332 | 35.05833814 | 0.67088518 | 5500024024213121906559.B | .H07.G08.E09.D10.B11.A12 |
| riboflavin | 35.75443541 | 35.08636713 | 0.668068286 | 6.41E+19 | .H07.G08.E09.D10.B11.A12 |
| sirolimus | 36.80957594 | 36.14209273 | 0.667483214 | 5500024017802120306174.A | .H01.G02.E03.D04.B05.A06 |
| tamoxifen | 35.41073796 | 34.74789333 | 0.662844629 | 614615111406.H05 | .H01.G02.E03.D04.B05.A06 |
| pseudopelletierine | 35.74918615 | 35.08636713 | 0.662819021 | 640641112706.D11 | .H07.G08.E09.D10.B11.A12 |
| famprofazone | 35.72076023 | 35.05833814 | 0.662422086 | 5500024024213121906559.G | .H07.G08.E09.D10.B11.A12 |
| syrosingopine | 35.74130524 | 35.08636713 | 0.654938112 | 640641112706.F07 | .H07.G08.E09.D10.B11.A12 |
| clorsulon | 35.75351534 | 35.11129025 | 0.642225094 | 640641112706.D06 | .H01.G02.E03.D04.B05 |
| thalidomide | 35.74898923 | 35.10850208 | 0.640487156 | 614615111406.G10 | .H07.G08.E09.D10.B11.A12 |
| homochlorcyclizine | 33.63769796 | 33.00109826 | 0.636599701 | 5500024024213121906560.F | .H07.G08.E09.D10.B11.A12 |
| 8-azaguanine | 38.27285393 | 37.63864853 | 0.634205401 | 5500024024213121906564.H | .H07.G08.D10.B11.A12 |
| benfluorex | 37.63020609 | 37.00786033 | 0.622345761 | 5500024024213121906563.H | .H01.G02.E03.D04.B05.A06 |
| prasterone | 37.86655834 | 37.24471367 | 0.621844672 | 5500024030700072107987.B | .H01.G02.E03.D04.B05.A06 |
| lanatoside C | 35.67985798 | 35.05833814 | 0.621519837 | 5500024024213121906559.D | .H07.G08.E09.D10.B11.A12 |
| diazoxide | 35.3657685 | 34.74789333 | 0.617875174 | 614615111406.G01 | .H01.G02.E03.D04.B05.A06 |
| genistein | 35.03766741 | 34.42118584 | 0.616481571 | 5500024030401071707292.B | .H01.G02.E03.D04.B05.A06 |
| zimeldine | 36.03898119 | 35.4281341 | 0.610847094 | 612613111306.G10 | .H07.G08.E09.D10.B11.A12 |
| practolol | 34.06755525 | 33.45797002 | 0.609585229 | 622623112706.H09 | .H07.G08.E09.D10.B11.A12 |
| tanespimycin | 37.91973731 | 37.31290545 | 0.606831858 | 5500024035736031208612.F | .G08.E09.D10.B11.A12 |
| fludroxycortide | 35.66496125 | 35.05833814 | 0.606623113 | 5.50E+28 | .H07.G08.E09.D10.B11.A12 |
| spiperone | 34.05813599 | 33.4530122 | 0.605123787 | 622623112706.F04 | .H01.G02.E03.B05.A06 |
| nordihydroguaiaretic a | 36.74215328 | 36.14209273 | 0.600060552 | 5.50E+27 | .H01.G02.E03.D04.B05.A06 |
| hycanthone | 34.05790944 | 33.45797002 | 0.59993942 | 622623112706.C09 | .H07.G08.E09.D10.B11.A12 |
| geldanamycin | 36.74037057 | 36.14209273 | 0.598277838 | 5500024017802120306174.D | .H01.G02.E03.D04.B05.A06 |
| thiamazole | 34.559231 | 33.96129529 | 0.597935713 | 5500024025833041107269.C | .H07.E09.D10.B11.A12 |
| scopolamine | 34.72791807 | 34.13330462 | 0.59461345 | 5500024030760072207028.B | .H01.G02.D04.B05.A06 |
| halcinonide | 35.65117568 | 35.05833814 | 0.592837537 | 5.50E+29 | .H07.G08.E09.D10.B11.A12 |
| monocrotaline | 35.6782273 | 35.08636713 | 0.591860175 | 640641112706.G09 | .H07.G08.E09.D10.B11.A12 |
| S-propranolol | 34.76790996 | 34.17633596 | 0.591574006 | 5.50E+33 | .H07.G08.E09.B11.A12 |
| mepacrine | 34.04252126 | 33.4530122 | 0.589509056 | 6.23E+13 | .H01.G02.E03.B05.A06 |
| nystatin | 35.71949145 | 35.13162255 | 0.587868895 | 5500024024214122006604.A | .H07.G08.E09.D10.B11.A12 |
| cloperastine | 34.5486814 | 33.96129529 | 0.587386113 | 5500024025833041107269.G | .H07.E09.D10.B11.A12 |
| ritodrine | 37.59247434 | 37.00786033 | 0.584614011 | 5.50E+23 | .H01.G02.E03.D04.B05.A06 |
| tolazamide | 35.71511685 | 35.13162255 | 0.5834943 | 5.50E+33 | .H07.G08.E09.D10.B11.A12 |
| tretinoin | 36.7741938 | 36.19585457 | 0.578339229 | EC2004032013AA | EC2004032009AA |

| | | | | | |
|---|---|---|---|---|---|
| naltrexone | 35.32314641 | 34.74789333 | 0.575253086 | 614615111406.H02 | .H01.G02.E03.D04.B05.A06 |
| furosemide | 34.02773041 | 33.4530122 | 0.574718209 | 622623112706.B06 | .H01.G02.E03.B05.A06 |
| oxytetracycline | 34.02769716 | 33.4530122 | 0.574684957 | 622623112706.G04 | .H01.G02.E03.B05.A06 |
| isoxsuprine | 35.83649986 | 35.2669347 | 0.569565158 | 612613111306.D02 | .H01.G02.E03.D04.B05.A06 |
| Prestwick-685 | 35.62753583 | 35.05833814 | 0.569197692 | 5.50E+33 | .H07.G08.E09.D10.B11.A12 |
| cefmetazole | 34.52638719 | 33.96076648 | 0.56562071 | 5500024025833041107269.D | .H01.G02.E03.D04.B05.A06 |
| pramocaine | 35.62365293 | 35.05833814 | 0.565314784 | 5500024024213121906559.C | .H07.G08.E09.D10.B11.A12 |
| isosorbide | 35.61874537 | 35.05833814 | 0.560407233 | 5500024024213121906559.F | .H07.G08.E09.D10.B11.A12 |
| isoconazole | 35.30703018 | 34.74789333 | 0.559136856 | 614615111406.G06 | .H01.G02.E03.D04.B05.A06 |
| sulconazole | 34.76884126 | 34.21031976 | 0.558521504 | 5500024030760072207028.F | .H07.G08.E09.D10.B11.A12 |
| naftifine | 34.7309977 | 34.17633596 | 0.554661745 | 5500024030760072207033.B | .H07.G08.E09.B11.A12 |
| altizide | 34.50738611 | 33.96076648 | 0.546619635 | 5500024025833041107269.D | .H01.G02.E03.D04.B05.A06 |
| etiocholanolone | 35.60466483 | 35.05833814 | 0.546326693 | 5500024024213121906559.A | .H07.G08.E09.D10.B11.A12 |
| fendiline | 33.99399733 | 33.4530122 | 0.540985126 | 622623112706.C03 | .H01.G02.E03.B05.A06 |
| beta-escin | 35.59642648 | 35.05833814 | 0.538088336 | 5500024024213121906559.C | .H07.G08.E09.D10.B11.A12 |
| scoulerine | 35.6484449 | 35.11129025 | 0.537154645 | 640641112706.B01 | .H01.G02.E03.D04.B05 |
| hydroquinine | 35.62128647 | 35.08636713 | 0.534919346 | 640641112706.D12 | .H07.G08.E09.D10.B11.A12 |
| benzethonium chlorid | 34.49512528 | 33.96076648 | 0.534358804 | 5500024025833041107269.G | .H01.G02.E03.D04.B05.A06 |
| metitepine | 33.98414598 | 33.45797002 | 0.526175965 | 622623112706.C11 | .H07.G08.E09.D10.B11.A12 |
| alpha-estradiol | 36.66821918 | 36.14209273 | 0.526126453 | 5500024017802120306174.G | .H01.G02.E03.D04.B05.A06 |
| 3-hydroxy-DL-kynurei | 37.53389624 | 37.00786033 | 0.526035912 | 5500024024213121906563.A | .H01.G02.E03.D04.B05.A06 |
| metanephrine | 35.95272 | 35.4281341 | 0.524585904 | 612613111306.F07 | .H07.G08.E09.D10.B11.A12 |
| butamben | 34.48262707 | 33.96076648 | 0.521860595 | 5500024025833041107269.C | .H01.G02.E03.D04.B05.A06 |
| N6-methyladenosine | 37.52815045 | 37.00786033 | 0.520290118 | 5500024024213121906563.G | .H01.G02.E03.D04.B05.A06 |
| demeclocycline | 34.4791273 | 33.96129529 | 0.517832007 | 5500024025833041107269.H | .H07.E09.D10.B11.A12 |
| metergoline | 33.97461272 | 33.45797002 | 0.516642707 | 6.23E+23 | .H07.G08.E09.D10.B11.A12 |
| gemfibrozil | 35.61971284 | 35.10850208 | 0.511210767 | 614615111406.C07 | .H07.G08.E09.D10.B11.A12 |
| naringin | 35.66183818 | 35.15552717 | 0.506311015 | 5500024024214122006604.H | .H01.G02.E03.D04.A06 |
| zalcitabine | 35.21489349 | 34.70962788 | 0.505265604 | 5500024030760072207033.B | .H01.G02.E03.D04.B05.A06 |
| etacrynic acid | 33.95771786 | 33.4530122 | 0.504705659 | 6.23E+16 | .H01.G02.E03.B05.A06 |
| bambuterol | 33.95723066 | 33.4530122 | 0.504218461 | 622623112706.A02 | .H01.G02.E03.B05.A06 |
| arecoline | 37.5117442 | 37.00786033 | 0.503883872 | 5500024024213121906563.A | .H01.G02.E03.D04.B05.A06 |
| clomipramine | 33.95487233 | 33.4530122 | 0.501860125 | 6.23E+17 | .H01.G02.E03.B05.A06 |
| fluphenazine | 37.81345797 | 37.31290545 | 0.50055252 | 5500024035736031208612.B | .G08.E09.D10.B11.A12 |
| estropipate | 34.46070831 | 33.96076648 | 0.499941836 | 5500024025833041107269.H | .H01.G02.E03.D04.B05.A06 |
| carcinine | 37.33828066 | 36.83996258 | 0.498318084 | 5500024024213121906563.H | .H07.G08.E09.D10.B11.A12 |
| cinchocaine | 35.75253337 | 35.2669347 | 0.485598666 | 612613111306.G03 | .H01.G02.E03.D04.B05.A06 |
| mianserin | 35.23212454 | 34.74789333 | 0.484231208 | 614615111406.D02 | .H01.G02.E03.D04.B05.A06 |
| atropine | 35.56859761 | 35.08636713 | 0.482230481 | 6.41E+21 | .H07.G08.E09.D10.B11.A12 |
| piperacetazine | 37.72711972 | 37.24787327 | 0.479246444 | 5500024035736031208612.C | .H01.G02.E03.D04.B05.A06 |
| prochlorperazine | 34.89801171 | 34.42118584 | 0.476825876 | 5500024030401071707292.F | .H01.G02.E03.D04.B05.A06 |
| zaprinast | 33.93112166 | 33.45797002 | 0.47315164 | 622623112706.D12 | .H07.G08.E09.D10.B11.A12 |
| hydrocotarnine | 35.55577107 | 35.08636713 | 0.469403938 | 640641112706.D08 | .H07.G08.E09.D10.B11.A12 |
| nordihydroguaiaretic a | 34.8899033 | 34.42118584 | 0.468717464 | 5.50E+27 | .H01.G02.E03.D04.B05.A06 |
| sirolimus | 37.78157442 | 37.31290545 | 0.468668964 | 5500024035736031208612.H | .G08.E09.D10.B11.A12 |
| cetirizine | 35.59342105 | 35.13162255 | 0.461798494 | 5500024024214122006604.G | .H07.G08.E09.D10.B11.A12 |
| estradiol | 36.60303743 | 36.14209273 | 0.460944704 | 5500024017802120306174.G | .H01.G02.E03.D04.B05.A06 |
| fluticasone | 35.16707276 | 34.70962788 | 0.457444884 | 5500024030760072207033.C | .H01.G02.E03.D04.B05.A06 |
| fulvestrant | 37.7655889 | 37.31290545 | 0.452683445 | 5500024035736031208612.B | .G08.E09.D10.B11.A12 |
| trihexyphenidyl | 35.47122921 | 35.01997127 | 0.451257943 | 5500024024213121906559.B | .H01.G02.E03.D04.B05.A06 |
| cyanocobalamin | 37.28830596 | 36.83996258 | 0.448343384 | 5500024024213121906563.F | .H07.G08.E09.D10.B11.A12 |
| clioquinol | 37.69105896 | 37.24471367 | 0.446345288 | 5.50E+26 | .H01.G02.E03.D04.B05.A06 |
| terfenadine | 35.19272003 | 34.74789333 | 0.444826699 | 6.15E+15 | .H01.G02.E03.D04.B05.A06 |
| triflusal | 35.55483573 | 35.11129025 | 0.44354548 | 640641112706.G04 | .H01.G02.E03.D04.B05 |
| wortmannin | 37.7522501 | 37.31290545 | 0.439344649 | 5500024035736031208612.A | .G08.E09.D10.B11.A12 |
| nitrofural | 35.59474325 | 35.15552717 | 0.439216087 | 5500024024214122006604.A | .H01.G02.E03.D04.A06 |
| pheneticillin | 34.39948686 | 33.96076648 | 0.438720379 | 5500024025833041107269.A | .H01.G02.E03.D04.B05.A06 |
| fenofibrate | 33.43929933 | 33.00109826 | 0.438201071 | 5500024024213121906560.C | .H07.G08.E09.D10.B11.A12 |
| ranitidine | 35.54536061 | 35.10850208 | 0.436858536 | 614615111406.H08 | .H07.G08.E09.D10.B11.A12 |
| epitiostanol | 35.1444931 | 34.70962788 | 0.434865223 | 5500024030760072207033.D | .H01.G02.E03.D04.B05.A06 |
| benzathine benzylper | 35.14394544 | 34.70962788 | 0.434317554 | 5500024030760072207033.A | .H01.G02.E03.D04.B05.A06 |
| naringenin | 37.27425794 | 36.83996258 | 0.434295361 | 5500024024213121906563.A | .H07.G08.E09.D10.B11.A12 |
| ramifenazone | 34.39249489 | 33.96076648 | 0.431728416 | 5500024025833041107269.B | .H01.G02.E03.D04.B05.A06 |
| oxybenzone | 37.67621872 | 37.24471367 | 0.431505045 | 5500024030700072107987.C | .H01.G02.E03.D04.B05.A06 |
| carbenoxolone | 34.56310345 | 34.13330462 | 0.429798832 | 5500024030760072207028.B | .H01.G02.D04.B05.A06 |
| sirolimus | 37.74114553 | 37.31290545 | 0.428240077 | 5500024035736031208612.A | .G08.E09.D10.B11.A12 |
| noscapine | 35.51424186 | 35.08636713 | 0.427874733 | 640641112706.H10 | .H07.G08.E09.D10.B11.A12 |
| mometasone | 35.53869827 | 35.11129025 | 0.427408016 | 640641112706.A01 | .H01.G02.E03.D04.B05 |
| betahistine | 35.55787998 | 35.13162255 | 0.426257424 | 5500024024214122006604.F | .H07.G08.E09.D10.B11.A12 |

| | | | | | |
|---|---|---|---|---|---|
| fusidic acid | 37.43272858 | 37.00786033 | 0.424868254 | 5500024024213121906563.C | .H01.G02.E03.D04.B05.A06 |
| desipramine | 33.88177664 | 33.45797002 | 0.423806625 | 622623112706.F07 | .H07.G08.E09.D10.B11.A12 |
| abamectin | 34.37997495 | 33.96076648 | 0.419208473 | 5.50E+22 | .H01.G02.E03.D04.B05.A06 |
| canadine | 35.43698806 | 35.01997127 | 0.417016786 | 5500024024213121906559.A | .H01.G02.E03.D04.B05.A06 |
| methotrexate | 35.84507929 | 35.4281341 | 0.416945189 | 612613111306.B12 | .H07.G08.E09.D10.B11.A12 |
| triflupromazine | 38.05548105 | 37.63864853 | 0.416832523 | 5500024024213121906564.D | .H07.G08.D10.B11.A12 |
| androsterone | 37.42427549 | 37.00786033 | 0.41641516 | 5500024024213121906563.B | .H01.G02.E03.D04.B05.A06 |
| gliquidone | 37.87393465 | 37.46016387 | 0.413770777 | 5500024030700072107987.D | .H07.G08.E09.D10.B11.A12 |
| flunarizine | 33.41153801 | 33.00109826 | 0.41043975 | 5500024024213121906560.F | .H07.G08.E09.D10.B11.A12 |
| alvespimycin | 36.55161251 | 36.14209273 | 0.409519782 | 5500024017802120306174.F | .H01.G02.E03.D04.B05.A06 |
| azaperone | 37.6560947 | 37.24787327 | 0.408221428 | 5500024035736031208612.C | .H01.G02.E03.D04.B05.A06 |
| urapidil | 37.64957319 | 37.24471367 | 0.404859523 | 5500024030700072107987.F | .H01.G02.E03.D04.B05.A06 |
| cyclobenzaprine | 37.24409511 | 36.83996258 | 0.404132536 | 5500024024213121906563.C | .H07.G08.E09.D10.B11.A12 |
| lovastatin | 35.53309492 | 35.13162255 | 0.40147237 | 5500024024214122006604.B | .H07.G08.E09.D10.B11.A12 |
| dexamethasone | 35.14869933 | 34.74789333 | 0.400806005 | 614615111406.B02 | .H01.G02.E03.D04.B05.A06 |
| ricinine | 34.35916224 | 33.96076648 | 0.39839576 | 5500024025833041107269.H | .H01.G02.E03.D04.B05.A06 |
| spectinomycin | 34.52704451 | 34.13330462 | 0.393739893 | 5500024030760072207028.H | .H01.G02.D04.B05.A06 |
| megestrol | 37.63684356 | 37.24471367 | 0.39212989 | 5500024030700072107987.C | .H01.G02.E03.D04.B05.A06 |
| coralyne | 37.39920972 | 37.00786033 | 0.391349389 | 5500024024213121906563.B | .H01.G02.E03.D04.B05.A06 |
| sulfametoxydiazine | 34.79168156 | 34.40088765 | 0.390793916 | 5500024030401071707292.G | .H07.G08.E09.D10.B11 |
| thioridazine | 35.65600009 | 35.2669347 | 0.389065388 | 612613111306.D03 | .H01.G02.E03.D04.B05.A06 |
| nicardipine | 33.84534335 | 33.45797002 | 0.38737333 | 622623112706.F11 | .H07.G08.E09.D10.B11.A12 |
| digitoxigenin | 37.22600045 | 36.83996258 | 0.386037867 | 5500024024213121906563.B | .H07.G08.E09.D10.B11.A12 |
| estradiol | 34.80676894 | 34.42118584 | 0.385583108 | 5500024030401071707292.G | .H01.G02.E03.D04.B05.A06 |
| tetrandrine | 34.34456867 | 33.96076648 | 0.383802192 | 5.50E+23 | .H01.G02.E03.D04.B05.A06 |
| Prestwick-559 | 35.49491423 | 35.11129025 | 0.383623982 | 6.41E+15 | .H01.G02.E03.D04.B05 |
| prochlorperazine | 36.52387802 | 36.14209273 | 0.381785296 | 5500024017802120306174.F | .H01.G02.E03.D04.B05.A06 |
| cefazolin | 34.34028334 | 33.96129529 | 0.378988049 | 5500024025833041107269.D | .H07.E09.D10.B11.A12 |
| amiloride | 35.64588671 | 35.2669347 | 0.378952002 | 612613111306.G04 | .H01.G02.E03.D04.B05.A06 |
| beclometasone | 34.51019659 | 34.13330462 | 0.376891971 | 5.50E+25 | .H01.G02.D04.B05.A06 |
| etanidazole | 34.33732281 | 33.96076648 | 0.376556332 | 5500024025833041107269.G | .H01.G02.E03.D04.B05.A06 |
| gallamine triethiodide | 35.12410601 | 34.74789333 | 0.376212682 | 614615111406.F03 | .H01.G02.E03.D04.B05.A06 |
| etynodiol | 37.61946375 | 37.24471367 | 0.374750083 | 5500024030700072107987.A | .H01.G02.E03.D04.B05.A06 |
| 10-methoxyharmalan | 35.48602664 | 35.11129025 | 0.374736386 | 640641112706.B03 | .H01.G02.E03.D04.B05 |
| flavoxate | 33.37364786 | 33.00109826 | 0.3725496 | 5500024024213121906560.H | .H07.G08.E09.D10.B11.A12 |
| artemisinin | 35.48347351 | 35.11129025 | 0.372183256 | 640641112706.H06 | .H01.G02.E03.D04.B05 |
| calycanthine | 35.45791681 | 35.08636713 | 0.371549687 | 640641112706.D07 | .H07.G08.E09.D10.B11.A12 |
| troleandomycin | 35.63773148 | 35.2669347 | 0.370796773 | 612613111306.H04 | .H01.G02.E03.D04.B05.A06 |
| lithocholic acid | 34.33155219 | 33.96129529 | 0.370256897 | 5500024025833041107269.C | .H07.E09.D10.B11.A12 |
| doxazosin | 34.58022057 | 34.21031976 | 0.369900815 | 5500024030760072207028.H | .H07.G08.E09.D10.B11.A12 |
| raubasine | 35.48036524 | 35.11129025 | 0.36907499 | 640641112706.A03 | .H01.G02.E03.D04.B05 |
| thiethylperazine | 37.61605625 | 37.24787327 | 0.368182973 | 5500024035736031208612.B | .H01.G02.E03.D04.B05.A06 |
| indoprofen | 34.49867569 | 34.13330462 | 0.365371075 | 5500024030760072207028.D | .H01.G02.D04.B05.A06 |
| piperidolate | 37.61241011 | 37.24787327 | 0.364536835 | 5500024035736031208612.G | .H01.G02.E03.D04.B05.A06 |
| luteolin | 34.57369658 | 34.21031976 | 0.36337682 | 5.50E+29 | .H07.G08.E09.D10.B11.A12 |
| benfotiamine | 35.41994013 | 35.05833814 | 0.361601984 | 5500024024213121906559.G | .H07.G08.E09.D10.B11.A12 |
| disopyramide | 33.36267214 | 33.00109826 | 0.361573877 | 5500024024213121906560.A | .H07.G08.E09.D10.B11.A12 |
| tanespimycin | 34.78233684 | 34.42118584 | 0.361151007 | 5500024030401071707292.D | .H01.G02.E03.D04.B05.A06 |
| terconazole | 35.49250883 | 35.13162255 | 0.36088628 | 5500024024214122006604.D | .H07.G08.E09.D10.B11.A12 |
| piperine | 37.19944927 | 36.83996258 | 0.359486696 | 5500024024213121906563.D | .H07.G08.E09.D10.B11.A12 |
| iohexol | 35.51458896 | 35.15552717 | 0.359061794 | 5500024024214122006604.A | .H01.G02.E03.D04.A06 |
| enalapril | 33.35958708 | 33.00109826 | 0.358488821 | 5500024024213121906560.C | .H07.G08.E09.D10.B11.A12 |
| rolitetracycline | 34.56826548 | 34.21031976 | 0.357945724 | 5500024030760072207028.G | .H07.G08.E09.D10.B11.A12 |
| fulvestrant | 36.49998052 | 36.14209273 | 0.357887792 | 5500024017802120306174.B | .H01.G02.E03.D04.B05.A06 |
| viomycin | 34.5326222 | 34.17633596 | 0.356286239 | 5500024030760072207033.A | .H07.G08.E09.B11.A12 |
| estradiol | 37.66648356 | 37.31290545 | 0.35357811 | 5500024035736031208612.A | .G08.E09.D10.B11.A12 |
| neomycin | 35.10142786 | 34.74789333 | 0.353534534 | 6.15E+17 | .H01.G02.E03.D04.B05.A06 |
| acemetacin | 33.35308632 | 33.00109826 | 0.351988058 | 5500024024213121906560.A | .H07.G08.E09.D10.B11.A12 |
| yohimbine | 35.43822305 | 35.08636713 | 0.351855926 | 640641112706.F09 | .H07.G08.E09.D10.B11.A12 |
| cisapride | 35.50609084 | 35.15552717 | 0.350563673 | 5500024024214122006604.D | .H01.G02.E03.D04.A06 |
| tolbutamide | 33.80933574 | 33.45935203 | 0.349983714 | 5500024024213121906560.C | .H01.G02.E03.D04.B05.A06 |
| trifluoperazine | 36.49169112 | 36.14209273 | 0.349598389 | 5500024017802120306174.D | .H01.G02.E03.D04.B05.A06 |
| ethambutol | 35.61518903 | 35.2669347 | 0.348254322 | 6.13E+15 | .H01.G02.E03.D04.B05.A06 |
| etodolac | 35.45632802 | 35.10850208 | 0.347825944 | 614615111406.H11 | .H07.G08.E09.D10.B11.A12 |
| metixene | 35.50190433 | 35.15552717 | 0.346377162 | 5500024024214122006604.C | .H01.G02.E03.D04.A06 |
| bisoprolol | 37.35351787 | 37.00786033 | 0.345657541 | 5500024024213121906563.D | .H01.G02.E03.D04.B05.A06 |
| R-atenolol | 35.47632857 | 35.13162255 | 0.344706021 | 5500024024214122006604.B | .H07.G08.E09.D10.B11.A12 |
| SR-95531 | 37.184414 | 36.83996258 | 0.344451421 | 5500024024213121906563.F | .H07.G08.E09.D10.B11.A12 |
| norfloxacin | 35.4511669 | 35.10850208 | 0.34266482 | 614615111406.H10 | .H07.G08.E09.D10.B11.A12 |

| | | | | | |
|---|---|---|---|---|---|
| proadifen | 34.74208912 | 34.40088765 | 0.341201474 | 5500024030401071707292.H | .H07.G08.E09.D10.B11 |
| sulfanilamide | 34.741365 | 34.40088765 | 0.340477348 | 5500024030401071707292.G | .H07.G08.E09.D10.B11 |
| cefsulodin | 34.47355288 | 34.13330462 | 0.340248266 | 5500024030760072207028.G | .H01.G02.D04.B05.A06 |
| ribostamycin | 34.73968054 | 34.40088765 | 0.338792895 | 5500024030401071707292.H | .H07.G08.E09.D10.B11 |
| cefadroxil | 37.1784657 | 36.83996258 | 0.338503125 | 5.50E+31 | .H07.G08.E09.D10.B11.A12 |
| Prestwick-689 | 34.29907614 | 33.96076648 | 0.338309659 | 5500024025833041107269.F( | .H01.G02.E03.D04.B05.A06 |
| tanespimycin | 34.75796603 | 34.42118584 | 0.336780194 | 5500024030401071707292.H | .H01.G02.E03.D04.B05.A06 |
| sulfadimethoxine | 34.29617986 | 33.96129529 | 0.334884574 | 5500024025833041107269.A( | .H07.E09.D10.B11.A12 |
| naftopidil | 35.04395602 | 34.70962788 | 0.334328139 | 5500024030760072207033.F( | .H01.G02.E03.D04.B05.A06 |
| dextromethorphan | 37.34207477 | 37.00786033 | 0.334214443 | 5.50E+25 | .H01.G02.E03.D04.B05.A06 |
| adrenosterone | 37.79412535 | 37.46016387 | 0.333961477 | 5500024030700072107987.H | .H07.G08.E09.D10.B11.A12 |
| trazodone | 33.33454348 | 33.00109826 | 0.333445219 | 5500024024213121906560.G | .H07.G08.E09.D10.B11.A12 |
| pipemidic acid | 37.57556025 | 37.24471367 | 0.330846576 | 5500024030700072107987.C | .H01.G02.E03.D04.B05.A06 |
| cefotiam | 35.48416268 | 35.15552717 | 0.328635517 | 5500024024214122006604.A( | .H01.G02.E03.D04.A06 |
| rescinnamine | 35.34806787 | 35.01997127 | 0.328096602 | 5500024024213121906559.H | .H01.G02.E03.D04.B05.A06 |
| cinchonidine | 35.41118563 | 35.08636713 | 0.324818501 | 640641112706.C11 | .H07.G08.E09.D10.B11.A12 |
| spiramycin | 34.28567047 | 33.96129529 | 0.324375186 | 5.50E+28 | .H07.E09.D10.B11.A12 |
| dilazep | 33.78270225 | 33.45935203 | 0.323350218 | 5500024024213121906560.H | .H01.G02.E03.D04.B05.A06 |
| vitexin | 35.34099467 | 35.01997127 | 0.321023404 | 5500024024213121906559.C | .H01.G02.E03.D04.B05.A06 |
| nifenazone | 35.42783726 | 35.10850208 | 0.319335189 | 614615111406.B10 | .H07.G08.E09.D10.B11.A12 |
| troglitazone | 36.4593759 | 36.14209273 | 0.317283173 | 5500024017802120306174.C | .H01.G02.E03.D04.B05.A06 |
| benperidol | 35.44867006 | 35.13162255 | 0.317047504 | 5500024024214122006604.F | .H07.G08.E09.D10.B11.A12 |
| piretanide | 37.56460247 | 37.24787327 | 0.316729197 | 5500024035736031208612.D | .H01.G02.E03.D04.B05.A06 |
| difenidol | 33.31614488 | 33.00109826 | 0.315046612 | 5500024024213121906560.H | .H07.G08.E09.D10.B11.A12 |
| dacarbazine | 35.40050081 | 35.08636713 | 0.314133681 | 640641112706.F08 | .H07.G08.E09.D10.B11.A12 |
| fluorocurarine | 34.27424115 | 33.96076648 | 0.313474677 | 5.50E+25 | .H01.G02.E03.D04.B05.A06 |
| benzonatate | 37.95103263 | 37.63864853 | 0.312384101 | 5500024024213121906564.F( | .H07.G08.D10.B11.A12 |
| N-acetylmuramic acic | 37.15137168 | 36.83996258 | 0.311409101 | 5500024024213121906563.D | .H07.G08.E09.D10.B11.A12 |
| oxyphenbutazone | 37.55910916 | 37.24787327 | 0.311235887 | 5500024035736031208612.A( | .H01.G02.E03.D04.B05.A06 |
| nabumetone | 37.770286 | 37.46016387 | 0.310122129 | 5500024030700072107987.H | .H07.G08.E09.D10.B11.A12 |
| carbachol | 34.52017387 | 34.21031976 | 0.309854113 | 5.50E+31 | .H07.G08.E09.D10.B11.A12 |
| ronidazole | 37.55687728 | 37.24787327 | 0.309004008 | 5500024035736031208612.F( | .H01.G02.E03.D04.B05.A06 |
| ivermectin | 35.05595801 | 34.74789333 | 0.308064681 | 614615111406.H06 | .H01.G02.E03.D04.B05.A06 |
| dapsone | 37.94633459 | 37.63864853 | 0.307686057 | 5500024024213121906564.B | .H07.G08.D10.B11.A12 |
| merbromin | 34.26737564 | 33.96129529 | 0.306080349 | 5500024025833041107269.A( | .H07.E09.D10.B11.A12 |
| dimethadione | 34.51598771 | 34.21031976 | 0.305667949 | 5500024030760072207028.G | .H07.G08.E09.D10.B11.A12 |
| primaquine | 37.14437926 | 36.83996258 | 0.304416682 | 5500024024213121906563.A( | .H07.G08.E09.D10.B11.A12 |
| cyproheptadine | 35.73157168 | 35.4281341 | 0.303437585 | 6.13E+18 | .H07.G08.E09.D10.B11.A12 |
| prednisone | 35.5700989 | 35.2669347 | 0.303164194 | 612613111306.F06 | .H01.G02.E03.D04.B05.A06 |
| benzbromarone | 37.30894268 | 37.00786033 | 0.301082348 | 5500024024213121906563.C | .H01.G02.E03.D04.B05.A06 |
| tremorine | 33.75276479 | 33.4530122 | 0.299752587 | 622623112706.B04 | .H01.G02.E03.B05.A06 |
| stachydrine | 35.38534073 | 35.08636713 | 0.298973601 | 640641112706.H08 | .H07.G08.E09.D10.B11.A12 |
| iocetamic acid | 34.42847179 | 34.13330462 | 0.295167174 | 5500024030760072207028.A( | .H01.G02.D04.B05.A06 |
| mexiletine | 33.75384478 | 33.45935203 | 0.294492756 | 5500024024213121906560.B( | .H01.G02.E03.D04.B05.A06 |
| clemastine | 33.29505652 | 33.00109826 | 0.293958252 | 5500024024213121906560.A | .H07.G08.E09.D10.B11.A12 |
| vinburnine | 35.37916753 | 35.08636713 | 0.2928004 | 640641112706.A09 | .H07.G08.E09.D10.B11.A12 |
| cefapirin | 34.69263782 | 34.40088765 | 0.291750168 | 5500024030401071707292.C | .H07.G08.E09.D10.B11 |
| levothyroxine sodium | 37.13077467 | 36.83996258 | 0.290812094 | 5500024024213121906563.G | .H07.G08.E09.D10.B11.A12 |
| streptozocin | 34.25073402 | 33.96076648 | 0.289967542 | 5500024025833041107269.B( | .H01.G02.E03.D04.B05.A06 |
| tetrahydroalstonine | 35.37616358 | 35.08636713 | 0.289796454 | 640641112706.G07 | .H07.G08.E09.D10.B11.A12 |
| acebutolol | 35.55627874 | 35.2669347 | 0.289344032 | 612613111306.C05 | .H01.G02.E03.D04.B05.A06 |
| trichostatin A | 33.2901582 | 33.00109826 | 0.289059935 | 5500024024213121906560.H | .H07.G08.E09.D10.B11.A12 |
| hymecromone | 34.49916319 | 34.21031976 | 0.28884343 | 5500024030760072207028.D | .H07.G08.E09.D10.B11.A12 |
| estradiol | 34.70641817 | 34.42118584 | 0.28523233 | 5500024030401071707292.A( | .H01.G02.E03.D04.B05.A06 |
| glimepiride | 35.30515732 | 35.01997127 | 0.285186054 | 5500024024213121906559.C | .H01.G02.E03.D04.B05.A06 |
| metrizamide | 37.12421106 | 36.83996258 | 0.284248484 | 5500024024213121906563.F | .H07.G08.E09.D10.B11.A12 |
| glipizide | 35.71139785 | 35.4281341 | 0.283263754 | 612613111306.H11 | .H07.G08.E09.D10.B11.A12 |
| trichostatin A | 36.42498967 | 36.14209273 | 0.282896942 | 5500024017802120306174.G | .H01.G02.E03.D04.B05.A06 |
| chenodeoxycholic aci | 33.28355158 | 33.00109826 | 0.28245332 | 5500024024213121906560.C | .H07.G08.E09.D10.B11.A12 |
| griseofulvin | 33.74166329 | 33.45935203 | 0.282311263 | 5500024024213121906560.H | .H01.G02.E03.D04.B05.A06 |
| trifluoperazine | 34.70019429 | 34.42118584 | 0.279008451 | 5500024030401071707292.D | .H01.G02.E03.D04.B05.A06 |
| ganciclovir | 34.48913801 | 34.21031976 | 0.278818256 | 5500024030760072207028.G | .H07.G08.E09.D10.B11.A12 |
| cyclopenthiazide | 34.98818433 | 34.70962788 | 0.27855645 | 5500024030760072207033.H | .H01.G02.E03.D04.B05.A06 |
| imipramine | 37.91595489 | 37.63864853 | 0.277306365 | 5.50E+28 | .H07.G08.D10.B11.A12 |
| praziquantel | 33.73004652 | 33.4530122 | 0.277034314 | 622623112706.C02 | .H01.G02.E03.B05.A06 |
| benzylpenicillin | 37.52433834 | 37.24787327 | 0.276465063 | 5500024035736031208612.B( | .H01.G02.E03.D04.B05.A06 |
| proguanil | 34.45207926 | 34.17633596 | 0.275743307 | 5500024030760072207033.H | .H07.G08.E09.B11.A12 |
| mephenytoin | 37.52191089 | 37.24787327 | 0.274037613 | 5500024035736031208612.A( | .H01.G02.E03.D04.B05.A06 |
| quipazine | 35.36000288 | 35.08636713 | 0.273635757 | 640641112706.A10 | .H07.G08.E09.D10.B11.A12 |

| | | | | | |
|---|---|---|---|---|---|
| propoxycaine | 37.52150503 | 37.24787327 | 0.273631759 | 550002403573603120861 2.A | .H01.G02.E03.D04.B05.A06 |
| tonzonium bromide | 37.51773545 | 37.24471367 | 0.273021778 | 5500024030700072107987.F | .H01.G02.E03.D04.B05.A06 |
| methylprednisolone | 33.72533409 | 33.4530122 | 0.272321889 | 622623112706.D01 | .H01.G02.E03.B05.A06 |
| LY-294002 | 36.4144016 | 36.14209273 | 0.272308874 | 5500024017802120306174.D | .H01.G02.E03.D04.B05.A06 |
| fluoxetine | 35.42681724 | 35.15552717 | 0.271290077 | 5500024024214122006604.B | .H01.G02.E03.D04.A06 |
| galantamine | 35.29110488 | 35.01997127 | 0.271133614 | 5500024024213121906559.H | .H01.G02.E03.D04.B05.A06 |
| tetramisole | 35.40112368 | 35.13162255 | 0.269501129 | 5500024024214122006604.C | .H07.G08.E09.D10.B11.A12 |
| theobromine | 34.4007191 | 34.13330462 | 0.267414488 | 5500024030760072207028.F | .H01.G02.D04.B05.A06 |
| corticosterone | 37.10679447 | 36.83996258 | 0.266831894 | 5500024024213121906563.H | .H07.G08.E09.D10.B11.A12 |
| tanespimycin | 37.57965012 | 37.31290545 | 0.266744666 | 5500024035736031208612.H | .G08.E09.D10.B11.A12 |
| (-)-atenolol | 37.50965017 | 37.24471367 | 0.264936498 | 5500024030700072107987.H | .H01.G02.E03.D04.B05.A06 |
| sulfamethoxypyridazi | 34.22504775 | 33.96129529 | 0.263752458 | 5500024025833041107269.G | .H07.E09.D10.B11.A12 |
| apomorphine | 35.6908308 | 35.4281341 | 0.262696699 | 612613111306.H08 | .H07.G08.E09.D10.B11.A12 |
| phenindione | 35.37271634 | 35.11129025 | 0.261426088 | 640641112706.G05 | .H01.G02.E03.D04.B05 |
| nafcillin | 34.39250555 | 34.13330462 | 0.259200936 | 5500024030760072207028.H | .H01.G02.D04.B05.A06 |
| ozagrel | 34.43548655 | 34.17633596 | 0.259150596 | 5500024030760072207033.H | .H07.G08.E09.B11.A12 |
| tiletamine | 37.71830192 | 37.46016387 | 0.258138055 | 5500024030700072107987.B | .H07.G08.E09.D10.B11.A12 |
| mebhydrolin | 37.09672925 | 36.83996258 | 0.256766672 | 5500024024213121906563.C | .H07.G08.E09.D10.B11.A12 |
| gabexate | 34.96497148 | 34.70962788 | 0.255343594 | 5500024030760072207033.A | .H01.G02.E03.D04.B05.A06 |
| repaglinide | 37.50217788 | 37.24787327 | 0.254304604 | 5500024035736031208612.F | .H01.G02.E03.D04.B05.A06 |
| fluvastatin | 34.4644417 | 34.21031976 | 0.254121943 | 5500024030760072207028.G | .H07.G08.E09.D10.B11.A12 |
| biperiden | 35.40866001 | 35.15552717 | 0.253132844 | 5500024024214122006604.A | .H01.G02.E03.D04.A06 |
| meglumine | 37.49775843 | 37.24471367 | 0.253044755 | 5500024030700072107987.H | .H01.G02.E03.D04.B05.A06 |
| thiocolchicoside | 35.36229225 | 35.11129025 | 0.251002004 | 6.41E+12 | .H01.G02.E03.D04.B05 |
| nilutamide | 37.49407628 | 37.24471367 | 0.249362604 | 5500024030700072107987.A | .H01.G02.E03.D04.B05.A06 |
| nifuroxazide | 35.38091959 | 35.13162255 | 0.249297034 | 5500024024214122006604.C | .H07.G08.E09.D10.B11.A12 |
| dorzolamide | 37.49643967 | 37.24787327 | 0.248566397 | 5500024035736031208612.D | .H01.G02.E03.D04.B05.A06 |
| guanabenz | 35.67656073 | 35.4281341 | 0.24842663 | 612613111306.A10 | .H07.G08.E09.D10.B11.A12 |
| prenylamine | 35.35950161 | 35.11129025 | 0.248211364 | 640641112706.C02 | .H01.G02.E03.D04.B05 |
| nalbuphine | 34.99512733 | 34.74789333 | 0.247233999 | 6.15E+12 | .H01.G02.E03.D04.B05.A06 |
| nordihydroguaiaretic a | 37.55996232 | 37.31290545 | 0.247056864 | 5.50E+33 | .G08.E09.D10.B11.A12 |
| telenzepine | 33.24782114 | 33.00109826 | 0.246722879 | 5.50E+29 | .H07.G08.E09.D10.B11.A12 |
| sulfamerazine | 35.30451131 | 35.05833814 | 0.246173167 | 5500024024213121906559.F | .H07.G08.E09.D10.B11.A12 |
| sulpiride | 35.51249626 | 35.2669347 | 0.245561553 | 612613111306.H06 | .H01.G02.E03.D04.B05.A06 |
| lactobionic acid | 37.08548915 | 36.83996258 | 0.245526569 | 5500024024213121906563.H | .H07.G08.E09.D10.B11.A12 |
| phenazopyridine | 34.20584883 | 33.96076648 | 0.245082355 | 5500024025833041107269.B | .H01.G02.E03.D04.B05.A06 |
| LY-294002 | 36.38678801 | 36.14209273 | 0.244695284 | 5500024017802120306174.B | .H01.G02.E03.D04.B05.A06 |
| rifampicin | 35.37556847 | 35.13162255 | 0.243945912 | 5500024024214122006604.D | .H07.G08.E09.D10.B11.A12 |
| streptomycin | 33.69663781 | 33.4530122 | 0.243625606 | 622623112706.B03 | .H01.G02.E03.B05.A06 |
| sulfadiazine | 37.8817727 | 37.63864853 | 0.243124173 | 5.50E+32 | .H07.G08.D10.B11.A12 |
| atropine methonitrate | 37.70249953 | 37.46016387 | 0.242335662 | 5500024030700072107987.F | .H07.G08.E09.D10.B11.A12 |
| salsolidin | 35.37385066 | 35.13162255 | 0.242228105 | 5500024024214122006604.H | .H07.G08.E09.D10.B11.A12 |
| phenelzine | 33.70024745 | 33.45935203 | 0.240895423 | 5500024024213121906560.C | .H01.G02.E03.D04.B05.A06 |
| cinchonine | 35.26081057 | 35.01997127 | 0.2408393 | 5500024024213121906559.G | .H01.G02.E03.D04.B05.A06 |
| adiphenine | 37.87886409 | 37.63864853 | 0.240215561 | 5500024024213121906564.A | .H07.G08.D10.B11.A12 |
| chloroquine | 35.35127037 | 35.11129025 | 0.239980115 | 640641112706.G06 | .H01.G02.E03.D04.B05 |
| crotamiton | 34.44908007 | 34.21031976 | 0.238760315 | 5500024030760072207028.C | .H07.G08.E09.D10.B11.A12 |
| mebeverine | 33.68595497 | 33.4530122 | 0.23294277 | 622623112706.B01 | .H01.G02.E03.B05.A06 |
| naproxen | 37.87145279 | 37.63864853 | 0.232804264 | 5500024024213121906564.A | .H07.G08.D10.B11.A12 |
| metamizole sodium | 35.28874956 | 35.05833814 | 0.230411414 | 5500024024213121906559.G | .H07.G08.E09.D10.B11.A12 |
| prazosin | 37.47495277 | 37.24471367 | 0.230239095 | 5500024030700072107987.B | .H01.G02.E03.D04.B05.A06 |
| idoxuridine | 35.49565291 | 35.2669347 | 0.228718204 | 6.13E+13 | .H01.G02.E03.D04.B05.A06 |
| ticlopidine | 35.49265798 | 35.2669347 | 0.225723274 | 612613111306.F03 | .H01.G02.E03.D04.B05.A06 |
| eticlopride | 34.4359716 | 34.21031976 | 0.225651848 | 5500024030760072207028.B | .H07.G08.E09.D10.B11.A12 |
| pimethixene | 33.2253343 | 33.00109826 | 0.224236038 | 5500024024213121906560.D | .H07.G08.E09.D10.B11.A12 |
| rimexolone | 34.4000052 | 34.17633596 | 0.223669247 | 5500024030760072207033.F | .H07.G08.E09.B11.A12 |
| nortriptyline | 33.22289526 | 33.00109826 | 0.221796998 | 5.50E+33 | .H07.G08.E09.D10.B11.A12 |
| nicotinic acid | 34.43187463 | 34.21031976 | 0.221554876 | 5.50E+32 | .H07.G08.E09.D10.B11.A12 |
| chloropyramine | 34.35440311 | 34.13330462 | 0.221098494 | 5500024030760072207028.C | .H01.G02.D04.B05.A06 |
| 1,4-chrysenequinone | 35.30723833 | 35.08636713 | 0.220871207 | 640641112706.D09 | .H07.G08.E09.D10.B11.A12 |
| cyproterone | 37.46836059 | 37.24787327 | 0.220487312 | 5500024035736031208612.H | .H01.G02.E03.D04.B05.A06 |
| gefitinib | 35.38197686 | 35.1615202 | 0.220456653 | EC2004073015AA | EC2004073014AA |
| naphazoline | 35.48724765 | 35.2669347 | 0.22031295 | 612613111306.H05 | .H01.G02.E03.D04.B05.A06 |
| amitriptyline | 37.85645485 | 37.63864853 | 0.217806318 | 5500024024213121906564.B | .H07.G08.D10.B11.A12 |
| homatropine | 37.85376647 | 37.63864853 | 0.215117938 | 5500024024213121906564.F | .H07.G08.D10.B11.A12 |
| lysergol | 37.05377726 | 36.83996258 | 0.213814681 | 5.50E+33 | .H07.G08.E09.D10.B11.A12 |
| acetylsalicylsalicylic a | 34.96048962 | 34.74789333 | 0.212596294 | 614615111406.F05 | .H01.G02.E03.D04.B05.A06 |
| Prestwick-972 | 37.67202658 | 37.46016387 | 0.211862711 | 5500024030700072107987.C | .H07.G08.E09.D10.B11.A12 |
| asiaticoside | 34.38800817 | 34.17633596 | 0.211672216 | 5500024030760072207033.H | .H07.G08.E09.B11.A12 |

| | | | | | |
|---|---|---|---|---|---|
| fludrocortisone | 33.67087347 | 33.45935203 | 0.211521441 | 5500024024213121906560.A | .H01.G02.E03.D04.B05.A06 |
| phenacetin | 35.34239372 | 35.13162255 | 0.210771164 | 5500024024214122006604.G | .H07.G08.E09.D10.B11.A12 |
| LY-294002 | 36.35282016 | 36.14209273 | 0.210727432 | 5500024017802120306174.F | .H01.G02.E03.D04.B05.A06 |
| fulvestrant | 34.63037571 | 34.42118584 | 0.209189872 | 5500024030401071707292.H | .H01.G02.E03.D04.B05.A06 |
| flurbiprofen | 37.45382797 | 37.24471367 | 0.209114301 | 5500024030700072107987.C | .H01.G02.E03.D04.B05.A06 |
| zuclopenthixol | 34.91866578 | 34.70962788 | 0.209037897 | 5500024030760072207033.B | .H01.G02.E03.D04.B05.A06 |
| vancomycin | 35.34038123 | 35.13162255 | 0.208758679 | 5500024024214122006604.A | .H07.G08.E09.D10.B11.A12 |
| tanespimycin | 34.6271963 | 34.42118584 | 0.206010466 | 5500024030401071707292.D | .H01.G02.E03.D04.B05.A06 |
| Prestwick-920 | 37.66530751 | 37.46016387 | 0.205143644 | 5500024030700072107987.F | .H07.G08.E09.D10.B11.A12 |
| alverine | 35.31356979 | 35.10850208 | 0.205067718 | 614615111406.D08 | .H07.G08.E09.D10.B11.A12 |
| vanoxerine | 33.66257021 | 33.45797002 | 0.204600195 | 622623112706.A09 | .H07.G08.E09.D10.B11.A12 |
| dosulepin | 35.31541492 | 35.11129025 | 0.204124666 | 640641112706.H05 | .H01.G02.E03.D04.B05 |
| thiamphenicol | 37.84009099 | 37.63864853 | 0.201442462 | 5500024024213121906564.B | .H07.G08.D10.B11.A12 |
| cefalotin | 34.16145245 | 33.96076648 | 0.20068597 | 5500024025833041107269.F | .H01.G02.E03.D04.B05.A06 |
| isometheptene | 37.65996372 | 37.46016387 | 0.199799847 | 5500024030700072107987.A | .H07.G08.E09.D10.B11.A12 |
| penbutolol | 34.37580294 | 34.17633596 | 0.199466985 | 5500024030760072207033.B | .H07.G08.E09.B11.A12 |
| (-)-MK-801 | 37.44322108 | 37.24471367 | 0.19850741 | 5.50E+22 | .H01.G02.E03.D04.B05.A06 |
| cortisone | 33.19911869 | 33.00109826 | 0.198020424 | 5500024024213121906560.F | .H07.G08.E09.D10.B11.A12 |
| pimozide | 33.65008572 | 33.4530122 | 0.197073519 | 6.23E+12 | .H01.G02.E03.B05.A06 |
| iproniazid | 35.30446001 | 35.10850208 | 0.195957938 | 614615111406.A08 | .H07.G08.E09.D10.B11.A12 |
| pirlindole | 37.65535527 | 37.46016387 | 0.195191401 | 5500024030700072107987.B | .H07.G08.E09.D10.B11.A12 |
| pirenperone | 35.3491142 | 35.15552717 | 0.193587029 | 5500024024214122006604.B | .H01.G02.E03.D04.A06 |
| haloperidol | 37.50456293 | 37.31290545 | 0.191657482 | 5500024035736031208612.A | .G08.E09.D10.B11.A12 |
| omeprazole | 35.32199018 | 35.13162255 | 0.190367624 | 5500024024214122006604.G | .H07.G08.E09.D10.B11.A12 |
| chlorpromazine | 37.82868968 | 37.63864853 | 0.190041152 | 5500024024213121906564.C | .H07.G08.D10.B11.A12 |
| haloperidol | 35.61454297 | 35.4281341 | 0.186408876 | 612613111306.B09 | .H07.G08.E09.D10.B11.A12 |
| doxepin | 33.18694792 | 33.00109826 | 0.185849652 | 5500024024213121906560.F | .H07.G08.E09.D10.B11.A12 |
| clozapine | 37.49816332 | 37.31290545 | 0.185257863 | 5500024035736031208612.C | .G08.E09.D10.B11.A12 |
| flumequine | 35.29372231 | 35.10850208 | 0.185220233 | 614615111406.D12 | .H07.G08.E09.D10.B11.A12 |
| betamethasone | 33.64227234 | 33.45797002 | 0.184302326 | 622623112706.H12 | .H07.G08.E09.D10.B11.A12 |
| caffeic acid | 34.39428805 | 34.21031976 | 0.183968298 | 5500024030760072207028.C | .H07.G08.E09.D10.B11.A12 |
| neostigmine bromide | 35.33529062 | 35.15552717 | 0.179763457 | 5500024024214122006604.F | .H01.G02.E03.D04.A06 |
| dicloxacillin | 35.33444155 | 35.15552717 | 0.178914383 | 5500024024214122006604.D | .H01.G02.E03.D04.A06 |
| diphemanil metilsulfa | 35.44499039 | 35.2669347 | 0.178055689 | 612613111306.C06 | .H01.G02.E03.D04.B05.A06 |
| cefixime | 37.0136016 | 36.83996258 | 0.173639025 | 5500024024213121906563.G | .H07.G08.E09.D10.B11.A12 |
| trichostatin A | 35.32909794 | 35.15552717 | 0.173570774 | 5500024024214122006604.C | .H01.G02.E03.D04.A06 |
| ceforanide | 35.32831328 | 35.15552717 | 0.172786112 | 5500024024214122006604.C | .H01.G02.E03.D04.A06 |
| gelsemine | 35.191971 | 35.01997127 | 0.171999731 | 5500024024213121906559.B | .H01.G02.E03.D04.B05.A06 |
| ifenprodil | 33.17304696 | 33.00109826 | 0.171948699 | 5500024024213121906560.H | .H07.G08.E09.D10.B11.A12 |
| velnacrine | 35.32743767 | 35.15552717 | 0.171910507 | 5500024024214122006604.G | .H01.G02.E03.D04.A06 |
| ergocalciferol | 35.32726004 | 35.15552717 | 0.171732876 | 5500024024214122006604.D | .H01.G02.E03.D04.A06 |
| nimesulide | 35.2797117 | 35.10850208 | 0.171209629 | 614615111406.D11 | .H07.G08.E09.D10.B11.A12 |
| aminophenazone | 34.91892465 | 34.74789333 | 0.171031318 | 614615111406.F04 | .H01.G02.E03.D04.B05.A06 |
| Prestwick-692 | 35.19086475 | 35.01997127 | 0.170893485 | 5500024024213121906559.A | .H01.G02.E03.D04.B05.A06 |
| geldanamycin | 34.59165827 | 34.42118584 | 0.170472432 | 5500024030401071707292.D | .H01.G02.E03.D04.B05.A06 |
| diphenylpyraline | 35.22834716 | 35.05833814 | 0.170009017 | 5500024024213121906559.A | .H07.G08.E09.D10.B11.A12 |
| letrozole | 34.87780351 | 34.70962788 | 0.168175627 | 5500024030760072207033.F | .H01.G02.E03.D04.B05.A06 |
| hemicholinium | 33.62411671 | 33.45797002 | 0.166146693 | 622623112706.F12 | .H07.G08.E09.D10.B11.A12 |
| proxymetacaine | 37.62339443 | 37.46016387 | 0.163230559 | 5500024030700072107987.G | .H07.G08.E09.D10.B11.A12 |
| flunisolide | 35.22076144 | 35.05833814 | 0.162423298 | 5500024024213121906559.H | .H07.G08.E09.D10.B11.A12 |
| trimethoprim | 33.62156699 | 33.45935203 | 0.162214957 | 5500024024213121906560.F | .H01.G02.E03.D04.B05.A06 |
| guanadrel | 34.12119464 | 33.96129529 | 0.159899355 | 5500024025833041107269.B | .H07.E09.D10.B11.A12 |
| ribavirin | 37.61922767 | 37.46016387 | 0.159063801 | 5500024030700072107987.A | .H07.G08.E09.D10.B11.A12 |
| cyclopentolate | 37.4063002 | 37.24787327 | 0.158426928 | 5500024035736031208612.F | .H01.G02.E03.D04.B05.A06 |
| betonicine | 35.21614924 | 35.05833814 | 0.157811101 | 5500024024213121906559.A | .H07.G08.E09.D10.B11.A12 |
| spironolactone | 34.90484882 | 34.74789333 | 0.156955489 | 6.15E+13 | .H01.G02.E03.D04.B05.A06 |
| trichostatin A | 34.55647422 | 34.40088765 | 0.155586574 | 5.50E+31 | .H07.G08.E09.D10.B11 |
| trichostatin A | 34.28854502 | 34.13330462 | 0.1552404 | 5500024030760072207028.F | .H01.G02.D04.B05.A06 |
| valproic acid | 36.29597245 | 36.14209273 | 0.153879722 | 5.50E+26 | .H01.G02.E03.D04.B05.A06 |
| etomidate | 34.33019335 | 34.17633596 | 0.153857393 | 5.50E+28 | .H07.G08.E09.B11.A12 |
| diphenhydramine | 37.79175257 | 37.63864853 | 0.153104043 | 5500024024213121906564.A | .H07.G08.D10.B11.A12 |
| florfenicol | 37.39565091 | 37.24471367 | 0.150937238 | 5.50E+25 | .H01.G02.E03.D04.B05.A06 |
| esculin | 34.36093462 | 34.21031976 | 0.150614868 | 5500024030760072207028.C | .H07.G08.E09.D10.B11.A12 |
| trifluridine | 37.39711816 | 37.24787327 | 0.149244882 | 5500024035736031208612.F | .H01.G02.E03.D04.B05.A06 |
| boldine | 35.16812124 | 35.01997127 | 0.148149967 | 5500024024213121906559.D | .H01.G02.E03.D04.B05.A06 |
| ceftazidime | 35.25894552 | 35.11129025 | 0.147655272 | 640641112706.F02 | .H01.G02.E03.D04.B05 |
| bretylium tosilate | 34.35499389 | 34.21031976 | 0.14467413 | 5500024030760072207028.B | .H07.G08.E09.D10.B11.A12 |
| carbimazole | 35.29987795 | 35.15552717 | 0.144350787 | 5.50E+22 | .H01.G02.E03.D04.A06 |
| haloperidol | 34.56547052 | 34.42118584 | 0.144284679 | 5500024030401071707292.H | .H01.G02.E03.D04.B05.A06 |

| | | | | | |
|---|---|---|---|---|---|
| calcium folinate | 34.10485477 | 33.96129529 | 0.143559483 | 5500024025833041107269.A | .H07.E09.D10.B11.A12 |
| methoxamine | 35.27497712 | 35.13162255 | 0.143354563 | 5500024024214122006604.C | .H07.G08.E09.D10.B11.A12 |
| pinacidil | 33.14407107 | 33.00109826 | 0.14297281 | 5500024024213121906560.B | .H07.G08.E09.D10.B11.A12 |
| benzocaine | 35.16145769 | 35.01997127 | 0.141486416 | 5500024024213121906559.A | .H01.G02.E03.D04.B05.A06 |
| suxibuzone | 34.10130452 | 33.96076648 | 0.140538038 | 5500024025833041107269.H | .H01.G02.E03.D04.B05.A06 |
| octopamine | 37.60009602 | 37.46016387 | 0.139932148 | 5500024030700072107987.G | .H07.G08.E09.D10.B11.A12 |
| chlorhexidine | 35.56573583 | 35.4281341 | 0.13760173 | 6.13E+23 | .H07.G08.E09.D10.B11.A12 |
| levocabastine | 34.31311256 | 34.17633596 | 0.136776599 | 5500024030760072207033.G | .H07.G08.E09.B11.A12 |
| alfadolone | 37.59569651 | 37.46016387 | 0.135532641 | 5500024030700072107987.D | .H07.G08.E09.D10.B11.A12 |
| butoconazole | 35.29027959 | 35.15552717 | 0.134752426 | 5500024024214122006604.G | .H01.G02.E03.D04.A06 |
| azapropazone | 37.59488945 | 37.46016387 | 0.13472558 | 5500024030700072107987.A | .H07.G08.E09.D10.B11.A12 |
| metampicillin | 35.2427841 | 35.10850208 | 0.134282026 | 614615111406.B12 | .H07.G08.E09.D10.B11.A12 |
| amoxapine | 35.56222063 | 35.4281341 | 0.13408653 | 612613111306.G11 | .H07.G08.E09.D10.B11.A12 |
| methyldopate | 34.84322702 | 34.70962788 | 0.133599139 | 5500024030760072207033.A | .H01.G02.E03.D04.B05.A06 |
| triprolidine | 33.13401685 | 33.00109826 | 0.132918582 | 5500024024213121906560.G | .H07.G08.E09.D10.B11.A12 |
| bupivacaine | 33.1339344 | 33.00109826 | 0.132836132 | 5500024024213121906560.B | .H07.G08.E09.D10.B11.A12 |
| acetylsalicylic acid | 34.55326048 | 34.42118584 | 0.132074641 | 5500024030401071707292.H | .H01.G02.E03.D04.B05.A06 |
| ciprofibrate | 37.37945946 | 37.24787327 | 0.131586188 | 5.50E+23 | .H01.G02.E03.D04.B05.A06 |
| budesonide | 35.24233208 | 35.11129025 | 0.131041832 | 640641112706.G03 | .H01.G02.E03.D04.B05 |
| calcium pantothenate | 36.97081228 | 36.83996258 | 0.130849701 | 5500024024213121906563.G | .H07.G08.E09.D10.B11.A12 |
| tranexamic acid | 34.87841711 | 34.74789333 | 0.130523782 | 614615111406.A03 | .H01.G02.E03.D04.B05.A06 |
| clemizole | 33.58757822 | 33.45935203 | 0.128226191 | 5500024024213121906560.G | .H01.G02.E03.D04.B05.A06 |
| nicergoline | 34.87552111 | 34.74789333 | 0.127627781 | 614615111406.F02 | .H01.G02.E03.D04.B05.A06 |
| gabapentin | 34.52829819 | 34.40088765 | 0.127410542 | 5500024030401071707292.C | .H07.G08.E09.D10.B11 |
| remoxipride | 37.58718324 | 37.46016387 | 0.12701937 | 5.50E+32 | .H07.G08.E09.D10.B11.A12 |
| bupropion | 33.57951553 | 33.4530122 | 0.126503333 | 6.23E+15 | .H01.G02.E03.B05.A06 |
| cefepime | 37.37425002 | 37.24787327 | 0.126376746 | 5500024035736031208612.A | .H01.G02.E03.D04.B05.A06 |
| ciclopirox | 35.28082212 | 35.15552717 | 0.125294949 | 5500024024214122006604.B | .H01.G02.E03.D04.A06 |
| fulvestrant | 34.54647456 | 34.42118584 | 0.125288728 | 5500024030401071707292.B | .H01.G02.E03.D04.B05.A06 |
| valproic acid | 37.43784206 | 37.31290545 | 0.12493661 | 5500024035736031208612.G | .G08.E09.D10.B11.A12 |
| Prestwick-642 | 35.14466333 | 35.01997127 | 0.124692056 | 5500024024213121906559.B | .H01.G02.E03.D04.B05.A06 |
| chloropyrazine | 37.3717267 | 37.24787327 | 0.123853429 | 5500024035736031208612.C | .H01.G02.E03.D04.B05.A06 |
| gliclazide | 35.2349668 | 35.11129025 | 0.123676548 | 640641112706.F01 | .H01.G02.E03.D04.B05 |
| naproxen | 34.08387316 | 33.96076648 | 0.123106683 | 5500024025833041107269.C | .H01.G02.E03.D04.B05.A06 |
| haloperidol | 36.26513979 | 36.14209273 | 0.123047063 | 5500024017802120306174.A | .H01.G02.E03.D04.B05.A06 |
| probenecid | 35.25335337 | 35.13162255 | 0.121730816 | 5500024024214122006604.H | .H07.G08.E09.D10.B11.A12 |
| nalidixic acid | 33.58103914 | 33.45935203 | 0.121687107 | 5500024024213121906560.G | .H01.G02.E03.D04.B05.A06 |
| LY-294002 | 36.26327087 | 36.14209273 | 0.121178143 | 5500024017802120306174.B | .H01.G02.E03.D04.B05.A06 |
| molsidomine | 35.23179249 | 35.11129025 | 0.120502237 | 640641112706.H03 | .H01.G02.E03.D04.B05 |
| sirolimus | 34.54055646 | 34.42118584 | 0.11937062 | 5500024030401071707292.H | .H01.G02.E03.D04.B05.A06 |
| papaverine | 35.20540841 | 35.08636713 | 0.119041286 | 640641112706.H12 | .H07.G08.E09.D10.B11.A12 |
| amikacin | 33.57646286 | 33.45797002 | 0.118492842 | 622623112706.B07 | .H07.G08.E09.D10.B11.A12 |
| aminoglutethimide | 33.11797238 | 33.00109826 | 0.116874119 | 5.50E+32 | .H07.G08.E09.D10.B11.A12 |
| 7-aminocephalospora | 36.95645273 | 36.83996258 | 0.116490156 | 5.50E+29 | .H07.G08.E09.D10.B11.A12 |
| thiamine | 35.22758925 | 35.11129025 | 0.116299003 | 640641112706.B04 | .H01.G02.E03.D04.B05 |
| amoxicillin | 37.12414874 | 37.00786033 | 0.116288412 | 5500024024213121906563.H | .H01.G02.E03.D04.B05.A06 |
| diethylstilbestrol | 34.07709867 | 33.96129529 | 0.115803379 | 5500024025833041107269.C | .H07.E09.D10.B11.A12 |
| trichostatin A | 34.07443744 | 33.96076648 | 0.113670964 | 5.50E+27 | .H01.G02.E03.D04.B05.A06 |
| pindolol | 34.86125417 | 34.74789333 | 0.113360845 | 614615111406.C04 | .H01.G02.E03.D04.B05.A06 |
| dyclonine | 33.11400381 | 33.00109826 | 0.112905544 | 5500024024213121906560.D | .H07.G08.E09.D10.B11.A12 |
| methotrexate | 33.57077372 | 33.45797002 | 0.112803706 | 622623112706.F10 | .H07.G08.E09.D10.B11.A12 |
| demecarium bromide | 35.198891 | 35.08636713 | 0.112523868 | 640641112706.C12 | .H07.G08.E09.D10.B11.A12 |
| citiolone | 35.16930027 | 35.05833814 | 0.110962129 | 5500024024213121906559.G | .H07.G08.E09.D10.B11.A12 |
| flutamide | 33.57001214 | 33.45935203 | 0.110660113 | 5500024024213121906560.C | .H01.G02.E03.D04.B05.A06 |
| fluphenazine | 33.56365079 | 33.4530122 | 0.110638585 | 622623112706.B02 | .H01.G02.E03.B05.A06 |
| ramipril | 37.35704908 | 37.24787327 | 0.109175806 | 5500024035736031208612.C | .H01.G02.E03.D04.B05.A06 |
| phensuximide | 34.28502753 | 34.17633596 | 0.10869157 | 5.50E+31 | .H07.G08.E09.B11.A12 |
| fosfosal | 34.24033954 | 34.13330462 | 0.107034921 | 5500024030760072207028.F | .H01.G02.D04.B05.A06 |
| estriol | 37.35415371 | 37.24787327 | 0.106280439 | 5.50E+26 | .H01.G02.E03.D04.B05.A06 |
| norethisterone | 33.10687841 | 33.00109826 | 0.105780148 | 5500024024213121906560.F | .H07.G08.E09.D10.B11.A12 |
| cyclic adenosine mon | 34.28206625 | 34.17633596 | 0.105730294 | 5500024030760072207033.C | .H07.G08.E09.B11.A12 |
| oxybuprocaine | 35.37194697 | 35.2669347 | 0.105012268 | 612613111306.F04 | .H01.G02.E03.D04.B05.A06 |
| isopropamide iodide | 34.5049957 | 34.40088765 | 0.104108049 | 5.50E+29 | .H07.G08.E09.D10.B11 |
| guanethidine | 33.5569603 | 33.4530122 | 0.103948101 | 622623112706.G05 | .H01.G02.E03.B05.A06 |
| alprenolol | 33.55595665 | 33.4530122 | 0.102944444 | 622623112706.C01 | .H01.G02.E03.B05.A06 |
| Prestwick-981 | 37.56091205 | 37.46016387 | 0.100748183 | 5.50E+33 | .H07.G08.E09.D10.B11.A12 |
| glycocholic acid | 35.25595058 | 35.15552717 | 0.100423417 | 5500024024214122006604.B | .H01.G02.E03.D04.A06 |
| cytisine | 35.18667621 | 35.08636713 | 0.100309077 | 6.41E+18 | .H07.G08.E09.D10.B11.A12 |
| nizatidine | 34.31061417 | 34.21031976 | 0.10029441 | 5500024030760072207028.D | .H07.G08.E09.D10.B11.A12 |

| | | | | | |
|---|---|---|---|---|---|
| raloxifene | 34.50058761 | 34.40088765 | 0.099699962 | 5500024030401071707292.B | .H07.G08.E09.D10.B11 |
| clozapine | 34.52086319 | 34.42118584 | 0.099677353 | 5500024030401071707292.C | .H01.G02.E03.D04.B05.A06 |
| naloxone | 35.52639671 | 35.4281341 | 0.098262613 | 612613111306.H09 | .H07.G08.E09.D10.B11.A12 |
| thioperamide | 34.30844147 | 34.21031976 | 0.098121714 | 5500024030760072207028.C | .H07.G08.E09.D10.B11.A12 |
| methapyrilene | 33.55537315 | 33.45797002 | 0.097403138 | 622623112706.H10 | .H07.G08.E09.D10.B11.A12 |
| levamisole | 35.20435663 | 35.10850208 | 0.095854557 | 614615111406.G09 | .H07.G08.E09.D10.B11.A12 |
| piracetam | 35.20712406 | 35.11129025 | 0.09583381 | 640641112706.H02 | .H01.G02.E03.D04.B05 |
| adipiodone | 37.55590979 | 37.46016387 | 0.09574592 | 5500024030700072107987.G | .H07.G08.E09.D10.B11.A12 |
| mepyramine | 33.54775313 | 33.4530122 | 0.094740926 | 622623112706.D02 | .H01.G02.E03.B05.A06 |
| bufexamac | 33.09504419 | 33.00109826 | 0.093945923 | 5500024024213121906560.F | .H07.G08.E09.D10.B11.A12 |
| hexamethonium brom | 35.3607717 | 35.2669347 | 0.093836997 | 6.13E+16 | .H01.G02.E03.D04.B05.A06 |
| foliosidine | 35.15156886 | 35.05833814 | 0.093230716 | 5500024024213121906559.B | .H07.G08.E09.D10.B11.A12 |
| diflunisal | 35.3581299 | 35.2669347 | 0.091195192 | 612613111306.C02 | .H01.G02.E03.D04.B05.A06 |
| metyrapone | 37.3344943 | 37.24471367 | 0.089780634 | 5500024030700072107987.H | .H01.G02.E03.D04.B05.A06 |
| flufenamic acid | 35.19827584 | 35.10850208 | 0.089773763 | 6.15E+18 | .H07.G08.E09.D10.B11.A12 |
| hexylcaine | 34.48977163 | 34.40088765 | 0.088883986 | 5500024030401071707292.H | .H07.G08.E09.D10.B11 |
| mafenide | 35.1957671 | 35.10850208 | 0.087265022 | 614615111406.A07 | .H07.G08.E09.D10.B11.A12 |
| rosiglitazone | 36.2289294 | 36.14209273 | 0.086836667 | 5500024017802120306174.C | .H01.G02.E03.D04.B05.A06 |
| ioversol | 34.29648343 | 34.21031976 | 0.086163671 | 5500024030760072207028.H | .H07.G08.E09.D10.B11.A12 |
| domperidone | 37.09343714 | 37.00786033 | 0.08557681 | 5500024024213121906563.A | .H01.G02.E03.D04.B05.A06 |
| cefoxitin | 34.4862331 | 34.40088765 | 0.085345453 | 5500024030401071707292.B | .H07.G08.E09.D10.B11 |
| chlorpropamide | 33.54325183 | 33.45797002 | 0.085281812 | 622623112706.G11 | .H07.G08.E09.D10.B11.A12 |
| levobunolol | 34.21799585 | 34.13330462 | 0.084691236 | 5500024030760072207028.B | .H01.G02.D04.B05.A06 |
| hecogenin | 34.48272339 | 34.40088765 | 0.081835739 | 5500024030401071707292.F | .H07.G08.E09.D10.B11 |
| reserpine | 34.21511046 | 34.13330462 | 0.081805839 | 5.50E+27 | .H01.G02.D04.B05.A06 |
| 4-hydroxyphenazone | 35.34601754 | 35.2669347 | 0.079082838 | 612613111306.B03 | .H01.G02.E03.D04.B05.A06 |
| xamoterol | 34.28917651 | 34.21031976 | 0.078856756 | 5500024030760072207028.A | .H07.G08.E09.D10.B11.A12 |
| clozapine | 37.08636984 | 37.00786033 | 0.078509508 | 5500024024213121906563.C | .H01.G02.E03.D04.B05.A06 |
| acetazolamide | 37.7164076 | 37.63864853 | 0.077759067 | 5.50E+29 | .H07.G08.D10.B11.A12 |
| cotinine | 35.5053473 | 35.4281341 | 0.077213199 | 612613111306.G09 | .H07.G08.E09.D10.B11.A12 |
| oxymetazoline | 35.18318199 | 35.10850208 | 0.074679917 | 614615111406.C08 | .H07.G08.E09.D10.B11.A12 |
| harmaline | 35.09339326 | 35.01997127 | 0.073421994 | 5500024024213121906559.D | .H01.G02.E03.D04.B05.A06 |
| acenocoumarol | 34.82067277 | 34.74789333 | 0.072779438 | 614615111406.C06 | .H01.G02.E03.D04.B05.A06 |
| carteolol | 36.9097886 | 36.83996258 | 0.06982602 | 5500024024213121906563.B | .H07.G08.E09.D10.B11.A12 |
| vinpocetine | 33.52274826 | 33.4530122 | 0.069736059 | 622623112706.F02 | .H01.G02.E03.B05.A06 |
| tiaprofenic acid | 35.20037754 | 35.13162255 | 0.068754985 | 5500024024214122006604.C | .H07.G08.E09.D10.B11.A12 |
| isradipine | 37.52832101 | 37.46016387 | 0.068157137 | 5500024030700072107987.D | .H07.G08.E09.D10.B11.A12 |
| iodixanol | 34.20099881 | 34.13330462 | 0.067694188 | 5500024030760072207028.A | .H01.G02.D04.B05.A06 |
| tenoxicam | 35.19931002 | 35.13162255 | 0.067687462 | 5500024024214122006604.A | .H07.G08.E09.D10.B11.A12 |
| bumetanide | 33.06875322 | 33.00109826 | 0.067654956 | 5500024024213121906560.A | .H07.G08.E09.D10.B11.A12 |
| trimipramine | 34.20072306 | 34.13330462 | 0.067418443 | 5500024030760072207028.D | .H01.G02.D04.B05.A06 |
| alfaxalone | 37.52675714 | 37.46016387 | 0.066593273 | 5500024030700072107987.C | .H07.G08.E09.D10.B11.A12 |
| loxapine | 35.49463248 | 35.4281341 | 0.066498386 | 612613111306.F08 | .H07.G08.E09.D10.B11.A12 |
| pronetalol | 34.77609581 | 34.70962788 | 0.066467933 | 5500024030760072207033.H | .H01.G02.E03.D04.B05.A06 |
| verapamil | 35.49455638 | 35.4281341 | 0.066422279 | 612613111306.H12 | .H07.G08.E09.D10.B11.A12 |
| trichostatin A | 33.52442624 | 33.45935203 | 0.065074213 | 5500024024213121906560.A | .H01.G02.E03.D04.B05.A06 |
| retrorsine | 35.08494225 | 35.01997127 | 0.064970979 | 5500024024213121906559.H | .H01.G02.E03.D04.B05.A06 |
| amylocaine | 35.3277281 | 35.2669347 | 0.060793392 | 612613111306.C03 | .H01.G02.E03.D04.B05.A06 |
| netilmicin | 34.23588078 | 34.17633596 | 0.05954482 | 5500024030760072207033.D | .H07.G08.E09.B11.A12 |
| sulfacetamide | 37.69791428 | 37.63864853 | 0.059265756 | 5500024024213121906564.C | .H07.G08.D10.B11.A12 |
| equilin | 34.26929567 | 34.21031976 | 0.058975919 | 5500024030760072207028.F | .H07.G08.E09.D10.B11.A12 |
| moracizine | 34.23456595 | 34.17633596 | 0.05822999 | 5.50E+29 | .H07.G08.E09.B11.A12 |
| metoclopramide | 33.5173708 | 33.45935203 | 0.058018773 | 5500024024213121906560.D | .H01.G02.E03.D04.B05.A06 |
| serotonin | 35.21320754 | 35.15552717 | 0.057680374 | 5500024024214122006604.C | .H01.G02.E03.D04.A06 |
| butyl hydroxybenzoat | 37.29992446 | 37.24471367 | 0.055210787 | 5500024030700072107987.H | .H01.G02.E03.D04.B05.A06 |
| minaprine | 35.3220152 | 35.2669347 | 0.055080495 | 612613111306.G01 | .H01.G02.E03.D04.B05.A06 |
| bezafibrate | 37.06177249 | 37.00786033 | 0.053912157 | 5500024024213121906563.F | .H01.G02.E03.D04.B05.A06 |
| folic acid | 35.140016 | 35.08636713 | 0.053648876 | 640641112706.A11 | .H07.G08.E09.D10.B11.A12 |
| sisomicin | 35.18359914 | 35.13162255 | 0.051976585 | 5500024024214122006604.C | .H07.G08.E09.D10.B11.A12 |
| Prestwick-682 | 35.07189667 | 35.01997127 | 0.051925403 | 5500024024213121906559.A | .H01.G02.E03.D04.B05.A06 |
| naftidrofuryl | 37.05951333 | 37.00786033 | 0.051653001 | 5500024024213121906563.H | .H01.G02.E03.D04.B05.A06 |
| mifepristone | 33.50335078 | 33.4530122 | 0.050338579 | 622623112706.D03 | .H01.G02.E03.B05.A06 |
| ketorolac | 37.51018283 | 37.46016387 | 0.050018964 | 5500024030700072107987.G | .H07.G08.E09.D10.B11.A12 |
| clebopride | 37.05174494 | 37.00786033 | 0.043884611 | 5500024024213121906563.C | .H01.G02.E03.D04.B05.A06 |
| triamterene | 37.6815796 | 37.63864853 | 0.042931074 | 5500024024213121906564.C | .H07.G08.D10.B11.A12 |
| trifluoperazine | 37.35502106 | 37.31290545 | 0.042115607 | 5500024035736031208612.D | .G08.E09.D10.B11.A12 |
| hydralazine | 33.49987518 | 33.45935203 | 0.040523155 | 5.50E+26 | .H01.G02.E03.D04.B05.A06 |
| metrifonate | 37.67890647 | 37.63864853 | 0.040257942 | 5500024024213121906564.G | .H07.G08.D10.B11.A12 |
| fluphenazine | 36.18229365 | 36.14209273 | 0.040200916 | 5500024017802120306174.B | .H01.G02.E03.D04.B05.A06 |

| | | | | | |
|---|---|---|---|---|---|
| isocorydine | 35.12653333 | 35.08636713 | 0.040166202 | 640641112706.A08 | .H07.G08.E09.D10.B11.A12 |
| myosmine | 35.09823982 | 35.05833814 | 0.039901677 | 5500024024213121906559.B | .H07.G08.E09.D10.B11.A12 |
| 3-acetylcoumarin | 34.24949016 | 34.21031976 | 0.039170404 | 5.50E+33 | .H07.G08.E09.D10.B11.A12 |
| ursodeoxycholic acid | 37.49922226 | 37.46016387 | 0.039058396 | 5500024030700072107987.H | .H07.G08.E09.D10.B11.A12 |
| trichostatin A | 35.09706913 | 35.05833814 | 0.03873099 | 5500024024213121906559.A | .H07.G08.E09.D10.B11.A12 |
| metolazone | 35.46472571 | 35.4281341 | 0.036591617 | 612613111306.G12 | .H07.G08.E09.D10.B11.A12 |
| haloperidol | 34.45755013 | 34.42118584 | 0.036364298 | 5500024030401071707292.A | .H01.G02.E03.D04.B05.A06 |
| capsaicin | 34.24484799 | 34.21031976 | 0.034528238 | 5500024030760072207028.F | .H07.G08.E09.D10.B11.A12 |
| estrone | 37.27777993 | 37.24471367 | 0.033066254 | 5500024030700072107987.G | .H01.G02.E03.D04.B05.A06 |
| fursultiamine | 34.74070665 | 34.70962788 | 0.031078766 | 5500024030760072207033.C | .H01.G02.E03.D04.B05.A06 |
| ondansetron | 37.27739586 | 37.24787327 | 0.029522582 | 5500024035736031208612.B | .H01.G02.E03.D04.B05.A06 |
| Prestwick-674 | 35.08619286 | 35.05833814 | 0.027854721 | 5500024024213121906559.F | .H07.G08.E09.D10.B11.A12 |
| zidovudine | 33.48362879 | 33.45797002 | 0.025658772 | 622623112706.G12 | .H07.G08.E09.D10.B11.A12 |
| tropine | 37.27268873 | 37.24787327 | 0.024815455 | 5500024035736031208612.C | .H01.G02.E03.D04.B05.A06 |
| primidone | 34.23501996 | 34.21031976 | 0.024700205 | 5500024030760072207028.A | .H07.G08.E09.D10.B11.A12 |
| lorglumide | 37.26870657 | 37.24471367 | 0.023992904 | 5500024030700072107987.F | .H01.G02.E03.D04.B05.A06 |
| 2-aminobenzenesulfo | 34.23399427 | 34.21031976 | 0.023674518 | 5500024030760072207028.A | .H07.G08.E09.D10.B11.A12 |
| fluorometholone | 33.98397241 | 33.96076648 | 0.023205937 | 5500024025833041107269.G | .H01.G02.E03.D04.B05.A06 |
| berberine | 35.10865807 | 35.08636713 | 0.022290941 | 640641112706.C09 | .H07.G08.E09.D10.B11.A12 |
| minoxidil | 35.2892172 | 35.2669347 | 0.022282493 | 612613111306.B02 | .H01.G02.E03.D04.B05.A06 |
| mevalolactone | 34.42315898 | 34.40088765 | 0.022271328 | 5500024030401071707292.F | .H07.G08.E09.D10.B11 |
| trimethylcolchicinic ac | 35.04210086 | 35.01997127 | 0.022129595 | 5.50E+26 | .H01.G02.E03.D04.B05.A06 |
| pepstatin | 36.86185788 | 36.83996258 | 0.021895302 | 5500024024213121906563.D | .H07.G08.E09.D10.B11.A12 |
| trichostatin A | 34.23190965 | 34.21031976 | 0.021589897 | 5500024030760072207028.B | .H07.G08.E09.D10.B11.A12 |
| methanthelinium bror | 37.2678029 | 37.24787327 | 0.019929624 | 5.50E+22 | .H01.G02.E03.D04.B05.A06 |
| cycloserine | 37.26720045 | 37.24787327 | 0.019327171 | 5.50E+25 | .H01.G02.E03.D04.B05.A06 |
| maprotiline | 33.47645352 | 33.45797002 | 0.018483506 | 622623112706.B10 | .H07.G08.E09.D10.B11.A12 |
| meclocycline | 36.85802787 | 36.83996258 | 0.018065291 | 5500024024213121906563.B | .H07.G08.E09.D10.B11.A12 |
| tridihexethyl | 34.19222238 | 34.17633596 | 0.015886422 | 5500024030760072207033.D | .H07.G08.E09.B11.A12 |
| erythromycin | 35.44302583 | 35.4281341 | 0.014891735 | 612613111306.G07 | .H07.G08.E09.D10.B11.A12 |
| proxyphylline | 37.47284953 | 37.46016387 | 0.012685659 | 5500024030700072107987.F | .H07.G08.E09.D10.B11.A12 |
| fluspirilene | 37.25736033 | 37.24471367 | 0.012646662 | 5500024030700072107987.D | .H01.G02.E03.D04.B05.A06 |
| pyrantel | 35.11995936 | 35.10850208 | 0.01145728 | 614615111406.G12 | .H07.G08.E09.D10.B11.A12 |
| betulinic acid | 36.85141925 | 36.83996258 | 0.011456674 | 5500024024213121906563.A | .H07.G08.E09.D10.B11.A12 |
| econazole | 33.01104367 | 33.00109826 | 0.009945407 | 5500024024213121906560.D | .H07.G08.E09.D10.B11.A12 |
| pentolonium | 33.46885093 | 33.45935203 | 0.0094989 | 5500024024213121906560.F | .H01.G02.E03.D04.B05.A06 |
| fusaric acid | 36.84931595 | 36.83996258 | 0.009353375 | 5500024024213121906563.H | .H07.G08.E09.D10.B11.A12 |
| biotin | 35.1638073 | 35.15552717 | 0.008280134 | 5500024024214122006604.G | .H01.G02.E03.D04.A06 |
| hesperidin | 37.01544172 | 37.00786033 | 0.007581389 | 5500024024213121906563.C | .H01.G02.E03.D04.B05.A06 |
| cefalonium | 34.71659351 | 34.70962788 | 0.006965625 | 5.50E+27 | .H01.G02.E03.D04.B05.A06 |
| ketoprofen | 33.46598399 | 33.45935203 | 0.006631958 | 5500024024213121906560.D | .H01.G02.E03.D04.B05.A06 |
| pregnenolone | 35.13590816 | 35.13162255 | 0.004285609 | 5500024024214122006604.B | .H07.G08.E09.D10.B11.A12 |
| heptaminol | 37.64242553 | 37.63864853 | 0.003777 | 5500024024213121906564.B | .H07.G08.D10.B11.A12 |
| chlortalidone | 33.45621549 | 33.4530122 | 0.003203284 | 622623112706.A01 | .H01.G02.E03.B05.A06 |
| Prestwick-664 | 35.06144086 | 35.05833814 | 0.003102718 | 5500024024213121906559.F | .H07.G08.E09.D10.B11.A12 |
| edrophonium chloride | 35.43018033 | 35.4281341 | 0.002046233 | 612613111306.F11 | .H07.G08.E09.D10.B11.A12 |
| trichostatin A | 34.71069717 | 34.70962788 | 0.001069294 | 5500024030760072207033.H | .H01.G02.E03.D04.B05.A06 |
| LY-294002 | 34.94387722 | 34.94387722 | 3.55E-14 | EC2004030503AA | EC2004030502AA |
| propidium iodide | 33.96059986 | 33.96076648 | -0.000166614 | 5500024025833041107269.A | .H01.G02.E03.D04.B05.A06 |
| profenamine | 34.20986437 | 34.21031976 | -0.000455381 | 5500024030760072207028.F | .H07.G08.E09.D10.B11.A12 |
| digoxigenin | 34.20845039 | 34.21031976 | -0.001869365 | 5500024030760072207028.B | .H07.G08.E09.D10.B11.A12 |
| LY-294002 | 34.41803608 | 34.42118584 | -0.003149758 | 5500024030401071707292.F | .H01.G02.E03.D04.B05.A06 |
| brinzolamide | 33.4536162 | 33.45797002 | -0.00435382 | 622623112706.C10 | .H07.G08.E09.D10.B11.A12 |
| finasteride | 35.0536549 | 35.05833814 | -0.004683239 | 5500024024213121906559.A | .H07.G08.E09.D10.B11.A12 |
| sirolimus | 34.41625502 | 34.42118584 | -0.004930816 | 5.50E+25 | .H01.G02.E03.D04.B05.A06 |
| ofloxacin | 33.45438203 | 33.45935203 | -0.004970003 | 5500024024213121906560.F | .H01.G02.E03.D04.B05.A06 |
| dexibuprofen | 37.23969399 | 37.24471367 | -0.005019679 | 5500024030700072107987.C | .H01.G02.E03.D04.B05.A06 |
| etofylline | 35.10324249 | 35.10850208 | -0.005259588 | 614615111406.G07 | .H07.G08.E09.D10.B11.A12 |
| flecainide | 33.95557119 | 33.96129529 | -0.005724103 | 5500024025833041107269.F | .H07.E09.D10.B11.A12 |
| pipenzolate bromide | 34.39485568 | 34.40088765 | -0.006031968 | 5.50E+28 | .H07.G08.E09.D10.B11 |
| altretamine | 37.23679683 | 37.24471367 | -0.007916842 | 5500024030700072107987.D | .H01.G02.E03.D04.B05.A06 |
| buspirone | 36.99934596 | 37.00786033 | -0.008514372 | 5.50E+26 | .H01.G02.E03.D04.B05.A06 |
| oleandomycin | 35.41934539 | 35.4281341 | -0.008788708 | 612613111306.F10 | .H07.G08.E09.D10.B11.A12 |
| bacampicillin | 36.83108699 | 36.83996258 | -0.008875591 | 5500024024213121906563.B | .H07.G08.E09.D10.B11.A12 |
| isoniazid | 34.73763968 | 34.74789333 | -0.010253653 | 614615111406.A01 | .H01.G02.E03.D04.B05.A06 |
| yohimbic acid | 35.00858615 | 35.01997127 | -0.011385123 | 5.50E+27 | .H01.G02.E03.D04.B05.A06 |
| iopamidol | 34.38930678 | 34.40088765 | -0.011580873 | 5500024030401071707292.C | .H07.G08.E09.D10.B11 |
| bergenin | 34.38777265 | 34.40088765 | -0.013115002 | 5500024030401071707292.D | .H07.G08.E09.D10.B11 |
| nadolol | 34.11942898 | 34.13330462 | -0.01387564 | 5500024030760072207028.A | .H01.G02.D04.B05.A06 |

| | | | | | |
|---|---|---|---|---|---|
| skimmianine | 33.94654789 | 33.96076648 | -0.014218588 | 5500024025833041107269.H | .H01.G02.E03.D04.B05.A06 |
| rosiglitazone | 37.29777222 | 37.31290545 | -0.015133235 | 5500024035736031208612.C | .G08.E09.D10.B11.A12 |
| tiapride | 33.44420536 | 33.45935203 | -0.015146669 | 5500024024213121906560.H | .H01.G02.E03.D04.B05.A06 |
| bromopride | 35.04153327 | 35.05833814 | -0.01680487 | 5500024024213121906559.F | .H07.G08.E09.D10.B11.A12 |
| kanamycin | 33.4397802 | 33.45797002 | -0.018189819 | 622623112706.D09 | .H07.G08.E09.D10.B11.A12 |
| procyclidine | 34.1146119 | 34.13330462 | -0.018692719 | 5500024030760072207028.G | .H01.G02.D04.B05.A06 |
| haloperidol | 37.29363558 | 37.31290545 | -0.019269868 | 5500024035736031208612.H | .G08.E09.D10.B11.A12 |
| methylbenzethonium | 35.03894628 | 35.05833814 | -0.019391857 | 5500024024213121906559.D | .H07.G08.E09.D10.B11.A12 |
| prilocaine | 33.43888513 | 33.45935203 | -0.020466896 | 5500024024213121906560.D | .H01.G02.E03.D04.B05.A06 |
| picotamide | 34.72486923 | 34.74789333 | -0.023024102 | 614615111406.D05 | .H01.G02.E03.D04.B05.A06 |
| tubocurarine chloride | 35.08740261 | 35.11129025 | -0.023887639 | 640641112706.C03 | .H01.G02.E03.D04.B05 |
| trichostatin A | 34.99590333 | 35.01997127 | -0.02406794 | 5500024024213121906559.F | .H01.G02.E03.D04.B05.A06 |
| trimetazidine | 35.08647252 | 35.11129025 | -0.024817726 | 6.41E+13 | .H01.G02.E03.D04.B05 |
| lynestrenol | 35.4029237 | 35.4281341 | -0.025210397 | 612613111306.B07 | .H07.G08.E09.D10.B11.A12 |
| carbarsone | 36.8146977 | 36.83996258 | -0.025264878 | 5500024024213121906563.G | .H07.G08.E09.D10.B11.A12 |
| flucloxacillin | 37.43382868 | 37.46016387 | -0.026335186 | 5500024030700072107987.D | .H07.G08.E09.D10.B11.A12 |
| rilmenidine | 37.43198271 | 37.46016387 | -0.028181156 | 5500024030700072107987.C | .H07.G08.E09.D10.B11.A12 |
| latamoxef | 34.18058615 | 34.21031976 | -0.029733609 | 5500024030760072207028.H | .H07.G08.E09.D10.B11.A12 |
| triamcinolone | 34.71772595 | 34.74789333 | -0.03016738 | 614615111406.B01 | .H01.G02.E03.D04.B05.A06 |
| eldeline | 35.0273913 | 35.05833814 | -0.030946839 | 5500024024213121906559.H | .H07.G08.E09.D10.B11.A12 |
| hydroxyzine | 35.39649782 | 35.4281341 | -0.031636273 | 6.13E+22 | .H07.G08.E09.D10.B11.A12 |
| isoxicam | 37.60632137 | 37.63864853 | -0.032327158 | 5500024024213121906564.C | .H07.G08.D10.B11.A12 |
| mephenesin | 33.42670707 | 33.45935203 | -0.032644958 | 5500024024213121906560.F | .H01.G02.E03.D04.B05.A06 |
| harpagoside | 34.67646533 | 34.70962788 | -0.033162552 | 5500024030760072207033.B | .H01.G02.E03.D04.B05.A06 |
| terguride | 37.21089761 | 37.24471367 | -0.033816065 | 5.50E+23 | .H01.G02.E03.D04.B05.A06 |
| phenylpropanolamine | 33.4237992 | 33.45797002 | -0.034170818 | 6.23E+18 | .H07.G08.E09.D10.B11.A12 |
| perphenazine | 35.39318626 | 35.4281341 | -0.034947836 | 612613111306.B10 | .H07.G08.E09.D10.B11.A12 |
| bemegride | 34.17535757 | 34.21031976 | -0.034962183 | 5500024030760072207028.C | .H07.G08.E09.D10.B11.A12 |
| colistin | 35.09645063 | 35.13162255 | -0.035171921 | 5500024024214122006604.C | .H07.G08.E09.D10.B11.A12 |
| tetracaine | 35.07438957 | 35.11129025 | -0.036900684 | 640641112706.C04 | .H01.G02.E03.D04.B05 |
| SR-95639A | 36.80216213 | 36.83996258 | -0.037800448 | 5500024024213121906563.C | .H07.G08.E09.D10.B11.A12 |
| Prestwick-1103 | 34.1373623 | 34.17633596 | -0.038973656 | 5500024030760072207033.A | .H07.G08.E09.B11.A12 |
| enilconazole | 37.42092909 | 37.46016387 | -0.039234783 | 5500024030700072107987.B | .H07.G08.E09.D10.B11.A12 |
| melatonin | 35.11596524 | 35.15552717 | -0.03956193 | 5500024024214122006604.F | .H01.G02.E03.D04.A06 |
| trichostatin A | 37.20663367 | 37.24787327 | -0.041239607 | 5500024035736031208612.D | .H01.G02.E03.D04.B05.A06 |
| cinnarizine | 33.41174337 | 33.4530122 | -0.041268829 | 622623112706.F03 | .H01.G02.E03.B05.A06 |
| flumetasone | 33.91983491 | 33.96129529 | -0.041460383 | 5500024025833041107269.G | .H07.E09.D10.B11.A12 |
| dicycloverine | 35.22532569 | 35.2669347 | -0.041609014 | 6.13E+17 | .H01.G02.E03.D04.B05.A06 |
| tobramycin | 35.08951895 | 35.13162255 | -0.042103601 | 5.50E+32 | .H07.G08.E09.D10.B11.A12 |
| pilocarpine | 35.11319478 | 35.15552717 | -0.04233239 | 5.50E+23 | .H01.G02.E03.D04.A06 |
| niflumic acid | 32.95695377 | 33.00109826 | -0.044144494 | 5500024024213121906560.C | .H07.G08.E09.D10.B11.A12 |
| desoxycortone | 37.20001594 | 37.24471367 | -0.044697734 | 5500024030700072107987.B | .H01.G02.E03.D04.B05.A06 |
| clozapine | 36.09730531 | 36.14209273 | -0.044787417 | 5500024017802120306174.C | .H01.G02.E03.D04.B05.A06 |
| lumicolchicine | 36.79510642 | 36.83996258 | -0.044856158 | 5500024024213121906563.F | .H07.G08.E09.D10.B11.A12 |
| meclozine | 35.1105889 | 35.15552717 | -0.044938268 | 5500024024214122006604.H | .H01.G02.E03.D04.A06 |
| debrisoquine | 33.41222845 | 33.45797002 | -0.045741567 | 622623112706.G07 | .H07.G08.E09.D10.B11.A12 |
| pempidine | 35.01173671 | 35.05833814 | -0.04660143 | 5500024024213121906559.H | .H07.G08.E09.D10.B11.A12 |
| hydrastinine | 35.0611964 | 35.10850208 | -0.047305676 | 614615111406.B07 | .H07.G08.E09.D10.B11.A12 |
| sulfinpyrazone | 33.40474931 | 33.4530122 | -0.048262892 | 622623112706.C05 | .H01.G02.E03.B05.A06 |
| dizocilpine | 34.69936464 | 34.74789333 | -0.048528685 | 614615111406.D03 | .H01.G02.E03.D04.B05.A06 |
| pyrazinamide | 35.08219943 | 35.13162255 | -0.04942312 | 5.50E+28 | .H07.G08.E09.D10.B11.A12 |
| cimetidine | 35.21726909 | 35.2669347 | -0.049665612 | 612613111306.H03 | .H01.G02.E03.D04.B05.A06 |
| cyclizine | 35.06142202 | 35.11129025 | -0.049868235 | 640641112706.D01 | .H01.G02.E03.D04.B05 |
| loracarbef | 34.12502085 | 34.17633596 | -0.051315111 | 5500024030760072207033.C | .H07.G08.E09.B11.A12 |
| saquinavir | 37.19583916 | 37.24787327 | -0.052034119 | 5500024035736031208612.G | .H01.G02.E03.D04.B05.A06 |
| sulfathiazole | 35.21481057 | 35.2669347 | -0.052124135 | 612613111306.H02 | .H01.G02.E03.D04.B05.A06 |
| protoveratrine A | 34.9667924 | 35.01997127 | -0.053178869 | 5.50E+23 | .H01.G02.E03.D04.B05.A06 |
| ioxaglic acid | 34.1214087 | 34.17633596 | -0.054927256 | 5500024030760072207033.C | .H07.G08.E09.B11.A12 |
| convolamine | 35.03126923 | 35.08636713 | -0.0550979 | 640641112706.C10 | .H07.G08.E09.D10.B11.A12 |
| ethionamide | 35.07537299 | 35.13162255 | -0.05624956 | 5500024024214122006604.B | .H07.G08.E09.D10.B11.A12 |
| lobelanidine | 35.05501755 | 35.11129025 | -0.056272699 | 640641112706.A02 | .H01.G02.E03.D04.B05 |
| colecalciferol | 35.09900986 | 35.15552717 | -0.056517305 | 5500024024214122006604.F | .H01.G02.E03.D04.A06 |
| 6-azathymine | 35.07489135 | 35.13162255 | -0.056731201 | 5500024024214122006604.H | .H07.G08.E09.D10.B11.A12 |
| diperodon | 33.39589964 | 33.4530122 | -0.057112565 | 622623112706.C06 | .H01.G02.E03.B05.A06 |
| bicuculline | 34.96244646 | 35.01997127 | -0.057524809 | 5500024024213121906559.F | .H01.G02.E03.D04.B05.A06 |
| tyloxapol | 37.18340806 | 37.24471367 | -0.061305615 | 5500024030700072107987.G | .H01.G02.E03.D04.B05.A06 |
| fenoprofen | 33.89992575 | 33.96129529 | -0.061369538 | 5500024025833041107269.G | .H07.E09.D10.B11.A12 |
| tracazolate | 34.6461178 | 34.70962788 | -0.063510078 | 5.50E+25 | .H01.G02.E03.D04.B05.A06 |
| ketotifen | 33.38912828 | 33.4530122 | -0.063883919 | 622623112706.A03 | .H01.G02.E03.B05.A06 |

| | | | | | |
|---|---|---|---|---|---|
| tetroquinone | 34.06925493 | 34.13330462 | -0.064049685 | 5.50E+22 | .H01.G02.D04.B05.A06 |
| chlorpromazine | 36.07800368 | 36.14209273 | -0.064089047 | 5500024017802120306174.F | .H01.G02.E03.D04.B05.A06 |
| ascorbic acid | 33.39251764 | 33.45797002 | -0.065452379 | 622623112706.D11 | .H07.G08.E09.D10.B11.A12 |
| testosterone | 36.94219041 | 37.00786033 | -0.065669915 | 5500024024213121906563.B | .H01.G02.E03.D04.B05.A06 |
| sulfasalazine | 35.04555672 | 35.11129025 | -0.065733528 | 640641112706.D03 | .H01.G02.E03.D04.B05 |
| butacaine | 34.33372104 | 34.40088765 | -0.067166606 | 5500024030401071707292.D | .H07.G08.E09.D10.B11 |
| lincomycin | 32.93284678 | 33.00109826 | -0.068251483 | 5500024024213121906560.G | .H07.G08.E09.D10.B11.A12 |
| indometacin | 32.93260888 | 33.00109826 | -0.068489382 | 5500024024213121906560.G | .H07.G08.E09.D10.B11.A12 |
| meprylcaine | 37.17930633 | 37.24787327 | -0.068566942 | 5500024035736031208612.H | .H01.G02.E03.D04.B05.A06 |
| kaempferol | 37.17892291 | 37.24787327 | -0.068950365 | 5500024035736031208612.B | .H01.G02.E03.D04.B05.A06 |
| xylazine | 34.95039088 | 35.01997127 | -0.069580391 | 5500024024213121906559.H | .H01.G02.E03.D04.B05.A06 |
| dantrolene | 33.38795315 | 33.45935203 | -0.071398883 | 5500024024213121906560.A | .H01.G02.E03.D04.B05.A06 |
| trichostatin A | 35.05917398 | 35.13162255 | -0.072448574 | 5500024024214122006604.F | .H07.G08.E09.D10.B11.A12 |
| sertaconazole | 37.17256885 | 37.24787327 | -0.075304427 | 5500024035736031208612.G | .H01.G02.E03.D04.B05.A06 |
| anabasine | 33.88201118 | 33.96076648 | -0.078755297 | 5500024025833041107269.G | .H01.G02.E03.D04.B05.A06 |
| (+/-)-catechin | 34.05073502 | 34.13330462 | -0.082569598 | 5500024030760072207028.C | .H01.G02.D04.B05.A06 |
| clidinium bromide | 34.31419324 | 34.40088765 | -0.086694412 | 5500024030401071707292.B | .H07.G08.E09.D10.B11 |
| pargyline | 35.02178553 | 35.10850208 | -0.086716544 | 614615111406.F11 | .H07.G08.E09.D10.B11.A12 |
| tomatidine | 34.99794551 | 35.08636713 | -0.088421614 | 640641112706.H11 | .H07.G08.E09.D10.B11.A12 |
| Prestwick-665 | 34.9692797 | 35.05833814 | -0.089058438 | 5.50E+31 | .H07.G08.E09.D10.B11.A12 |
| rosiglitazone | 34.33068292 | 34.42118584 | -0.090502915 | 5500024030401071707292.C | .H01.G02.E03.D04.B05.A06 |
| alpha-estradiol | 37.22206246 | 37.31290545 | -0.090842993 | 5500024035736031208612.G | .G08.E09.D10.B11.A12 |
| racecadotril | 34.99530435 | 35.08636713 | -0.091062782 | 640641112706.B07 | .H07.G08.E09.D10.B11.A12 |
| bethanechol | 34.08499864 | 34.17633596 | -0.091337317 | 5500024030760072207033.B | .H07.G08.E09.B11.A12 |
| spaglumic acid | 34.08358887 | 34.17633596 | -0.092747082 | 5500024030760072207033.D | .H07.G08.E09.B11.A12 |
| scopolamine N-oxide | 35.01524555 | 35.10850208 | -0.09325653 | 614615111406.F08 | .H07.G08.E09.D10.B11.A12 |
| trichostatin A | 34.0830534 | 34.17633596 | -0.093282555 | 5500024030760072207033.G | .H07.G08.E09.B11.A12 |
| propylthiouracil | 35.03816644 | 35.13162255 | -0.093456114 | 5500024024214122006604.F | .H07.G08.E09.D10.B11.A12 |
| midecamycin | 35.33394023 | 35.4281341 | -0.094193871 | 612613111306.D07 | .H07.G08.E09.D10.B11.A12 |
| bephenium hydroxyna | 37.1492529 | 37.24471367 | -0.095460766 | 5500024030700072107987.D | .H01.G02.E03.D04.B05.A06 |
| mesoridazine | 35.01479435 | 35.11129025 | -0.096495901 | 640641112706.F06 | .H01.G02.E03.D04.B05 |
| L-methionine sulfoxim | 35.03489721 | 35.13162255 | -0.096725339 | 5500024024214122006604.G | .H07.G08.E09.D10.B11.A12 |
| nipecotic acid | 37.35989886 | 37.46016387 | -0.100265011 | 5.50E+28 | .H07.G08.E09.D10.B11.A12 |
| chlorogenic acid | 36.73809708 | 36.83996258 | -0.101865497 | 5500024024213121906563.A | .H07.G08.E09.D10.B11.A12 |
| trichostatin A | 34.31544814 | 34.42118584 | -0.105737695 | 5500024030401071707292.G | .H01.G02.E03.D04.B05.A06 |
| promazine | 34.95211497 | 35.05833814 | -0.106223167 | 5500024024213121906559.G | .H07.G08.E09.D10.B11.A12 |
| antazoline | 33.34639035 | 33.4530122 | -0.106621852 | 622623112706.F01 | .H01.G02.E03.B05.A06 |
| amrinone | 34.29403864 | 34.40088765 | -0.106849012 | 5500024030401071707292.D | .H07.G08.E09.D10.B11 |
| paromomycin | 34.02586829 | 34.13330462 | -0.107436326 | 5500024030760072207028.B | .H01.G02.D04.B05.A06 |
| hydrochlorothiazide | 35.15907038 | 35.2669347 | -0.107864327 | 612613111306.D05 | .H01.G02.E03.D04.B05.A06 |
| phenazone | 35.15858827 | 35.2669347 | -0.108346433 | 612613111306.C01 | .H01.G02.E03.D04.B05.A06 |
| benzydamine | 33.34217576 | 33.4530122 | -0.110836441 | 622623112706.G03 | .H01.G02.E03.B05.A06 |
| dropropizine | 32.88978885 | 33.00109826 | -0.111309412 | 5500024024213121906560.C | .H07.G08.E09.D10.B11.A12 |
| pyrithyldione | 34.28926404 | 34.40088765 | -0.111623613 | 5500024030401071707292.A | .H07.G08.E09.D10.B11 |
| sulfabenzamide | 34.90802453 | 35.01997127 | -0.111946737 | 5500024024213121906559.B | .H01.G02.E03.D04.B05.A06 |
| thioproperazine | 34.63557815 | 34.74789333 | -0.112315178 | 614615111406.C02 | .H01.G02.E03.D04.B05.A06 |
| aminophylline | 34.09643693 | 34.21031976 | -0.113882823 | 5500024030760072207028.F | .H07.G08.E09.D10.B11.A12 |
| Prestwick-967 | 34.59485537 | 34.70962788 | -0.114772509 | 5500024030760072207033.D | .H01.G02.E03.D04.B05.A06 |
| valproic acid | 34.30427396 | 34.42118584 | -0.116911879 | 5500024030401071707292.F | .H01.G02.E03.D04.B05.A06 |
| mepenzolate bromide | 34.94060932 | 35.05833814 | -0.117728819 | 5500024024213121906559.H | .H07.G08.E09.D10.B11.A12 |
| pancuronium bromide | 34.59163714 | 34.70962788 | -0.11799074 | 5500024030760072207033.G | .H01.G02.E03.D04.B05.A06 |
| conessine | 34.90181244 | 35.01997127 | -0.118158833 | 5500024024213121906559.G | .H01.G02.E03.D04.B05.A06 |
| 3-acetamidocoumarir | 34.58757838 | 34.70962788 | -0.122049504 | 5500024030760072207033.A | .H01.G02.E03.D04.B05.A06 |
| diethylcarbamazine | 32.87882631 | 33.00109826 | -0.122271954 | 5500024024213121906560.D | .H07.G08.E09.D10.B11.A12 |
| simvastatin | 34.01086392 | 34.13330462 | -0.122440701 | 5.50E+26 | .H01.G02.D04.B05.A06 |
| brompheniramine | 36.71667842 | 36.83996258 | -0.123284159 | 5500024024213121906563.C | .H07.G08.E09.D10.B11.A12 |
| moxisylyte | 37.51457586 | 37.63864853 | -0.124072669 | 5500024024213121906564.F | .H07.G08.D10.B11.A12 |
| phthalylsulfathiazole | 34.08534211 | 34.21031976 | -0.124977651 | 5500024030760072207028.G | .H07.G08.E09.D10.B11.A12 |
| delsoline | 33.83547083 | 33.96076648 | -0.125295646 | 5500024025833041107269.F | .H01.G02.E03.D04.B05.A06 |
| sulfamonomethoxine | 34.27436122 | 34.40088765 | -0.126526431 | 5500024030401071707292.A | .H07.G08.E09.D10.B11 |
| trimethadione | 35.00410645 | 35.13162255 | -0.127516108 | 5500024024214122006604.D | .H07.G08.E09.D10.B11.A12 |
| amodiaquine | 33.32507519 | 33.4530122 | -0.127937016 | 622623112706.D05 | .H01.G02.E03.B05.A06 |
| bepridil | 36.87976231 | 37.00786033 | -0.128098021 | 5500024024213121906563.F | .H01.G02.E03.D04.B05.A06 |
| betulin | 34.0481478 | 34.17633596 | -0.128188161 | 5500024030760072207033.F | .H07.G08.E09.B11.A12 |
| Prestwick-857 | 34.00451536 | 34.13330462 | -0.128789253 | 5500024030760072207028.B | .H01.G02.D04.B05.A06 |
| valproic acid | 34.29159356 | 34.42118584 | -0.129592273 | 5500024030401071707292.A | .H01.G02.E03.D04.B05.A06 |
| talampicillin | 34.0454833 | 34.17633596 | -0.130852651 | 5500024030760072207033.F | .H07.G08.E09.B11.A12 |
| aciclovir | 35.29579398 | 35.4281341 | -0.132340115 | 612613111306.A09 | .H07.G08.E09.D10.B11.A12 |
| roxithromycin | 34.00053214 | 34.13330462 | -0.13277248 | 5500024030760072207028.G | .H01.G02.D04.B05.A06 |

| | | | | | |
|---|---|---|---|---|---|
| pralidoxime | 37.11112982 | 37.24471367 | -0.13358385 | 5500024030700072107987.H | .H01.G02.E03.D04.B05.A06 |
| troglitazone | 37.17754442 | 37.31290545 | -0.135361033 | 5500024035736031208612.C | .G08.E09.D10.B11.A12 |
| SC-58125 | 35.02595355 | 35.1615202 | -0.135566651 | EC2004073016AA | EC2004073014AA |
| sulfamethizole | 33.82225991 | 33.96076648 | -0.13850657 | 5500024025833041107269.B | .H01.G02.E03.D04.B05.A06 |
| dimenhydrinate | 32.86170981 | 33.00109826 | -0.139388454 | 5500024024213121906560.C | .H07.G08.E09.D10.B11.A12 |
| acacetin | 34.91883414 | 35.05833814 | -0.139504 | 5500024024213121906559.D | .H07.G08.E09.D10.B11.A12 |
| pheniramine | 35.12713499 | 35.2669347 | -0.139799719 | 612613111306.C04 | .H01.G02.E03.D04.B05.A06 |
| sulmazole | 34.88000673 | 35.01997127 | -0.139964539 | 5500024024213121906559.C | .H01.G02.E03.D04.B05.A06 |
| tolmetin | 33.99297122 | 34.13330462 | -0.140333399 | 5500024030760072207028.C | .H01.G02.D04.B05.A06 |
| etifenin | 34.99099508 | 35.13162255 | -0.140627472 | 5500024024214122006604.F | .H07.G08.E09.D10.B11.A12 |
| metacycline | 34.56748089 | 34.70962788 | -0.142146989 | 5500024030760072207033.H | .H01.G02.E03.D04.B05.A06 |
| ciprofloxacin | 35.2858149 | 35.4281341 | -0.142319199 | 6.13E+19 | .H07.G08.E09.D10.B11.A12 |
| propofol | 34.06510768 | 34.21031976 | -0.145212072 | 5500024030760072207028.D | .H07.G08.E09.D10.B11.A12 |
| selegiline | 34.98361077 | 35.13162255 | -0.148011788 | 5500024024214122006604.H | .H07.G08.E09.D10.B11.A12 |
| palmatine | 34.87189502 | 35.01997127 | -0.148076253 | 5500024024213121906559.F | .H01.G02.E03.D04.B05.A06 |
| fipexide | 33.30459395 | 33.4530122 | -0.148418256 | 622623112706.F05 | .H01.G02.E03.B05.A06 |
| trioxysalen | 33.81130935 | 33.96076648 | -0.149457126 | 5500024025833041107269.F | .H01.G02.E03.D04.B05.A06 |
| estradiol | 35.9921818 | 36.14209273 | -0.149910925 | 5500024017802120306174.A | .H01.G02.E03.D04.B05.A06 |
| droperidol | 36.85728516 | 37.00786033 | -0.150575167 | 5500024024213121906563.C | .H01.G02.E03.D04.B05.A06 |
| molindone | 34.55820532 | 34.70962788 | -0.151422557 | 5.50E+22 | .H01.G02.E03.D04.B05.A06 |
| ampicillin | 35.27599959 | 35.4281341 | -0.15213451 | 612613111306.D12 | .H07.G08.E09.D10.B11.A12 |
| chlorcyclizine | 34.9061437 | 35.05833814 | -0.152194444 | 5500024024213121906559.C | .H07.G08.E09.D10.B11.A12 |
| diclofenamide | 34.0567124 | 34.21031976 | -0.153607357 | 5500024030760072207028.H | .H07.G08.E09.D10.B11.A12 |
| mefloquine | 34.59395921 | 34.74789333 | -0.153934117 | 614615111406.H03 | .H01.G02.E03.D04.B05.A06 |
| famotidine | 35.27026367 | 35.4281341 | -0.157870429 | 612613111306.D11 | .H07.G08.E09.D10.B11.A12 |
| hydrastine hydrochlor | 34.95188004 | 35.11129025 | -0.159410207 | 640641112706.C05 | .H01.G02.E03.D04.B05 |
| lomefloxacin | 33.29557725 | 33.45935203 | -0.163774778 | 5.50E+25 | .H01.G02.E03.D04.B05.A06 |
| oxetacaine | 35.10311129 | 35.2669347 | -0.163823417 | 612613111306.D01 | .H01.G02.E03.D04.B05.A06 |
| idazoxan | 37.08052691 | 37.24471367 | -0.164186759 | 5500024030700072107987.D | .H01.G02.E03.D04.B05.A06 |
| dihydrostreptomycin | 34.58161734 | 34.74789333 | -0.166275993 | 614615111406.C03 | .H01.G02.E03.D04.B05.A06 |
| iobenguane | 34.94493689 | 35.11129025 | -0.16635336 | 6.41E+16 | .H01.G02.E03.D04.B05 |
| carbamazepine | 37.47138203 | 37.63864853 | -0.167266499 | 5500024024213121906564.F | .H07.G08.D10.B11.A12 |
| ornidazole | 34.94045329 | 35.10850208 | -0.168048782 | 614615111406.D07 | .H07.G08.E09.D10.B11.A12 |
| Prestwick-1085 | 37.07882914 | 37.24787327 | -0.169044137 | 5500024035736031208612.F | .H01.G02.E03.D04.B05.A06 |
| mefexamide | 34.93684519 | 35.10850208 | -0.171656883 | 614615111406.B09 | .H07.G08.E09.D10.B11.A12 |
| physostigmine | 34.91437446 | 35.08636713 | -0.171992668 | 640641112706.C07 | .H07.G08.E09.D10.B11.A12 |
| pizotifen | 37.28740369 | 37.46016387 | -0.172760181 | 5500024030700072107987.C | .H07.G08.E09.D10.B11.A12 |
| trichostatin A | 37.13814641 | 37.31290545 | -0.174759045 | 5500024035736031208612.G | .G08.E09.D10.B11.A12 |
| flucytosine | 37.06959475 | 37.24471367 | -0.175118916 | 5500024030700072107987.G | .H01.G02.E03.D04.B05.A06 |
| Prestwick-864 | 33.95782095 | 34.13330462 | -0.175483662 | 5500024030760072207028.F | .H01.G02.D04.B05.A06 |
| tiratricol | 34.93062786 | 35.10850208 | -0.177874211 | 614615111406.G11 | .H07.G08.E09.D10.B11.A12 |
| pirenzepine | 34.56779829 | 34.74789333 | -0.180095036 | 614615111406.D06 | .H01.G02.E03.D04.B05.A06 |
| LY-294002 | 34.23999279 | 34.42118584 | -0.181193047 | 5500024030401071707292.B | .H01.G02.E03.D04.B05.A06 |
| butirosin | 33.77693419 | 33.96076648 | -0.18383229 | 5500024025833041107269.F | .H01.G02.E03.D04.B05.A06 |
| methazolamide | 34.21676288 | 34.40088765 | -0.184124772 | 5500024030401071707292.C | .H07.G08.E09.D10.B11 |
| pentamidine | 34.94481375 | 35.13162255 | -0.186808806 | 5500024024214122006604.F | .H07.G08.E09.D10.B11.A12 |
| articaine | 37.27280565 | 37.46016387 | -0.187358215 | 5500024030700072107987.B | .H07.G08.E09.D10.B11.A12 |
| alvespimycin | 37.12059947 | 37.31290545 | -0.192305978 | 5500024035736031208612.F | .G08.E09.D10.B11.A12 |
| oxantel | 36.81449914 | 37.00786033 | -0.193361191 | 5500024024213121906563.F | .H01.G02.E03.D04.B05.A06 |
| ticarcillin | 37.05436916 | 37.24787327 | -0.193504119 | 5500024035736031208612.D | .H01.G02.E03.D04.B05.A06 |
| pyrimethamine | 35.07070664 | 35.2669347 | -0.196228065 | 612613111306.F02 | .H01.G02.E03.D04.B05.A06 |
| chlorpromazine | 37.11652481 | 37.31290545 | -0.196380639 | 5500024035736031208612.F | .G08.E09.D10.B11.A12 |
| tacrine | 36.81067444 | 37.00786033 | -0.197185895 | 5500024024213121906563.F | .H01.G02.E03.D04.B05.A06 |
| levopropoxyphene | 33.978593 | 34.17633596 | -0.197742956 | 5500024030760072207033.A | .H07.G08.E09.B11.A12 |
| 1,5-isoquinolinediol | 34.96314706 | 35.1615202 | -0.198373145 | EC2004073017AA | EC2004073014AA |
| N-acetyl-L-aspartic ac | 36.64112699 | 36.83996258 | -0.198835584 | 5500024024213121906563.D | .H07.G08.E09.D10.B11.A12 |
| etamsylate | 34.51071178 | 34.70962788 | -0.198916104 | 5500024030760072207033.F | .H01.G02.E03.D04.B05.A06 |
| indapamide | 33.26021853 | 33.45935203 | -0.199133498 | 5500024024213121906560.B | .H01.G02.E03.D04.B05.A06 |
| fulvestrant | 37.11259634 | 37.31290545 | -0.200309113 | 5500024035736031208612.H | .G08.E09.D10.B11.A12 |
| zomepirac | 34.19851769 | 34.40088765 | -0.202369955 | 5500024030401071707292.F | .H07.G08.E09.D10.B11 |
| levodopa | 35.06256 | 35.2669347 | -0.204374703 | 612613111306.G06 | .H01.G02.E03.D04.B05.A06 |
| dehydrocholic acid | 35.22232082 | 35.4281341 | -0.205813282 | 6.13E+21 | .H07.G08.E09.D10.B11.A12 |
| atractyloside | 33.75471287 | 33.96129529 | -0.206582423 | 5500024025833041107269.B | .H07.E09.D10.B11.A12 |
| benzthiazide | 33.9260692 | 34.13330462 | -0.207235413 | 5500024030760072207028.G | .H01.G02.D04.B05.A06 |
| alimemazine | 34.19338782 | 34.40088765 | -0.207499824 | 5500024030401071707292.B | .H07.G08.E09.D10.B11 |
| promethazine | 37.03658839 | 37.24471367 | -0.208125283 | 5500024030700072107987.B | .H01.G02.E03.D04.B05.A06 |
| tropicamide | 33.2501084 | 33.45935203 | -0.209243629 | 5.50E+23 | .H01.G02.E03.D04.B05.A06 |
| dexpanthenol | 37.42851792 | 37.63864853 | -0.210130604 | 5500024024213121906564.F | .H07.G08.D10.B11.A12 |
| doxylamine | 35.05660704 | 35.2669347 | -0.210327668 | 612613111306.F01 | .H01.G02.E03.D04.B05.A06 |

| | | | | | |
|---|---|---|---|---|---|
| bacitracin | 37.24884459 | 37.46016387 | -0.211319279 | 5500024030700072107987.H | .H07.G08.E09.D10.B11.A12 |
| esculetin | 37.24882544 | 37.46016387 | -0.211338427 | 5500024030700072107987.F | .H07.G08.E09.D10.B11.A12 |
| sirolimus | 34.20969878 | 34.42118584 | -0.211487055 | 5500024030401071707292.A | .H01.G02.E03.D04.B05.A06 |
| terguride | 37.0314847 | 37.24471367 | -0.21322897 | 5500024030700072107987.C | .H01.G02.E03.D04.B05.A06 |
| betaxolol | 33.24326526 | 33.45797002 | -0.214704753 | 622623112706.G09 | .H07.G08.E09.D10.B11.A12 |
| guaifenesin | 33.74627096 | 33.96129529 | -0.215024333 | 5500024025833041107269.C | .H07.E09.D10.B11.A12 |
| khellin | 35.05157545 | 35.2669347 | -0.215359252 | 612613111306.A05 | .H01.G02.E03.D04.B05.A06 |
| vincamine | 33.24327295 | 33.45935203 | -0.216079075 | 5500024024213121906560.A | .H01.G02.E03.D04.B05.A06 |
| resveratrol | 34.89512015 | 35.11129025 | -0.216170097 | 640641112706.G01 | .H01.G02.E03.D04.B05 |
| meropenem | 37.03023977 | 37.24787327 | -0.217633509 | 5.50E+27 | .H01.G02.E03.D04.B05.A06 |
| fluphenazine | 34.20282426 | 34.42118584 | -0.218361572 | 5500024030401071707292.B | .H01.G02.E03.D04.B05.A06 |
| oxolamine | 33.91467484 | 34.13330462 | -0.218629778 | 5500024030760072207028.D | .H01.G02.D04.B05.A06 |
| bisacodyl | 34.93662371 | 35.15552717 | -0.218903458 | 5500024024214122006604.F | .H01.G02.E03.D04.A06 |
| prochlorperazine | 36.78787887 | 37.00786033 | -0.219981463 | 5500024024213121906563.D | .H01.G02.E03.D04.B05.A06 |
| digoxin | 34.9322808 | 35.15552717 | -0.223246366 | 5500024024214122006604.H | .H01.G02.E03.D04.A06 |
| etofenamate | 34.48570743 | 34.70962788 | -0.223920453 | 5500024030760072207033.G | .H01.G02.E03.D04.B05.A06 |
| theophylline | 33.90936263 | 34.13330462 | -0.223941985 | 5500024030760072207028.H | .H01.G02.D04.B05.A06 |
| Prestwick-675 | 34.83049714 | 35.05833814 | -0.227840996 | 5.50E+32 | .H07.G08.E09.D10.B11.A12 |
| alpha-estradiol | 34.19329469 | 34.42118584 | -0.227891147 | 5500024030401071707292.G | .H01.G02.E03.D04.B05.A06 |
| fenbufen | 33.23046029 | 33.45935203 | -0.228891741 | 5.50E+22 | .H01.G02.E03.D04.B05.A06 |
| piroxicam | 34.87911037 | 35.10850208 | -0.229391702 | 614615111406.H09 | .H07.G08.E09.D10.B11.A12 |
| progesterone | 34.92611883 | 35.15552717 | -0.229408338 | 5500024024214122006604.H | .H01.G02.E03.D04.A06 |
| hydroflumethiazide | 37.40568287 | 37.63864853 | -0.232965658 | 5.50E+31 | .H07.G08.D10.B11.A12 |
| chlormezanone | 33.22463911 | 33.45797002 | -0.23333091 | 622623112706.B09 | .H07.G08.E09.D10.B11.A12 |
| pentoxyverine | 36.77296958 | 37.00786033 | -0.234890751 | 5500024024213121906563.G | .H01.G02.E03.D04.B05.A06 |
| piromidic acid | 33.89833615 | 34.13330462 | -0.234968464 | 5500024030760072207028.F | .H01.G02.D04.B05.A06 |
| niridazole | 34.91975529 | 35.15552717 | -0.235771879 | 5.50E+26 | .H01.G02.E03.D04.A06 |
| quinisocaine | 34.78391123 | 35.01997127 | -0.236060042 | 5500024024213121906559.D | .H01.G02.E03.D04.B05.A06 |
| dexpropranolol | 37.00784484 | 37.24787327 | -0.240028437 | 5500024035736031208612.G | .H01.G02.E03.D04.B05.A06 |
| quercetin | 34.89044508 | 35.13162255 | -0.241177472 | 5500024024214122006604.A | .H07.G08.E09.D10.B11.A12 |
| cantharidin | 37.00250182 | 37.24471367 | -0.242211855 | 5500024030700072107987.F | .H01.G02.E03.D04.B05.A06 |
| furaltadone | 33.71809197 | 33.96129529 | -0.243203317 | 5500024025833041107269.F | .H07.E09.D10.B11.A12 |
| iopromide | 34.15753803 | 34.40088765 | -0.243349618 | 5500024030401071707292.A | .H07.G08.E09.D10.B11 |
| miconazole | 35.02340337 | 35.2669347 | -0.243531334 | 612613111306.F05 | .H01.G02.E03.D04.B05.A06 |
| sulfachlorpyridazine | 34.81262612 | 35.05833814 | -0.245712021 | 5500024024213121906559.D | .H07.G08.E09.D10.B11.A12 |
| trichostatin A | 34.83798273 | 35.08636713 | -0.248384394 | 640641112706.B10 | .H07.G08.E09.D10.B11.A12 |
| sulfaguanidine | 35.01744272 | 35.2669347 | -0.249491986 | 612613111306.B01 | .H01.G02.E03.D04.B05.A06 |
| harman | 34.77035833 | 35.01997127 | -0.249612936 | 5500024024213121906559.D | .H01.G02.E03.D04.B05.A06 |
| nitrofurantoin | 33.20886067 | 33.45935203 | -0.250491358 | 5500024024213121906560.F | .H01.G02.E03.D04.B05.A06 |
| aztreonam | 34.85781011 | 35.10850208 | -0.250691965 | 614615111406.C12 | .H07.G08.E09.D10.B11.A12 |
| gibberellic acid | 34.45713904 | 34.70962788 | -0.25248884 | 5500024030760072207033.G | .H01.G02.E03.D04.B05.A06 |
| zoxazolamine | 36.75509387 | 37.00786033 | -0.252766456 | 5500024024213121906563.G | .H01.G02.E03.D04.B05.A06 |
| azlocillin | 34.14428487 | 34.40088765 | -0.25660278 | 5500024030401071707292.D | .H07.G08.E09.D10.B11 |
| moxonidine | 34.45287065 | 34.70962788 | -0.256757231 | 5500024030760072207033.D | .H01.G02.E03.D04.B05.A06 |
| meticrane | 37.37884146 | 37.63864853 | -0.259807065 | 5500024024213121906564.H | .H07.G08.D10.B11.A12 |
| etidronic acid | 33.8724712 | 34.13330462 | -0.260833419 | 5500024030760072207028.H | .H01.G02.D04.B05.A06 |
| guanfacine | 36.7462256 | 37.00786033 | -0.261634733 | 5.50E+22 | .H01.G02.E03.D04.B05.A06 |
| trichostatin A | 35.0009978 | 35.2669347 | -0.265936908 | 612613111306.G05 | .H01.G02.E03.D04.B05.A06 |
| orphenadrine | 33.19308114 | 33.45935203 | -0.266270892 | 5500024024213121906560.C | .H01.G02.E03.D04.B05.A06 |
| nifurtimox | 34.44224034 | 34.70962788 | -0.267387544 | 5500024030760072207033.G | .H01.G02.E03.D04.B05.A06 |
| sparteine | 34.75116839 | 35.01997127 | -0.268802883 | 5500024024213121906559.G | .H01.G02.E03.D04.B05.A06 |
| amiprilose | 33.86250501 | 34.13330462 | -0.27079961 | 5.50E+23 | .H01.G02.D04.B05.A06 |
| estradiol | 36.7363699 | 37.00786033 | -0.271490428 | 5500024024213121906563.B | .H01.G02.E03.D04.B05.A06 |
| glycopyrronium bromi | 33.68783247 | 33.96129529 | -0.27346282 | 5500024025833041107269.D | .H07.E09.D10.B11.A12 |
| eucatropine | 33.6860598 | 33.96129529 | -0.275235486 | 5500024025833041107269.F | .H07.E09.D10.B11.A12 |
| bendroflumethiazide | 33.68380822 | 33.96129529 | -0.277487065 | 5500024025833041107269.F | .H07.E09.D10.B11.A12 |
| liothyronine | 33.85524931 | 34.13330462 | -0.278055304 | 5500024030760072207028.H | .H01.G02.D04.B05.A06 |
| natamycin | 36.96969994 | 37.24787327 | -0.278173332 | 5500024035736031208612.H | .H01.G02.E03.D04.B05.A06 |
| aminocaproic acid | 37.18195332 | 37.46016387 | -0.27821055 | 5.50E+29 | .H07.G08.E09.D10.B11.A12 |
| nisoxetine | 37.18140438 | 37.46016387 | -0.278759484 | 5500024030700072107987.F | .H07.G08.E09.D10.B11.A12 |
| dioxybenzone | 36.9654051 | 37.24471367 | -0.279308575 | 5500024030700072107987.B | .H01.G02.E03.D04.B05.A06 |
| roxarsone | 33.89411256 | 34.17633596 | -0.282223396 | 5500024030760072207033.G | .H07.G08.E09.B11.A12 |
| carisoprodol | 36.55405103 | 36.83996258 | -0.285911544 | 5500024024213121906563.G | .H07.G08.E09.D10.B11.A12 |
| aminohippuric acid | 36.95729557 | 37.24471367 | -0.287418098 | 5500024030700072107987.F | .H01.G02.E03.D04.B05.A06 |
| lisinopril | 32.71242183 | 33.00109826 | -0.288676436 | 5500024024213121906560.H | .H07.G08.E09.D10.B11.A12 |
| metformin | 37.3473092 | 37.63864853 | -0.291339327 | 5500024024213121906564.D | .H07.G08.D10.B11.A12 |
| drofenine | 34.10829735 | 34.40088765 | -0.2925903 | 5500024030401071707292.F | .H07.G08.E09.D10.B11 |
| ethisterone | 33.16535665 | 33.45935203 | -0.293995377 | 5500024024213121906560.A | .H01.G02.E03.D04.B05.A06 |
| oxybutynin | 33.15679555 | 33.4530122 | -0.296216648 | 622623112706.G01 | .H01.G02.E03.B05.A06 |

| | | | | | |
|---|---|---|---|---|---|
| furazolidone | 33.83597628 | 34.13330462 | -0.297328339 | 5500024030760072207028.A( | .H01.G02.D04.B05.A06 |
| deptropine | 37.16104221 | 37.46016387 | -0.299121661 | 5500024030700072107987.A( | .H07.G08.E09.D10.B11.A12 |
| trichostatin A | 37.15982442 | 37.46016387 | -0.300339444 | 5500024030700072107987.G | .H07.G08.E09.D10.B11.A12 |
| loperamide | 35.12205899 | 35.4281341 | -0.306075108 | 612613111306.C09 | .H07.G08.E09.D10.B11.A12 |
| Prestwick-984 | 34.40089176 | 34.70962788 | -0.308736123 | 5500024030760072207033.H | .H01.G02.E03.D04.B05.A06 |
| propranolol | 33.90035856 | 34.21031976 | -0.309961201 | 5500024030760072207028.B | .H07.G08.E09.D10.B11.A12 |
| denatonium benzoate | 37.1480139 | 37.46016387 | -0.312149964 | 5.50E+31 | .H07.G08.E09.D10.B11.A12 |
| solasodine | 34.74483395 | 35.05833814 | -0.313504189 | 5500024024213121906559.H | .H07.G08.E09.D10.B11.A12 |
| phentolamine | 33.14338365 | 33.45935203 | -0.315968376 | 5500024024213121906560.B( | .H01.G02.E03.D04.B05.A06 |
| buflomedil | 36.52345469 | 36.83996258 | -0.31650789 | 5500024024213121906563.B( | .H07.G08.E09.D10.B11.A12 |
| clomifene | 36.69070222 | 37.00786033 | -0.31715811 | 5500024024213121906563.G | .H01.G02.E03.D04.B05.A06 |
| heliotrine | 34.73632539 | 35.05833814 | -0.322012751 | 5500024024213121906559.F( | .H07.G08.E09.D10.B11.A12 |
| felbinac | 33.88561977 | 34.21031976 | -0.324699986 | 5500024030760072207028.A( | .H07.G08.E09.D10.B11.A12 |
| ciclosporin | 36.51302164 | 36.83996258 | -0.326940943 | 5500024024213121906563.C | .H07.G08.E09.D10.B11.A12 |
| lymecycline | 33.84880532 | 34.17633596 | -0.327530639 | 5500024030760072207033.F( | .H07.G08.E09.B11.A12 |
| canavanine | 34.69207838 | 35.01997127 | -0.327892886 | 5500024024213121906559.F( | .H01.G02.E03.D04.B05.A06 |
| lidocaine | 34.93703463 | 35.2669347 | -0.329900073 | 612613111306.B06 | .H01.G02.E03.D04.B05.A06 |
| ouabain | 36.67557531 | 37.00786033 | -0.332285022 | 5500024024213121906563.A( | .H01.G02.E03.D04.B05.A06 |
| chloramphenicol | 37.30591632 | 37.63864853 | -0.332732213 | 5500024024213121906564.H | .H07.G08.D10.B11.A12 |
| trichostatin A | 36.67357514 | 37.00786033 | -0.334285189 | 5500024024213121906563.D | .H01.G02.E03.D04.B05.A06 |
| LY-294002 | 36.97836664 | 37.31290545 | -0.334538813 | 5500024035736031208612.F | .G08.E09.D10.B11.A12 |
| doxycycline | 34.0653892 | 34.40088765 | -0.335498445 | 5500024030401071707292.B | .H07.G08.E09.D10.B11 |
| proglumide | 33.12330898 | 33.45935203 | -0.336043046 | 5500024024213121906560.B( | .H01.G02.E03.D04.B05.A06 |
| benzamil | 34.72211854 | 35.05833814 | -0.336219603 | 5500024024213121906559.B( | .H07.G08.E09.D10.B11.A12 |
| tiabendazole | 34.793074 | 35.13162255 | -0.338548553 | 5.50E+29 | .H07.G08.E09.D10.B11.A12 |
| kawain | 33.11947988 | 33.45935203 | -0.339872153 | 5500024024213121906560.G | .H01.G02.E03.D04.B05.A06 |
| halofantrine | 37.11953012 | 37.46016387 | -0.340633745 | 5500024030700072107987.D | .H07.G08.E09.D10.B11.A12 |
| epirizole | 37.29636492 | 37.63864853 | -0.342283613 | 5500024024213121906564.F( | .H07.G08.D10.B11.A12 |
| Chicago Sky Blue 6B | 36.49611758 | 36.83996258 | -0.343844999 | 5500024024213121906563.D | .H07.G08.E09.D10.B11.A12 |
| imipenem | 34.7664686 | 35.11129025 | -0.344821649 | 640641112706.F05 | .H01.G02.E03.D04.B05 |
| amprolium | 34.92157098 | 35.2669347 | -0.345363724 | 6.13E+12 | .H01.G02.E03.D04.B05.A06 |
| acetylsalicylic acid | 36.96199131 | 37.31290545 | -0.350914141 | 5500024035736031208612.H | .G08.E09.D10.B11.A12 |
| cloxacillin | 34.75750575 | 35.10850208 | -0.350996329 | 614615111406.A09 | .H07.G08.E09.D10.B11.A12 |
| trichostatin A | 34.75440601 | 35.10850208 | -0.354096064 | 6.15E+19 | .H07.G08.E09.D10.B11.A12 |
| midodrine | 34.39368705 | 34.74789333 | -0.354206279 | 614615111406.A05 | .H01.G02.E03.D04.B05.A06 |
| diltiazem | 35.07338808 | 35.4281341 | -0.354746013 | 612613111306.C08 | .H07.G08.E09.D10.B11.A12 |
| LY-294002 | 36.95787404 | 37.31290545 | -0.355031407 | 5500024035736031208612.B( | .G08.E09.D10.B11.A12 |
| sitosterol | 34.35409907 | 34.70962788 | -0.355528814 | 5500024030760072207033.F( | .H01.G02.E03.D04.B05.A06 |
| trichostatin A | 34.75370482 | 35.11129025 | -0.357585432 | 640641112706.D02 | .H01.G02.E03.D04.B05 |
| cefotaxime | 34.38400123 | 34.74789333 | -0.363892103 | 614615111406.C01 | .H01.G02.E03.D04.B05.A06 |
| diloxanide | 33.84486409 | 34.21031976 | -0.365455667 | 5500024030760072207028.A( | .H07.G08.E09.D10.B11.A12 |
| meclofenamic acid | 34.74188851 | 35.10850208 | -0.366613569 | 614615111406.A11 | .H07.G08.E09.D10.B11.A12 |
| proscillaridin | 34.34077882 | 34.70962788 | -0.368849062 | 5.50E+26 | .H01.G02.E03.D04.B05.A06 |
| sulfaquinoxaline | 33.59180214 | 33.96076648 | -0.36896434 | 5500024025833041107269.D | .H01.G02.E03.D04.B05.A06 |
| josamycin | 35.05567715 | 35.4281341 | -0.372456952 | 612613111306.C10 | .H07.G08.E09.D10.B11.A12 |
| Prestwick-1084 | 36.87232086 | 37.24787327 | -0.375552414 | 5500024035736031208612.H | .H01.G02.E03.D04.B05.A06 |
| dicoumarol | 33.57948872 | 33.96129529 | -0.381806568 | 5.50E+33 | .H07.E09.D10.B11.A12 |
| sulfamethoxazole | 33.07656102 | 33.45935203 | -0.38279101 | 5500024024213121906560.H | .H01.G02.E03.D04.B05.A06 |
| novobiocin | 33.75035785 | 34.13330462 | -0.382946763 | 5500024030760072207028.G | .H01.G02.D04.B05.A06 |
| pentetic acid | 33.82655991 | 34.21031976 | -0.383759841 | 5500024030760072207028.D | .H07.G08.E09.D10.B11.A12 |
| metronidazole | 34.8823267 | 35.2669347 | -0.384608008 | 612613111306.A04 | .H01.G02.E03.D04.B05.A06 |
| decamethonium brom | 34.32476269 | 34.70962788 | -0.38486519 | 5500024030760072207033.B( | .H01.G02.E03.D04.B05.A06 |
| valproic acid | 34.0352853 | 34.42118584 | -0.385900535 | 5.50E+26 | .H01.G02.E03.D04.B05.A06 |
| pivmecillinam | 33.79033351 | 34.17633596 | -0.386002446 | 5500024030760072207033.B( | .H07.G08.E09.B11.A12 |
| valproic acid | 36.92306498 | 37.31290545 | -0.38984047 | 5.50E+32 | .G08.E09.D10.B11.A12 |
| carbinoxamine | 34.01052542 | 34.40088765 | -0.390362233 | 5500024030401071707292.D | .H07.G08.E09.D10.B11 |
| diflorasone | 34.62349094 | 35.01997127 | -0.396480325 | 5500024024213121906559.F( | .H01.G02.E03.D04.B05.A06 |
| tocainide | 34.31297453 | 34.70962788 | -0.396653354 | 5500024030760072207033.C | .H01.G02.E03.D04.B05.A06 |
| trichostatin A | 35.02689515 | 35.4281341 | -0.401238946 | 612613111306.C11 | .H07.G08.E09.D10.B11.A12 |
| ebselen | 33.99673574 | 34.40088765 | -0.404151912 | 5500024030401071707292.F | .H07.G08.E09.D10.B11 |
| ginkgolide A | 36.43321091 | 36.83996258 | -0.406751668 | 5.50E+32 | .H07.G08.E09.D10.B11.A12 |
| torasemide | 33.76889588 | 34.17633596 | -0.407440078 | 5500024030760072207033.F | .H07.G08.E09.B11.A12 |
| trifluoperazine | 32.59096302 | 33.00109826 | -0.410135242 | 5.50E+31 | .H07.G08.E09.D10.B11.A12 |
| bromperidol | 34.69770013 | 35.11129025 | -0.413590117 | 640641112706.F04 | .H01.G02.E03.D04.B05 |
| pridinol | 33.98196236 | 34.40088765 | -0.418925291 | 5500024030401071707292.F( | .H07.G08.E09.D10.B11 |
| cefotetan | 36.41761351 | 36.83996258 | -0.422349068 | 5500024024213121906563.F | .H07.G08.E09.D10.B11.A12 |
| chlorpromazine | 33.99700466 | 34.42118584 | -0.424181178 | 5500024030401071707292.F( | .H01.G02.E03.D04.B05.A06 |
| lisuride | 35.00227256 | 35.4281341 | -0.425861536 | 612613111306.A11 | .H07.G08.E09.D10.B11.A12 |
| fluvoxamine | 34.28301494 | 34.70962788 | -0.426612945 | 5500024030760072207033.F( | .H01.G02.E03.D04.B05.A06 |

| | | | | | |
|---|---|---|---|---|---|
| aconitine | 34.65720032 | 35.08636713 | -0.429166808 | 640641112706.B09 | .H07.G08.E09.D10.B11.A12 |
| pentoxifylline | 34.67524653 | 35.10850208 | -0.433255543 | 614615111406.A10 | .H07.G08.E09.D10.B11.A12 |
| oxprenolol | 36.81323908 | 37.24787327 | -0.434634199 | 5500024035736031208612.D | .H01.G02.E03.D04.B05.A06 |
| atovaquone | 34.69634873 | 35.13162255 | -0.435273827 | 5.50E+31 | .H07.G08.E09.D10.B11.A12 |
| tolfenamic acid | 34.67171275 | 35.10850208 | -0.436789324 | 614615111406.B08 | .H07.G08.E09.D10.B11.A12 |
| fenspiride | 34.67046029 | 35.10850208 | -0.438041783 | 6.15E+21 | .H07.G08.E09.D10.B11.A12 |
| nefopam | 33.02105908 | 33.45935203 | -0.438292952 | 5500024024213121906560.D | .H01.G02.E03.D04.B05.A06 |
| lasalocid | 33.69488816 | 34.13330462 | -0.438416459 | 5500024030760072207028.A | .H01.G02.D04.B05.A06 |
| antimycin A | 34.66740049 | 35.10850208 | -0.441101589 | 614615111406.F07 | .H07.G08.E09.D10.B11.A12 |
| cefuroxime | 33.51728164 | 33.96076648 | -0.44348484 | 5500024025833041107269.D | .H01.G02.E03.D04.B05.A06 |
| clotrimazole | 33.00008867 | 33.4530122 | -0.452923531 | 622623112706.H05 | .H01.G02.E03.B05.A06 |
| trichostatin A | 36.385786 | 36.83996258 | -0.454176577 | 5500024024213121906563.H | .H07.G08.E09.D10.B11.A12 |
| cefaclor | 34.67470972 | 35.13162255 | -0.456912835 | 5500024024214122006604.D | .H07.G08.E09.D10.B11.A12 |
| allantoin | 37.18005661 | 37.63864853 | -0.458591919 | 5500024024213121906564.G | .H07.G08.D10.B11.A12 |
| dl-alpha tocopherol | 36.38040487 | 36.83996258 | -0.459557708 | 5500024024213121906563.F | .H07.G08.E09.D10.B11.A12 |
| acetohexamide | 37.17336593 | 37.63864853 | -0.465282602 | 5500024024213121906564.A | .H07.G08.D10.B11.A12 |
| scopoletin | 36.98993748 | 37.46016387 | -0.470226386 | 5500024030700072107987.C | .H07.G08.E09.D10.B11.A12 |
| valproic acid | 33.94793719 | 34.42118584 | -0.473248651 | 5500024030401071707292.G | .H01.G02.E03.D04.B05.A06 |
| paclitaxel | 34.9526173 | 35.4281341 | -0.475516796 | 612613111306.A08 | .H07.G08.E09.D10.B11.A12 |
| hydroxyachillin | 34.54331938 | 35.01997127 | -0.476651892 | 5500024024213121906559.C | .H01.G02.E03.D04.B05.A06 |
| memantine | 34.22951189 | 34.70962788 | -0.480115995 | 5500024030760072207033.B | .H01.G02.E03.D04.B05.A06 |
| gramine | 34.53923521 | 35.01997127 | -0.480736059 | 5.50E+22 | .H01.G02.E03.D04.B05.A06 |
| ambroxol | 32.97592086 | 33.45797002 | -0.482049156 | 622623112706.A07 | .H07.G08.E09.D10.B11.A12 |
| chlorprothixene | 36.52375139 | 37.00786033 | -0.484108942 | 5500024024213121906563.G | .H01.G02.E03.D04.B05.A06 |
| cefalexin | 36.51940747 | 37.00786033 | -0.488452857 | 5500024024213121906563.F | .H01.G02.E03.D04.B05.A06 |
| trapidil | 36.96949359 | 37.46016387 | -0.490670279 | 5500024030700072107987.C | .H07.G08.E09.D10.B11.A12 |
| wortmannin | 33.92661721 | 34.42118584 | -0.494568627 | 5500024030401071707292.A | .H01.G02.E03.D04.B05.A06 |
| trichostatin A | 37.14399211 | 37.63864853 | -0.494656417 | 5500024024213121906564.H | .H07.G08.D10.B11.A12 |
| procainamide | 36.50519985 | 37.00786033 | -0.50266048 | 5500024024213121906563.H | .H01.G02.E03.D04.B05.A06 |
| tetracycline | 34.23448643 | 34.74789333 | -0.513406898 | 614615111406.B03 | .H01.G02.E03.D04.B05.A06 |
| LY-294002 | 36.79636007 | 37.31290545 | -0.516545382 | 5500024035736031208612.D | .G08.E09.D10.B11.A12 |
| Prestwick-860 | 33.68668442 | 34.21031976 | -0.52363534 | 5.50E+28 | .H07.G08.E09.D10.B11.A12 |
| suprofen | 33.60480655 | 34.13330462 | -0.528498063 | 5500024030760072207028.D | .H01.G02.D04.B05.A06 |
| valproic acid | 35.61074846 | 36.14209273 | -0.531344272 | 5500024017802120306174.G | .H01.G02.E03.D04.B05.A06 |
| LY-294002 | 33.88785705 | 34.42118584 | -0.533328792 | 5500024030401071707292.B | .H01.G02.E03.D04.B05.A06 |
| 15-delta prostaglandi | 33.88547671 | 34.42118584 | -0.535709127 | 5500024030401071707292.C | .H01.G02.E03.D04.B05.A06 |
| procaine | 37.08846715 | 37.63864853 | -0.550181378 | 5500024024213121906564.G | .H07.G08.D10.B11.A12 |
| phenformin | 32.90840197 | 33.45935203 | -0.550950059 | 5.50E+27 | .H01.G02.E03.D04.B05.A06 |
| mesalazine | 36.692977 | 37.24787327 | -0.554896276 | 5500024035736031208612.A | .H01.G02.E03.D04.B05.A06 |
| diprophylline | 37.07781462 | 37.63864853 | -0.56083391 | 5.50E+33 | .H07.G08.D10.B11.A12 |
| laudanosine | 34.5467555 | 35.11129025 | -0.564534752 | 640641112706.C06 | .H01.G02.E03.D04.B05 |
| Trolox C | 34.54310794 | 35.11129025 | -0.568182313 | 640641112706.D05 | .H01.G02.E03.D04.B05 |
| lansoprazole | 33.60468108 | 34.17633596 | -0.571654879 | 5500024030760072207033.C | .H07.G08.E09.B11.A12 |
| xylometazoline | 34.52454533 | 35.10850208 | -0.583956747 | 6.15E+22 | .H07.G08.E09.D10.B11.A12 |
| troglitazone | 33.82891193 | 34.42118584 | -0.592273904 | 5500024030401071707292.C | .H01.G02.E03.D04.B05.A06 |
| baclofen | 34.82809146 | 35.4281341 | -0.600042637 | 612613111306.C12 | .H07.G08.E09.D10.B11.A12 |
| sotalol | 34.10250401 | 34.70962788 | -0.60712387 | 5.50E+23 | .H01.G02.E03.D04.B05.A06 |
| risperidone | 33.56792066 | 34.17633596 | -0.608415299 | 5500024030760072207033.G | .H07.G08.E09.B11.A12 |
| trimethobenzamide | 34.65367669 | 35.2669347 | -0.613258011 | 612613111306.A03 | .H01.G02.E03.D04.B05.A06 |
| norcyclobenzaprine | 34.51078749 | 35.13162255 | -0.620835062 | 5500024024214122006604.G | .H07.G08.E09.D10.B11.A12 |
| cinoxacin | 33.77856269 | 34.40088765 | -0.622324954 | 5.50E+32 | .H07.G08.E09.D10.B11 |
| oxamniquine | 34.07257057 | 34.70962788 | -0.637057315 | 5500024030760072207033.D | .H01.G02.E03.D04.B05.A06 |
| sulindac | 36.99721672 | 37.63864853 | -0.641431806 | 5500024024213121906564.D | .H07.G08.D10.B11.A12 |
| securinine | 33.75354916 | 34.40088765 | -0.647338487 | 5500024030401071707292.C | .H07.G08.E09.D10.B11 |
| acepromazine | 34.43444908 | 35.08636713 | -0.651918046 | 640641112706.C08 | .H07.G08.E09.D10.B11.A12 |
| harmine | 34.42943532 | 35.08636713 | -0.656931809 | 640641112706.G10 | .H07.G08.E09.D10.B11.A12 |
| ethosuximide | 34.45075606 | 35.10850208 | -0.657746015 | 614615111406.C10 | .H07.G08.E09.D10.B11.A12 |
| chlorphenesin | 34.44596803 | 35.10850208 | -0.662534049 | 614615111406.C09 | .H07.G08.E09.D10.B11.A12 |
| LY-294002 | 36.64945726 | 37.31290545 | -0.663448195 | 5500024035736031208612.B | .G08.E09.D10.B11.A12 |
| protriptyline | 36.79527889 | 37.46016387 | -0.664884983 | 5500024030700072107987.F | .H07.G08.E09.D10.B11.A12 |
| valproic acid | 35.46842782 | 36.14209273 | -0.673664911 | 5500024017802120306174.A | .H01.G02.E03.D04.B05.A06 |
| flupentixol | 36.33403983 | 37.00786033 | -0.673820502 | 5500024024213121906563.D | .H01.G02.E03.D04.B05.A06 |
| captopril | 34.58962303 | 35.2669347 | -0.677311679 | 612613111306.D06 | .H01.G02.E03.D04.B05.A06 |
| trichostatin A | 34.05545584 | 34.74789333 | -0.692437487 | 614615111406.A02 | .H01.G02.E03.D04.B05.A06 |
| harmol | 34.41753298 | 35.11129025 | -0.693757266 | 640641112706.A05 | .H01.G02.E03.D04.B05 |
| paracetamol | 33.51340966 | 34.21031976 | -0.696910099 | 5500024030760072207028.H | .H07.G08.E09.D10.B11.A12 |
| 15-delta prostaglandi | 35.43906865 | 36.14209273 | -0.703024077 | 5500024017802120306174.C | .H01.G02.E03.D04.B05.A06 |
| valproic acid | 35.43804264 | 36.14209273 | -0.704050084 | 5500024017802120306174.F | .H01.G02.E03.D04.B05.A06 |
| mefenamic acid | 36.93424583 | 37.63864853 | -0.704402703 | 5500024024213121906564.C | .H07.G08.D10.B11.A12 |

| | | | | | |
|---|---|---|---|---|---|
| trichostatin A | 36.53196366 | 37.24471367 | -0.712750013 | 5500024030700072107987.F( | .H01.G02.E03.D04.B05.A06 |
| ketoconazole | 36.29077157 | 37.00786033 | -0.71708876 | 5500024024213121906563.D | .H01.G02.E03.D04.B05.A06 |
| apramycin | 33.96989456 | 34.70962788 | -0.739733323 | 5500024030760072207033.F( | .H01.G02.E03.D04.B05.A06 |
| amiodarone | 34.3991054 | 35.15552717 | -0.756421771 | 5500024024214122006604.F( | .H01.G02.E03.D04.A06 |
| valproic acid | 36.52329317 | 37.31290545 | -0.789612283 | 5500024035736031208612.F( | .G08.E09.D10.B11.A12 |
| LY-294002 | 33.62724215 | 34.42118584 | -0.793943682 | 5500024030401071707292.D | .H01.G02.E03.D04.B05.A06 |
| azathioprine | 34.60302041 | 35.4281341 | -0.825113691 | 612613111306.D09 | .H07.G08.E09.D10.B11.A12 |
| moroxydine | 34.59662545 | 35.4281341 | -0.831508647 | 612613111306.D08 | .H07.G08.E09.D10.B11.A12 |
| ellipticine | 34.25374815 | 35.08636713 | -0.832618974 | 640641112706.F12 | .H07.G08.E09.D10.B11.A12 |
| thiostrepton | 34.29535691 | 35.13162255 | -0.836265649 | 5500024024214122006604.H | .H07.G08.E09.D10.B11.A12 |
| sulfaphenazole | 36.79940102 | 37.63864853 | -0.839247512 | 5500024024213121906564.H | .H07.G08.D10.B11.A12 |
| pentetrazol | 34.24705969 | 35.10850208 | -0.861442385 | 614615111406.H12 | .H07.G08.E09.D10.B11.A12 |
| 6-benzylaminopurine | 32.59311787 | 33.45935203 | -0.866234155 | 5500024024213121906560.D | .H01.G02.E03.D04.B05.A06 |
| N-acetyl-L-leucine | 36.37466345 | 37.24471367 | -0.870050224 | 5.50E+27 | .H01.G02.E03.D04.B05.A06 |
| oxolinic acid | 34.22589914 | 35.10850208 | -0.882602933 | 614615111406.F12 | .H07.G08.E09.D10.B11.A12 |
| valproic acid | 36.3909057 | 37.31290545 | -0.921999757 | 5500024035736031208612.A( | .G08.E09.D10.B11.A12 |
| dequalinium chloride | 36.04453234 | 37.00786033 | -0.963327991 | 5500024024213121906563.F( | .H01.G02.E03.D04.B05.A06 |
| rosiglitazone | 35.1615202 | 36.12983309 | -0.968312888 | EC2004032003AA | EC2004032002AA |
| etamivan | 34.14048032 | 35.11129025 | -0.970809931 | 6.41E+17 | .H01.G02.E03.D04.B05 |
| trichostatin A | 33.44787813 | 34.42118584 | -0.973307707 | 5500024030401071707292.C | .H01.G02.E03.D04.B05.A06 |
| morantel | 36.66372349 | 37.63864853 | -0.97492504 | 5500024024213121906564.G | .H07.G08.D10.B11.A12 |
| todralazine | 36.65041631 | 37.63864853 | -0.988232219 | 5500024024213121906564.G | .H07.G08.D10.B11.A12 |
| tolnaftate | 34.26373984 | 35.2669347 | -1.003194866 | 612613111306.A02 | .H01.G02.E03.D04.B05.A06 |
| sodium phenylbutyrat | 35.6465771 | 36.67804214 | -1.031465036 | EC2004042305AA | EC2004042302AA |
| chrysin | 36.42113217 | 37.46016387 | -1.039031699 | 5500024030700072107987.H | .H07.G08.E09.D10.B11.A12 |
| hyoscyamine | 34.03465862 | 35.10850208 | -1.073843455 | 6.15E+23 | .H07.G08.E09.D10.B11.A12 |
| troglitazone | 35.03613777 | 36.12983309 | -1.093695327 | EC2004032004AA | EC2004032002AA |
| verteporfin | 36.15210901 | 37.24787327 | -1.095764266 | 5500024035736031208612.F( | .H01.G02.E03.D04.B05.A06 |
| propafenone | 33.99580021 | 35.11129025 | -1.115490042 | 640641112706.F03 | .H01.G02.E03.D04.B05 |
| nifedipine | 36.44715009 | 37.63864853 | -1.191498437 | 5500024024213121906564.D | .H07.G08.D10.B11.A12 |
| solanine | 33.75689601 | 35.01997127 | -1.263075263 | 5500024024213121906559.D | .H01.G02.E03.D04.B05.A06 |
| parthenolide | 33.80511904 | 35.11129025 | -1.306171207 | 640641112706.C01 | .H01.G02.E03.D04.B05 |
| vorinostat | 33.09407401 | 34.42118584 | -1.327111825 | 5.50E+23 | .H01.G02.E03.D04.B05.A06 |
| apigenin | 35.5107929 | 36.83996258 | -1.329169676 | 5.50E+28 | .H07.G08.E09.D10.B11.A12 |
| rofecoxib | 34.74809467 | 36.12983309 | -1.381738421 | EC2004032005AA | EC2004032002AA |
| estradiol | 34.78813617 | 36.19585457 | -1.4077184 | EC2004032010AA | EC2004032009AA |
| wortmannin | 34.70389207 | 36.19585457 | -1.491962493 | EC2004032012AA | EC2004032009AA |
| tetraethylenepentamii | 35.09089966 | 36.67804214 | -1.587142485 | EC2004042306AA | EC2004042302AA |
| raloxifene | 34.58157554 | 36.19585457 | -1.61427903 | EC2004032011AA | EC2004032009AA |
| clopamide | 31.71959007 | 33.45797002 | -1.738379944 | 6.23E+22 | .H07.G08.E09.D10.B11.A12 |
| dirithromycin | 33.28705768 | 35.11129025 | -1.824232574 | 640641112706.H04 | .H01.G02.E03.D04.B05 |
| chlortetracycline | 33.4197275 | 35.4281341 | -2.008406601 | 612613111306.A07 | .H07.G08.E09.D10.B11.A12 |
| ajmaline | 32.97105775 | 35.11129025 | -2.140232496 | 640641112706.A04 | .H01.G02.E03.D04.B05 |
| trichostatin A | 33.97743969 | 36.14209273 | -2.164653037 | 5500024017802120306174.C | .H01.G02.E03.D04.B05.A06 |
| azacyclonol | 33.22679593 | 35.4281341 | -2.201338173 | 612613111306.F12 | .H07.G08.E09.D10.B11.A12 |
| lobeline | 32.87491587 | 35.08636713 | -2.211451256 | 6.41E+23 | .H07.G08.E09.D10.B11.A12 |
| vorinostat | 33.91319192 | 36.14209273 | -2.228900814 | 5.50E+23 | .H01.G02.E03.D04.B05.A06 |
| glibenclamide | 31.08342575 | 33.4530122 | -2.369586453 | 622623112706.H02 | .H01.G02.E03.B05.A06 |
| clindamycin | 32.36273132 | 34.74789333 | -2.385162005 | 614615111406.F01 | .H01.G02.E03.D04.B05.A06 |
| chlorzoxazone | 32.72003682 | 35.10850208 | -2.388465254 | 614615111406.F09 | .H07.G08.E09.D10.B11.A12 |
| valproic acid | 34.27194979 | 36.67804214 | -2.406092354 | EC2004042303AA | EC2004042302AA |
| ampyrone | 32.33060324 | 34.74789333 | -2.41729009 | 614615111406.A04 | .H01.G02.E03.D04.B05.A06 |
| trichostatin A | 34.53995848 | 37.31290545 | -2.772946968 | 5500024035736031208612.C | .G08.E09.D10.B11.A12 |
| piperlongumine | 32.25981733 | 35.08636713 | -2.8265498 | 640641112706.F11 | .H07.G08.E09.D10.B11.A12 |
| danazol | 32.54635143 | 35.4281341 | -2.881782672 | 612613111306.B08 | .H07.G08.E09.D10.B11.A12 |
| colforsin | 32.06885637 | 35.03613777 | -2.9672814 | EC2005081605AA | EC2005081602AA |
| valproic acid | 33.10374681 | 36.67804214 | -3.574295328 | EC2004042304AA | EC2004042302AA |
| estradiol | 31.08260455 | 35.03613777 | -3.953533213 | EC2005081604AA | EC2005081602AA |
| vorinostat | 33.34232906 | 37.31290545 | -3.970576396 | 5.50E+29 | .G08.E09.D10.B11.A12 |
